# Topological supramolecular network enabled highly conductive and stretchable organic bioelectronics

**DOI:** 10.1101/2022.01.16.476423

**Authors:** Yuanwen Jiang, Zhitao Zhang, Yi-Xuan Wang, Deling Li, Charles-Théophile Coen, Ernie Hwaun, Gan Chen, Hung-Chin Wu, Donglai Zhong, Simiao Niu, Weichen Wang, Aref Saberi, Jian-Cheng Lai, Yang Wang, Artem A. Trotsyuk, Kang Yong Loh, Chien-Chung Shih, Wenhui Xu, Kui Liang, Kailiang Zhang, Wenping Hu, Wang Jia, Zhen Cheng, Reinhold H. Dauskardt, Geoffrey C. Gurtner, Jeffery B.-H. Tok, Karl Deisseroth, Ivan Soltesz, Zhenan Bao

**Affiliations:** Department of Chemical Engineering, Stanford University, Stanford, CA 94305, USA; Tianjin Key Laboratory of Molecular Optoelectronic Sciences, Department of Chemistry, School of Science, Tianjin University, Tianjin, 300072, China; Department of Radiology, Molecular Imaging Program at Stanford (MIPS), Stanford University, Stanford, CA 94305, USA; Department of Neurosurgery, Beijing Tiantan Hospital, Capital Medical University, Beijing, 100070, China; Department of Neurosurgery, Stanford University, Stanford, CA 94305, USA; Department of Materials Science and Engineering, Stanford University, Stanford, CA 94305, USA; State Key Laboratory of Coordination Chemistry, School of Chemistry and Chemical Engineering, Nanjing University, Nanjing 210093, China; Department of Bioengineering, Stanford University, CA 94305, USA; Department of Surgery, Stanford University, Stanford, CA 94305, USA; Department of Chemistry, Stanford Chemistry, Engineering & Medicine for Human Health (ChEM-H), Stanford University, Stanford, CA 94305, USA; BOE Technology Center, BOE Technology Group Co., Ltd, Beijing, 100176, China; Department of Psychiatry and Behavioral Sciences, Stanford University, Stanford, CA 94305, USA; Howard Hughes Medical Institute, Stanford University, Stanford, CA 94305, USA

## Abstract

Intrinsically stretchable bioelectronic devices based on soft and conducting organic materials have been regarded as the ideal interface for seamless and biocompatible integration with the human body. However, the grand challenge remains for the conducting polymer to possess both high mechanical ductility and good electrical conduction at cellular level feature sizes. This longstanding material limitation in organic bioelectronics has impeded the full exploitation of its unique benefits. Here, we introduce a new molecular engineering strategy based on rationally designed topological supramolecular networks, which allows effective decoupling of competing effects from multiple molecular building blocks to meet complex requirements. We achieve two orders of magnitude improvement in the conductivity under 100% strain in physiological environment, along with the capability for direct photopatterning down to 2 μm. These unprecedented capabilities allow us to realize previously inaccessible bioelectronic applications including high-resolution monitoring of ‘soft and malleable’ creatures, e.g., octopus, and localized neuromodulation down to single nucleus precision for controlling organ-specific activities through delicate tissues, e.g., brainstem.

---

Implantable and wearable bioelectronic systems are essential in numerous biomedical applications, ranging from multimodal monitoring of physiological signals for disease diagnosis(*1-6*), programmable modulation of neural or cardiac activities for therapeutics(*7-12*), restoration of lost sensorimotor functions for prosthetics(*13-17*), and even augmented reality(*18*). However, many of these existing devices experienced performance degradation, and even failure, when operating in a dynamic-moving tissue environment(*19*). This stems primarily from the mechanical mismatches (e.g., modulus and stretchability), which inevitably lead to either interfacial delamination, fibrotic encapsulation or gradual tissue scarring(*20*).

To maintain effective electrical signal exchanges across electrode-bio interfaces, efforts have been made to render rigid electronics and inorganic materials compliant to soft biological tissues(*21, 22*). Meanwhile, intrinsically stretchable organic electronics is fast-emerging as a promising candidate with several unique advantages(*23*). First, they do not suffer from the inherent trade-off between overall system stretchability and device density found for rigid materials. Therefore, high resolution mapping/interrogation can be realized for conformal biointerfaces(*24, 25*) (**Fig. 1A**). Second, the unique electronic-ionic dual conduction mechanism of conducting polymers (CPs) reduces the electrode-tissue interfacial impedance, thus allowing high recording fidelity and efficient stimulation charge injection(*26-28*). However, electrical conductivities of existing stretchable CPs are too low after being micro-fabricated into bioelectronic devices(*23*). As a result, rigid metal interconnects are still required, which greatly diminished the advantages of soft CPs(*3*). Even with stretchable microcracked-metal structures, they are limited to certain thickness and width to maintain modest stretchability without severe degradation of electrical conduction(*29*). These restrictions usually result in high sheet resistance and large feature sizes of >50 μm(*30*).

**Fig. 1.**
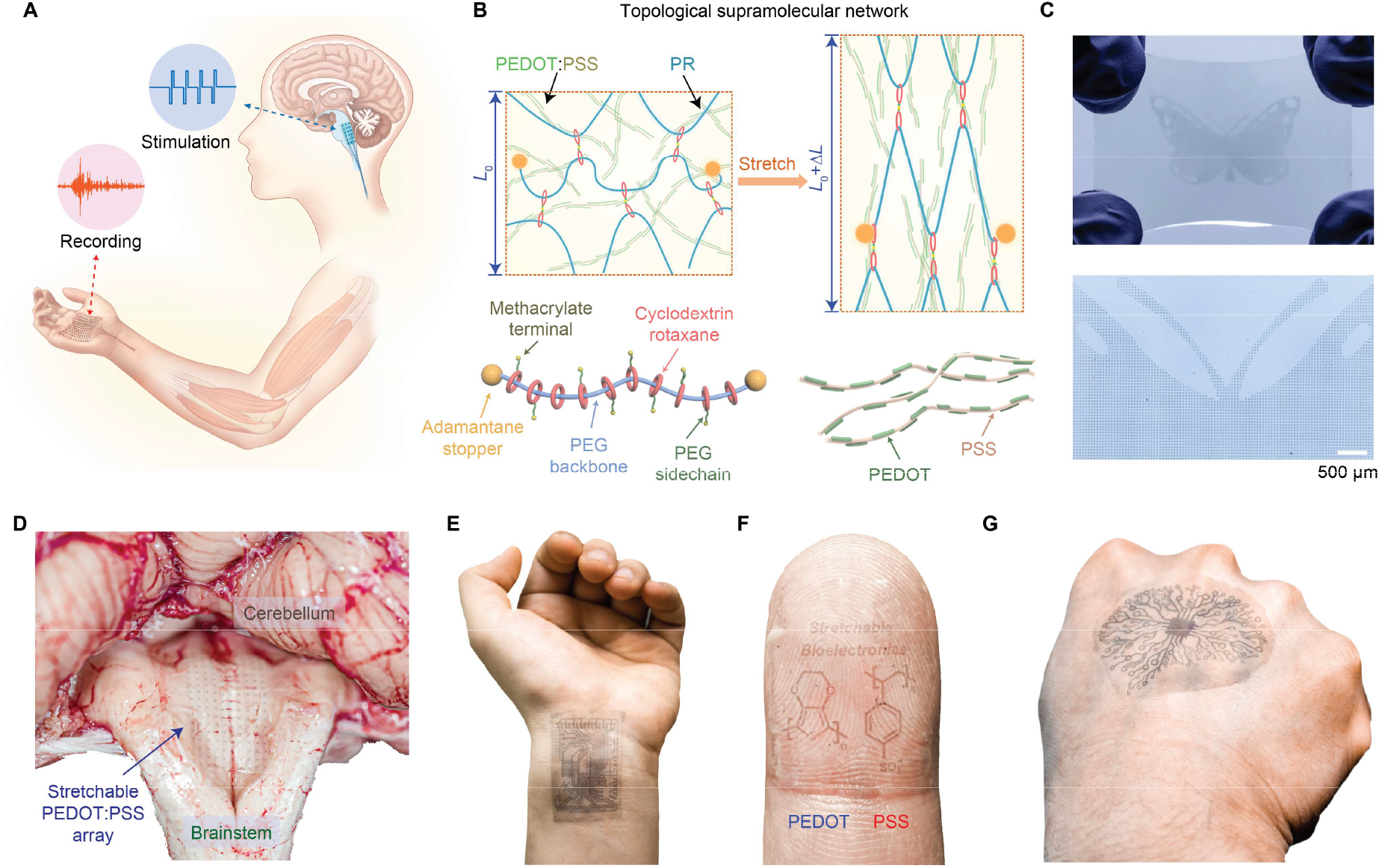
Intrinsically stretchable organic electronics for multimodal and conformal biointerfaces. **(A)** Stretchable multielectrode array can form seamless interfaces with multiple organs for bidirectional interrogations with high precision. **(B)** Schematic diagram illustrating the intrinsically stretchable topological supramolecular network with the key molecular building blocks of polyrotaxane monomers. **(C)** The topological supramolecular network allows direct photopatterning of large area high density stretchable electrode array. **(D-G)** Photographic images showing the conformal interface between stretchable PEDOT:PSS devices and underlying tissues, including brainstem (**D**), wrist (**E**), finger (**F**), and the back of hand (**G**).

To address this longstanding bottleneck, we describe here a rationally designed topological supramolecular network to simultaneously enable three important advancements in bioelectronics: (i) biocompatible and stretchable CPs with record-breaking conductivity (∼2700 S cm^-1^ at 0% strain; ∼6000 S cm^-1^ at 100% strain), (ii) direct photo-patternability down to 2 μm feature sizes, and (iii) high stretchability maintained after micro-fabrication with no crack formation under at least 100% strain (**Fig. 1B-C**), which are essential for low-impedance and seamless biointegration. All previously reported stretchable CPs in physiological environment showed substantial cracks after ∼25% strain with poor conductivities of <10 S cm^-1^ at 100% strain. Our key hypothesis is that incorporating topology into the molecular design may decouple multiple competing effects to meet complex requirements(*31*). We chose ‘mechanically-interlocked’ structures because they possess large conformational freedom from the mobile junctions(*32-35*). Therefore, using a single supramolecular crosslinker, we could simultaneously enable intrinsic stretchability, high conductivity, and direct photopatternability for CPs (**Fig. 1B**). These unprecedented properties allowed us to demonstrate new capabilities in bioelectronic applications, including: (i) cellular resolution recording of muscle signals from human palm as well as highly malleable and soft-bodied octopus, and (ii) localized neuromodulation through delicate brainstem down to single nucleus level for precise controls of individual muscle groups at tongue, whisker, and neck (**Fig. 1D-G**).

Poly(3,4-ethylenedioxythiophene):polystyrene sulfonate (PEDOT:PSS) is among the highest performing and most investigated CPs for bioelectronic devices(*3, 23, 27*). It is an aqueous suspension consisting of colloidal particles with PEDOT-rich cores and PSS-rich shells. As a result, pristine PEDOT:PSS films have low conductivity (<10 S cm^-1^) and cracks at <10% strain due to the sparse PEDOT conducting connections between particles(*26, 36*). Although ionic and molecular additives can improve both conductivity and stretchability(*37*), after solvent treatment during device fabrication and immersion in aqueous biological environment, performance of existing PEDOT:PSS films dropped substantially (initial conductivity of <50 S cm^-1^; crack onset strain of <20%)(*38*) because the non-crosslinked additives were washed away and lost their effects. To achieve high conductivity and stretchability with stable operation in biological environment, we designed a crosslinkable supramolecular additive, based on a polyrotaxane (PR) structure. Specifically, it is comprised of a polyethylene glycol (PEG) backbone and sliding cyclodextrins (CDs) functionalized with PEG methacrylate (PEGMA) sidechains, to induce high conductivity, stretchability, and photo-patternability (**Fig. 2A-B**, synthesis and characterizations in **figs. S1-9**). While PEG was known to induce aggregation of PEDOT to enhance conductivity(*39*), its strong tendency to crystalize would lead to phase separation and poor stretchability. In our design, we hypothesized that the incorporation of sliding CD units may prevent the crystallization of PEG to provide a better stretchability. Thus, our single additive molecular design with PR-PEGMA should fulfill all our set forth requirements, while containing a minimal amount of inert components, and we termed the topological network enabled stretchable conducting polymer bioelectronics as ‘TopoE’.

**Fig. 2.**
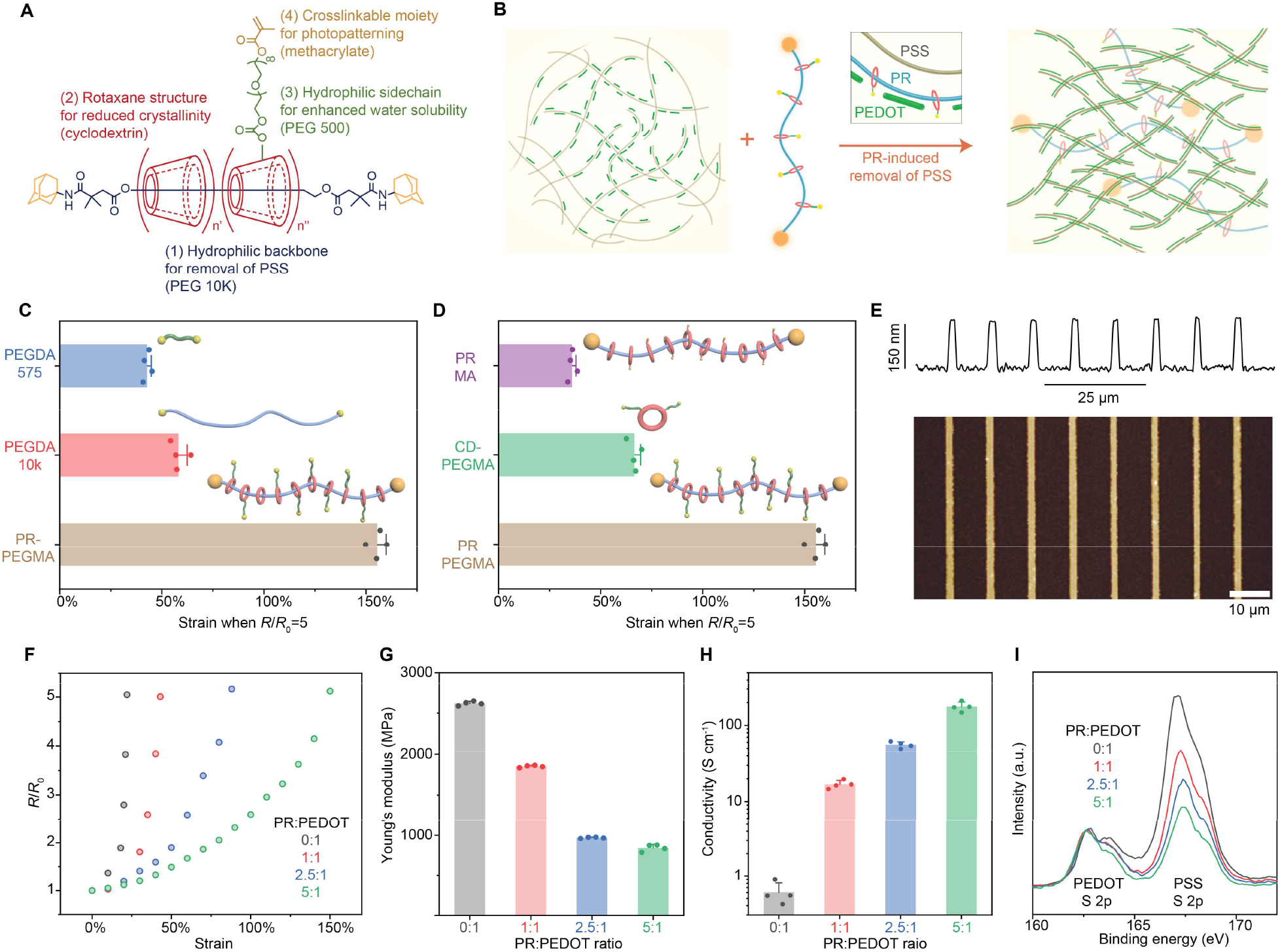
Polyrotaxane-based topological network enables simultaneously enhanced conductivity, stretchability, and photopatternability of PEDOT:PSS. **(A)** Chemical structure of polyrotaxane polyethylene glycol methacrylate (PR-PEGMA) and individual roles of each building block. **(B)** Schematic diagram illustrating the interaction between PR and PEDOT:PSS for enhanced conductivity. **(C-D)** Stretching tests showing that PR blended PEDOT:PSS could substantially enhance stretchability compared to other control samples. **(E)** AFM height image and corresponding surface profile of a photopatterned TopoE array with 2-μm width. **(F)** Resistance change as a function of strain for TopoE films with different PR over PEDOT:PSS dry mass weight ratios. All films in **F-I** were UV crosslinked after blending followed by washing in water and blow drying. **(G)** Statistical comparison of Young’s moduli measured by nanoindentation for different TopoE films indicating that PR can reduce the overall film stiffness. **(H)** Four-point probe measurements showing enhanced film conductivity with higher PR content. **(I)** XPS profiles indicating reduced PSS content as PR:PEDOT ratio increases in the film.

To validate our hypothesis, we first investigated samples with only the PEG diacrylate (PEGDA) backbone (molecular weight 10k, PEGDA-10k) or sidechain PEGDA (molecular weight 575, PEGDA-575) blended into PEDOT:PSS for crosslinking (see **Methods** in Supporting Information). In both cases, only a slight improvement in stretchability was observed, i.e., from ∼5% to ∼50% or 30% strain, respectively (**Fig. 2C, fig. S11**). Further analyses using atomic force microscopy (AFM) and grazing incident X-ray diffraction spectroscopy (GIXD) revealed that the films were inhomogeneous and suffered severe microphase separations, most likely due to the crystallization of PEG (**figs. S12-13**). For PEGylated cyclodextrin (CD-PEGMA), its poor solubility in water prevented uniform blending with PEDOT:PSS, such that the final crack onset strain was measured to be only ∼30%. Similarly, for PR-MA without any PEG sidechains, it was also observed that it could not dissolve well in water to enable good blending with PEDOT:PSS (**Fig. 2D, fig. S11**).

However, PR-PEGMA was found to have the balance between molecular topology and chemical polarity. It has good solubility in water due to the PEG-based backbone and sidechains, yet low crystallinity due to the bulky CD rings (**fig. S13**). As a result, the photo-crosslinked TopoE film with PR-PEGMA/PEDOT:PSS was uniform and can be stretched up to 150% strain. The number of sidechains on each PR-PEGMA molecule was also found to be important, with eight PEGMA chains being the optimal condition with the best stretchability (**fig. S14**). Finally, we investigated other common multi-arm PEGDA derivatives and observed that only the pulley-topology with CD was effective in enhancing the stretchability of PEDOT:PSS (**fig. S15**). Next, we observed that the TopoE film can be directly photopatterned down to 2 μm (**Fig. 2E, fig. S16**), while the crosslinked film after development could still maintain stretchability (**fig. S17**). With these developments, we chose PR-PEGMA_8_ (abbreviated as PR) for all future characterizations and fabrications.

As the PR content increased, higher stretchability (**Fig. 2F, fig. S18**) and lower Young’s modulus were obtained (**Fig. 2G**). Importantly, a higher PR content also led to a better PEDOT:PSS conductivity with a significant enhancement of over two orders of magnitude compared to pristine PEDOT:PSS (**Fig. 2H**). X-ray photoelectron spectroscopy (XPS) revealed a decreasing trend of PSS over PEDOT ratio as the PR-PEGMA content increased, indicating a significant portion of insulating PSS was removed after blending and water development (**Fig. 2I**). UV-vis and Raman spectroscopy also confirmed that less PSS remained in the TopoE film (**fig. S19**). With PR, its polar PEG chains replaced a portion of PSS in interacting with PEDOT, which led to the enhanced aggregation of PEDOT(*39*) (**Fig. 2B, fig. S20**). In addition to increased PEDOT content, AFM phase images of the blended films further revealed that the microscale morphology of PEDOT gradually evolved from grain-like particles to percolated micro-webs, which was favored for enhanced charge transport, especially under strain (**figs. S21-22**).

Next, we aim to further improve the conductivity of TopoE. We hypothesized that conductivity of TopoE can be enhanced by sulfuric acid treatment, while still maintaining a high stretchability with the crosslinked network, which we termed as ‘TopoE-S’ (**Fig. 3A**). We first confirmed that the topological network could survive the acid-dipping process using XPS depth profiling investigation (**fig. S23**). We also used GIXD to track the change of PEDOT crystallization (**figs. 3b-c**). The neat TopoE film showed a scattering profile with a weak (100) diffraction peak, corresponding to the PEDOT lamellar packing. After acid treatment, the TopoE-S film showed high-order diffraction peaks with strong intensities while the (100) peak shifted to a larger *q* value (0.49 vs 0.46 Å^-1^), suggesting denser lamellar packing and longer range order(*40*). In addition, the π-π (020) peak also emerged in the in-plane direction, indicating longer range order for π-π stackings between PEDOTs(*41*). Besides the increased PEDOT interactions, AFM phase images showed that the microscopic morphology of PEDOT changed to interconnected fibers after acid treatment (**Fig. 3D**). With both these changes, an one order of magnitude enhancement of the film conductivity than TopoE and three orders of magnitude enhancement than pristine PEDOT:PSS film up to ∼2700 S cm^-1^ was obtained for TopoE-S (**Fig. 3E**). We note that the TopoE-S film was also optically transparent in the visible range with a transmittance of 94% at 550 nm and a sheet resistance as low as 69 Ω sq^-1^. These values for stretchable TopoE-S are comparable to state-of-art non-stretchable PEDOT:PSS(*40*), carbon nanotube(*42*), silver nanowire films(*43*), and even widely used indium tin oxide (ITO)(*44*) based transparent conductor (**fig. S24**).

**Fig. 3.**
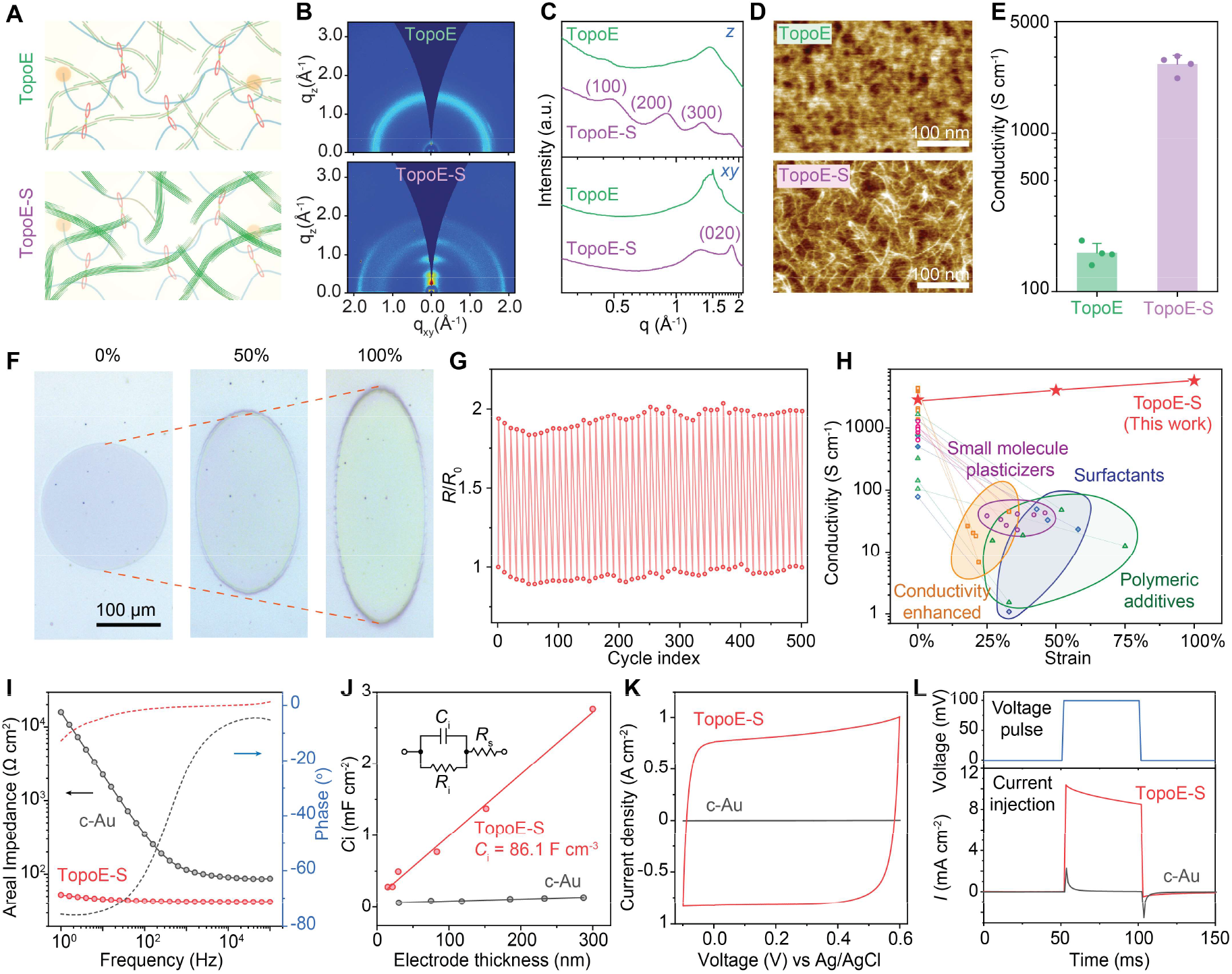
Fully-crosslinked topological network can afford post-treatment towards record high conductivity and stretchability. **(A)** Schematic diagram illustrating the change of crystallinity by acid treatment. **(B-C)** 2D spectra (**B**) and 1D profile (**C**) of GIXD collected from TopoE and TopoE-S. **(D)** AFM phase images showing the morphological changes of TopoE and TopoE-S. **(E)** Electrical measurement showing enhanced conductivity after acid treatment for TopoE-S. **(F)** Optical microscope images showing the shape evolution of a TopoE-S pattern during stretching without inducing any cracks. **(G)** Cyclic stretching test between 0% to 100% strain showing reversible resistance changes of TopoE-S. **(H)** Conductivity over strain plots showing high conductivity under strain for TopoE-S versus literature reported PEDOT:PSS after water soaking. All previously reported methods suffer from severe conductivity degradation from soaking and/or from strain (detailed data and references in fig. S29). **(I)** EIS measurements showing the reduced impedance of TopoE-S compared to stretchable cracked Au (c-Au). **(J)** Interfacial capacitance from TopoE-S and Au as a function of electrode volume. **(K)** CV sans showing the enhanced charge storage capacity of TopoE-S over c-Au. **(L)** Current measurements following transient voltage pulses showing better charge injection of TopoE-S than c-Au.

The highly conductive TopoE-S film can be directly photo-patterned in one step while previous PEDOT:PSS patterns had to go through multi-step lithography using photoresist together with metal-protected etching process(*38*). The photo-patterned circular structure could be stretched to an elliptical shape at 100% strain, while remaining intact without observable cracks (**Fig. 3F, figs. S25-26**). AFM phase images and corresponding fast Fourier transform (FFT) spectrograms of the TopoE-S film under strain showed that the PEDOT micro-fibers became aligned along the stretching direction and return to isotropic orientations when strain was released (**fig. S27**). GIXD spectra confirmed the retention of the PEDOT crystallization in directions both parallel and perpendicular to the strained axis (**fig. S28**). Finally, the cyclic stretching testing between 0-100% strain for the TopoE-S film showed reversibly conductivity for at least 500 cycles (**Fig. 3G**). For bioelectronics application, the PEDOT:PSS electrode needs to be immersed in physiological environment to remove any soluble or toxic additives. After such an essential treatment, previously reported PEDOT:PSS systems with conductivity-enhancement treatment(*40*), small molecule plasticizers(*37*), surfactant blending(*45*) or polymeric additives(*46*) all suffered from substantial drops of conductivities upon stretching although the polymeric additives were slightly better than the small molecule ones (**Fig. 3H, fig. S29**). In sharp contrast, our TopoE-S was able to maintain its high performance at ∼2700 S cm^-1^ initial conductivity and ∼6000 S cm^-1^ at 100% strain with chain alignments (**Fig. 3H, figs. S27 & S29**).

Directly related to bioelectronic applications, electrochemical impedance of TopoE-S in phosphate buffered saline (PBS) solution is lower than stretchable cracked Au (c-Au) in all frequency ranges (**Fig. 3I**). c-Au electrode is among the best stretchable electrodes for bioelectronics (*47*). Its high-frequency unit-area impedance is determined by thickness and feature size, which are constrained by crack size. Its low-frequency unit-area impedance is several orders of magnitude higher than TopoE-S because the effective interfacial capacitance for TopoE-S is much larger than that of c-Au (**Fig. 3J**)(*41*). Such a large interfacial capacitance is also responsible for high charge storage capacity (CSC) (**Fig. 3k**) and efficient charge injection (**Fig. 3L**), which are both important for low-voltage electrical stimulation(*38*). The high electrical conductivity is essential for TopoE-S to outperform c-Au at all frequency ranges (**fig. S30**). Otherwise, if the electrical conductivity of PEDOT is lower than ∼300 S cm^-1^, the reduced impedance of PEDOT versus c-Au is only expected in the low frequency regime. The impedance value of the TopoE-S electrode remained stable in PBS for at least one month (**fig. S31**). When the TopoE-S electrode was at 100% strain, its impedance only showed slight increase due to the electrode geometry change without inducing any cracks (**fig. S32**). The above obtained results suggested that TopoE-S will be ideal for both electrophysiological recording and electrical stimulation.

Next, we developed a fabrication process for high-density stretchable electrode arrays by optimizing the chemical orthogonality and surface energy of each elastomeric layers (**Figs. 4A-B, figs. S10 & S33**). The as-fabricated stretchable TopoE-S array has a narrow distribution of both the impedance and the phase angle (**fig. S34**). Since the entire electrode array (as thin as 20 μm) is made of soft and elastic materials, it can be easily attached onto human skin with great conformability (**Fig. 4C**). As a result, it allowed high-density surface electromyography (sEMG) recording even on moving muscle. Previously, typical inter-electrode distance (IED) for sEMG devices is usually ∼5 mm or larger because the high impedance of rigid electrodes and poor skin contact do not allow reliable recording with high density(*48*). Here, because of the low impedance of PEDOT:PSS and low modulus of the entire electrode array, we successfully reduced the IED down to 500 μm with electrode width of 100 μm, while still being able to capture high fidelity sEMG signals (**Fig. 4B**). In addition, the low unit-area impedance enabled our stretchable electrode array could resolve sEMG propagation dynamics using small electrode area with unprecedented spatial resolution (**Figs. 4D-E**). Besides recording dynamic information, the integrated EMG signals over time was also distinct enough to differentiate other hand gestures in a highly reproducible manner (**figs. S35-40**). Further analysis of these subtle muscle activities, especially on the cellular scale, may offer critical insights to the study of kinesiology and serve as the foundation for future skin electronics as a non-invasive and robust human-machine interface for prosthetics applications.

**Fig. 4.**
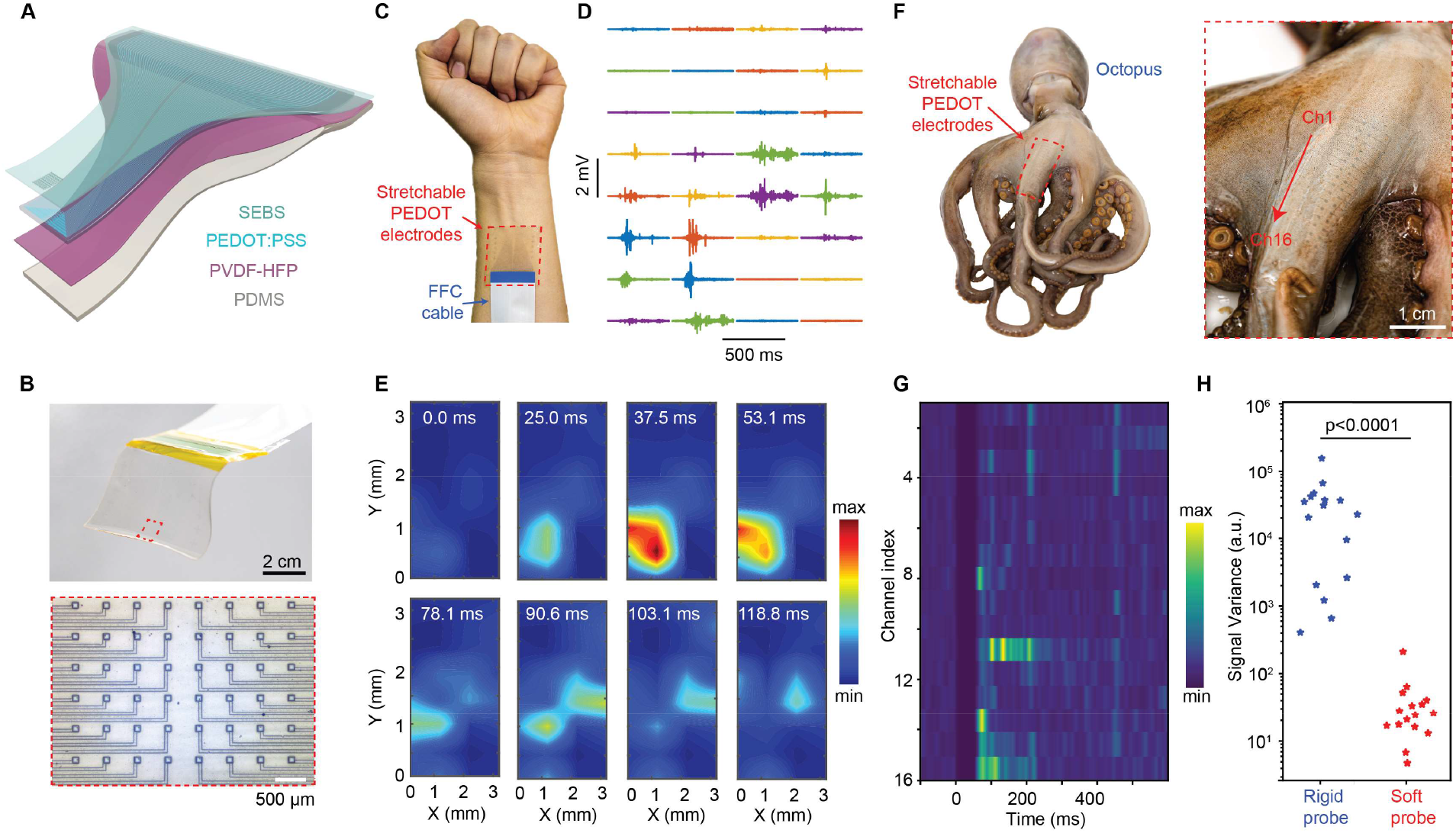
Intrinsically stretchable electrode array allows high-resolution electrophysiological monitoring of deformable tissues. **(A)** Exploded view of the multilayered stacks for the stretchable electrode array. **(B)** Photograph of an as-fabricated device (upper row) and an optical microscope image of the active site (lower row). **(C)** Photographic image of the sEMG measurement setup including a tissue conforming stretchable electrode array and a flexible flat cable (FFC) for input/output communications. **(D)** Representative EMG recording traces during a fist gesture with distinct spatiotemporal dynamics of individual channels. **(E)** Temporal evolution of EMG activity across different channels corresponding to the raw traces in (**D**) while fisting. **(F)** Photographic images of a stretchable electrode array attached to an octopus arm. **(G)** Peristimulus time histogram of evoked EMG activities of the octopus during muscle twitching. **(H)** Comparison of the signal variance during resting state recorded by a rigid probe versus a soft and stretchable probe.

Another unique new capability enabled by the stretchable electrode array is to measure sEMG signals from soft-bodied creatures, such as an octopus, whose muscles can undergo much larger deformations than human’s (**Fig. 4F**). We observed that upon electrical stimulation of the octopus arm, our stretchable sEMG array could consistently record the muscle activity dynamics with good signal-to-noise ratio (SNR) (**Fig. 4G, figs. S41-42**). On the contrary, a rigid probe made of PEDOT:PSS on polyimide was observed to simply slip along the muscle surface due to its inability to follow the tissue contour, resulting in extremely noisy signals, i.e. low SNR (**Fig. 4H, figs. S43-44**). Moving forward, our stretchable electrode array may be further extended to soft robotic applications where robust operation under extreme deformation is needed.

Finally, stretchable bioelectronics made with rigid inorganic materials require special structural designs at compromised device densities while intrinsically stretchable TopoE-S allows simultaneously high-density and stretchable electrode array (**fig. S45**), which is critical to the interrogation of biological activities at intricate locations with high resolution. In this regard, brainstem would be a perfect testbed for several reasons (**Fig. 5A, fig. S46**). First, brainstem is naturally curved and will experience severe strains during the movements of cervical spines(*21, 49*). It serves as the central hub for motor controls of almost all facial and neck motions through ten pairs of cranial nerves(*50*). It also regulates cardiac and respiratory functions. Thus, a high-density electrode array will be instrumental for precise control of downstream activities. Furthermore, soft and stretchable devices are essential to mitigate any possible damages from neck movement to the brainstem.

**Fig. 5.**
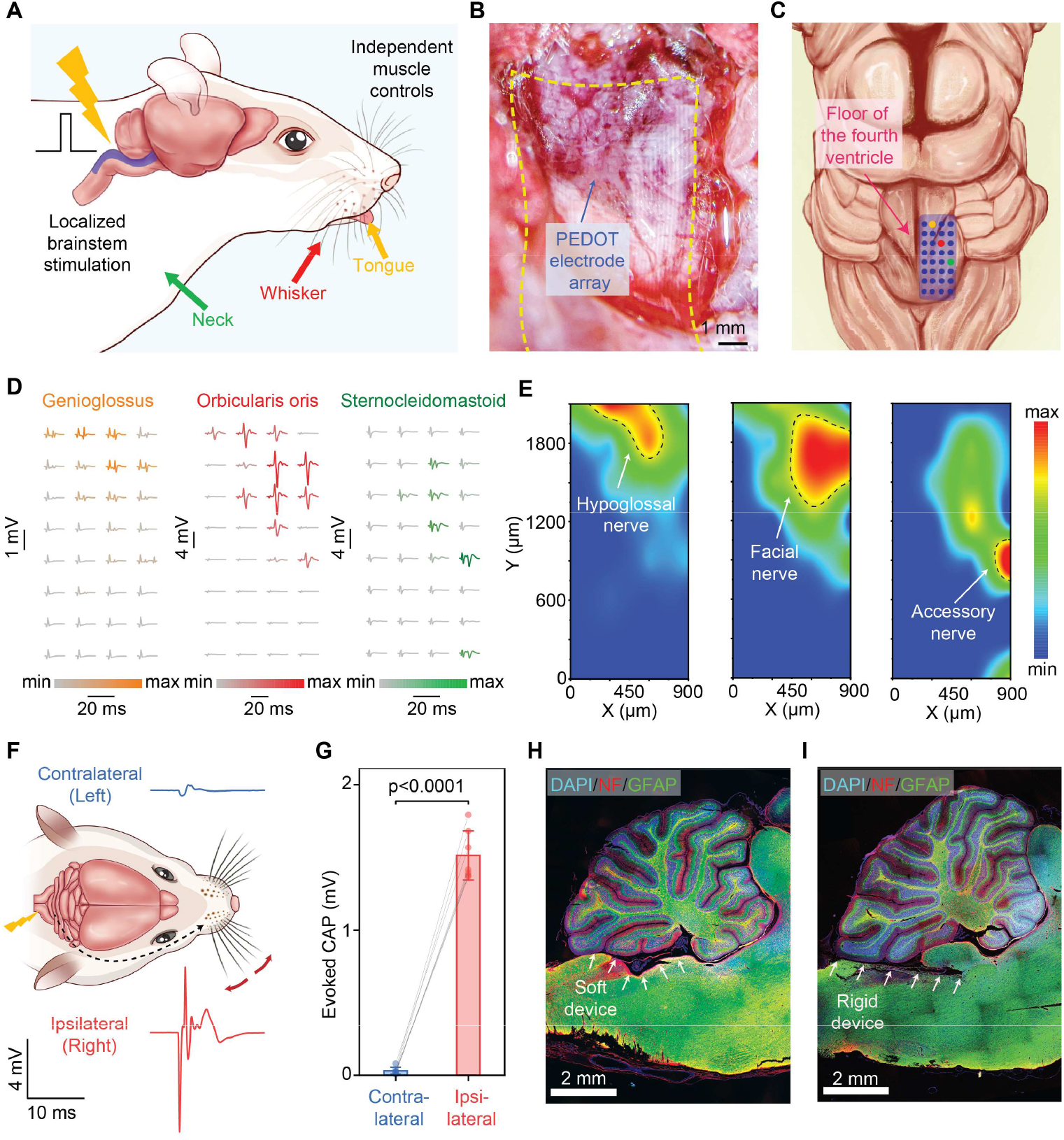
Intrinsically stretchable electrode array allows localized neuromodulation for precise control of independent muscle activities. **(A)** Schematic diagram illustrating the application of the stretchable electrode array for precise neuromodulation through localized brainstem stimulation. **(B)** Microscope image of a stretchable electrode array conforming to the curved floor of the fourth ventricle. **(C)** Schematic illustration of a multi-electrode array placed on the right side of the brainstem. **(D)** Evoked muscle activities recorded at the tongue (left), whisker (middle), and neck (right) following electrical stimulation at the brainstem. **(E)** Activation maps based on the muscle activities depicting the spatial distribution of different nucleus (marked by dashed lines) in the brainstem with downstream connections to the hypoglossal nerve (left), facial nerve (middle), and accessory nerve (right). **(F)** Schematic diagram and representative data traces showing the side specificity of the brainstem stimulation. **(G)** Statistical analyses showing the preferred activation of ipsilateral targets. **(H-I)** Immunohistological staining of a brain slice following the insertion of the soft and stretchable electrode array (**H**) and a rigid device (**I**) along the floor of the fourth ventricle between the brainstem and the cerebellum.

We first showed that our stretchable electrode array, when placed on the floor of the fourth ventricle, could easily follow the underlying curvature to form intimate contact (**Fig. 5B, fig. S46**). Next, we delivered current pulses to individual electrodes to stimulate tongue, whisker, and neck individually and simultaneously recorded EMG and motion signals at those locations were used to confirm the organ-specific stimulation (**Fig. 5C**). After sweeping the entire electrode array with 300 μm IED and 50 μm electrode width, we observed a clear evolution of muscle electrophysiological signals and corresponding movements with unprecedented spatial precision (**Fig. 5D, figs. S47-49**). After normalizing EMG activities elicited by each electrode, we were able to construct three high-resolution activation maps that correlated well with individual nucleuses for hypoglossal, facial, and accessory nerves that innervated genioglossus, orbicularis oris, and sternocleidomastoid, respectively(*50*) (**Fig. 5E, fig. S50**). Moreover, besides independent controls of different muscle groups, the high-resolution stimulation through our stretchable electrode array also showed strong side specificity. Briefly, only the muscles on the same side (i.e., ipsilateral) of the electrode would be selectively stimulated, in accordance with the functional organization of corresponding cranial nerves(*50*) (**Figs. 5F-G, figs. S50-51**). The amplitude of the elicited muscle signals could also be modulated by the intensity of the input stimulus, suggesting the potential to fine tune the neuromodulation with even more controls (**fig. S52**). Finally, immuno-histological analysis indicated that our electrode array did not induce observable tissue damages or inflammatory responses when placed between the cerebellum and the brainstem, whereas rigid plastic probes supported on polyimide substrates with Young’s modulus at a few GPa caused severe damage by cutting into the soft brainstem, possibly due to frequent neck movements, shortly after implantation (**Fig. 5H-I, figs. 53-55**). This work clearly demonstrated the need and importance in using soft electronics for biocompatible neural interfaces. Collectively, our ability to modulate neural activities down to single nucleus precision not only offered immediate benefits for clinical translation in neurosurgical procedures where high-resolution functional maps are demanded, but also opens up a new ‘playground’ for brain-machine interfaces where previously challenging brain structures, such as brainstem, may now be incorporated.

In conclusion, we utilize molecular topology to design a single additive to address the longstanding bottleneck of poor conductivity for stretchable polymer bioelectronics, in which our TopoE-S material achieved record-high conductivity, stretchability, and photo-patternability. Our fabricated soft and biocompatible electrode arrays enabled previously inaccessible applications, including ∼10 times higher resolution EMG recording, unprecedented mapping on a soft live creature, e.g. octopus, and localized neuromodulation down to single nucleus for high-precision control of individual muscle activities through the fragile brainstem. These capabilities may be further extended to construct a stand-alone and closed-loop system that allows ultimate integration with human body. Our transparent and stretchable high conductivity electrode having figure-of-merit which is on par with the widely used ITO will also enable other stretchable electronics applications, such as displays, solar cells, transistors, and photodetectors.

## Acknowledgments

We thank K. Sun, D. Liu, and J. Tang for technical supports. We thank Agfa-Gevaert N.V for providing PEDOT:PSS solution, Asahi Kasei for providing SEBS, and Daikin Co. for providing PVDF-HFP.

## Author contributions

Y.J. and Z.B. conceived the project. Y.J., Z.Z., and Y.-X.W., designed the materials and carried out synthesis. Y.J., Z.Z., Y.-X.W., C.-T.C., D.Z., G.C., H.-C.W., S.N., W.W., A.S., J.L., Y.W., A.T., K.Y.L., C.-C.S., W.X., K.L., and K.Z. performed testing and characterizations. Y.J., E.H. and I.S. designed and performed the octopus-related experiments. Y.J. and D.L. carried out the brainstem-related experiments. Y.J., Z.Z, Y.-X.W., and Z.B. wrote the paper and incorporated comments and edits from all authors.

## Data and materials availability

All data are available in the main text or the supplementary materials.

## Supplementary Material

Materials and Methods

Figs. S1 to S55

References (*1-10*)

## References and Notes

1. K. J. Yu et al., Bioresorbable silicon electronics for transient spatiotemporal mapping of electrical activity from the cerebral cortex. Nat Mater 15, 782–791 (2016). doi:10.1038/nmat4624.

2. S. K. Kang et al., Bioresorbable silicon electronic sensors for the brain. Nature 530, 71–76 (2016). doi:10.1038/nature16492.

3. D. Khodagholy et al., NeuroGrid: recording action potentials from the surface of the brain. Nat Neurosci 18, 310–315 (2015). doi:10.1038/nn.3905.

4. J. Viventi et al., Flexible, foldable, actively multiplexed, high-density electrode array for mapping brain activity in vivo. Nat Neurosci 14, 1599–1605 (2011). doi:10.1038/nn.2973.

5. H. Fang et al., Capacitively Coupled Arrays of Multiplexed Flexible Silicon Transistors for Long-Term Cardiac Electrophysiology. Nat Biomed Eng 1, (2017). doi:10.1038/s41551-017-0038.

6. J. Liu et al., Syringe-injectable electronics. Nature Nanotechnology 10, 629–636 (2015). doi:10.1038/nnano.2015.115.

7. D. Ghezzi et al., A polymer optoelectronic interface restores light sensitivity in blind rat retinas. Nat Photonics 7, 400–406 (2013). doi:10.1038/nphoton.2013.34.

8. Y. Jiang et al., Rational design of silicon structures for optically controlled multiscale biointerfaces. Nat Biomed Eng 2, 508–521 (2018). doi:10.1038/s41551-018-0230-1.

9. K. Mathieson et al., Photovoltaic Retinal Prosthesis with High Pixel Density. Nat Photonics 6, 391–397 (2012). doi:10.1038/nphoton.2012.104.

10. J. Koo et al., Wireless bioresorbable electronic system enables sustained nonpharmacological neuroregenerative therapy. Nat Med 24, 1830–1836 (2018). doi:10.1038/s41591-018-0196-2.

11. A. D. Mickle et al., A wireless closed-loop system for optogenetic peripheral neuromodulation. Nature 565, 361–365 (2019). doi:10.1038/s41586-018-0823-6.

12. F. Michoud et al., Epineural optogenetic activation of nociceptors initiates and amplifies inflammation. Nat Biotechnol, (2020). doi:10.1038/s41587-020-0673-2.

13. I. R. Minev et al., Biomaterials. Electronic dura mater for long-term multimodal neural interfaces. Science 347, 159–163 (2015). doi:10.1126/science.1260318.

14. D. J. Chew et al., A microchannel neuroprosthesis for bladder control after spinal cord injury in rat. Sci Transl Med 5, 210ra155 (2013). doi:10.1126/scitranslmed.3007186.

15. N. Wenger et al., Spatiotemporal neuromodulation therapies engaging muscle synergies improve motor control after spinal cord injury. Nat Med 22, 138–145 (2016). doi:10.1038/nm.4025.

16. M. Capogrosso et al., A brain-spine interface alleviating gait deficits after spinal cord injury in primates. Nature 539, 284–288 (2016). doi:10.1038/nature20118.

17. F. B. Wagner et al., Targeted neurotechnology restores walking in humans with spinal cord injury. Nature 563, 65–71 (2018). doi:10.1038/s41586-018-0649-2.

18. X. Yu et al., Skin-integrated wireless haptic interfaces for virtual and augmented reality. Nature 575, 473–479 (2019). doi:10.1038/s41586-019-1687-0.

19. J. C. Barrese et al., Failure mode analysis of silicon-based intracortical microelectrode arrays in non-human primate. Journal of Neural Engineering 10, 066014 (2013). doi:10.1088/1741-2560/10/6/066014.

20. J. W. Salatino, K. A. Ludwig, T. D. Y. Kozai, E. K. Purcell, Glial responses to implanted electrodes in the brain. Nature Biomedical Engineering 1, 862–877 (2017). doi:10.1038/s41551-017-0154-1.

21. S. P. Lacour, G. Courtine, J. Guck, Materials and technologies for soft implantable neuroprostheses. Nature Reviews Materials 1, 16063 (2016). doi:10.1038/natrevmats.2016.63.

22. E. Song, J. Li, S. M. Won, W. Bai, J. A. Rogers, Materials for flexible bioelectronic systems as chronic neural interfaces. Nature Materials 19, 590–603 (2020). doi:10.1038/s41563-020-0679-7.

23. T. Someya, Z. Bao, G. G. Malliaras, The rise of plastic bioelectronics. Nature 540, 379–385 (2016). doi:10.1038/nature21004.

24. S. Wang et al., Skin electronics from scalable fabrication of an intrinsically stretchable transistor array. Nature 555, 83–88 (2018). doi:10.1038/nature25494.

25. J. Xu et al., Highly stretchable polymer semiconductor films through the nanoconfinement effect. Science 355, 59 (2017). doi:10.1126/science.aah4496.

26. S. Inal, G. G. Malliaras, J. Rivnay, Benchmarking organic mixed conductors for transistors. Nat Commun 8, 1767 (2017). doi:10.1038/s41467-017-01812-w.

27. B. D. Paulsen, K. Tybrandt, E. Stavrinidou, J. Rivnay, Organic mixed ionic–electronic conductors. Nature Materials 19, 13–26 (2020). doi:10.1038/s41563-019-0435-z.

28. H. Yuk, B. Lu, X. Zhao, Hydrogel bioelectronics. Chem Soc Rev 48, 1642–1667 (2019). doi:10.1039/c8cs00595h.

29. T. Adrega, S. P. Lacour, Stretchable gold conductors embedded in PDMS and patterned by photolithography: fabrication and electromechanical characterization. Journal of Micromechanics and Microengineering 20, 055025 (2010).

30. T. Baetens, E. Pallecchi, V. Thomy, S. Arscott, Cracking effects in squashable and stretchable thin metal films on PDMS for flexible microsystems and electronics. Sci Rep 8, 9492 (2018). doi:10.1038/s41598-018-27798-z.

31. Y. Gu et al., Photoswitching topology in polymer networks with metal–organic cages as crosslinks. Nature 560, 65–69 (2018). doi:10.1038/s41586-018-0339-0.

32. L. F. Hart et al., Material properties and applications of mechanically interlocked polymers. Nature Reviews Materials, (2021). doi:10.1038/s41578-021-00278-z.

33. S. Choi, T. W. Kwon, A. Coskun, J. W. Choi, Highly elastic binders integrating polyrotaxanes for silicon microparticle anodes in lithium ion batteries. Science 357, 279–283 (2017). doi:10.1126/science.aal4373.

34. A. Bin Imran et al., Extremely stretchable thermosensitive hydrogels by introducing slide-ring polyrotaxane cross-linkers and ionic groups into the polymer network. Nature Communications 5, 5124 (2014). doi:10.1038/ncomms6124.

35. H. Gotoh et al., Optically transparent, high-toughness elastomer using a polyrotaxane cross-linker as a molecular pulley. Science Advances 4, eaat7629 (2018). doi:10.1126/sciadv.aat7629.

36. J. Rivnay et al., Structural control of mixed ionic and electronic transport in conducting polymers. Nat Commun 7, 11287 (2016). doi:10.1038/ncomms11287.

37. Y. Wang et al., A highly stretchable, transparent, and conductive polymer. Sci Adv 3, e1602076 (2017). doi:10.1126/sciadv.1602076.

38. Y. Liu et al., Soft and elastic hydrogel-based microelectronics for localized low-voltage neuromodulation. Nat Biomed Eng 3, 58–68 (2019). doi:10.1038/s41551-018-0335-6.

39. D. Alemu Mengistie, P.-C. Wang, C.-W. Chu, Effect of molecular weight of additives on the conductivity of PEDOT:PSS and efficiency for ITO-free organic solar cells. Journal of Materials Chemistry A 1, 9907 (2013). doi:10.1039/c3ta11726j.

40. N. Kim et al., Highly Conductive PEDOT:PSS Nanofibrils Induced by Solution-Processed Crystallization. Advanced Materials 26, 2268–2272 (2014). doi:10.1002/adma.201304611.

41. S. M. Kim et al., Influence of PEDOT:PSS crystallinity and composition on electrochemical transistor performance and long-term stability. Nat Commun 9, 3858 (2018). doi:10.1038/s41467-018-06084-6.

42. D. S. Hecht et al., Carbon-nanotube film on plastic as transparent electrode for resistive touch screens. Journal of the Society for Information Display 17, 941 (2009). doi:10.1889/jsid17.11.941.

43. L. Hu, H. S. Kim, J. Y. Lee, P. Peumans, Y. Cui, Scalable coating and properties of transparent, flexible, silver nanowire electrodes. ACS Nano 4, 2955–2963 (2010). doi:10.1021/nn1005232.

44. Z. Chen et al., Fabrication of Highly Transparent and Conductive Indium–Tin Oxide Thin Films with a High Figure of Merit via Solution Processing. Langmuir 29, 13836–13842 (2013). doi:10.1021/la4033282.

45. S. Savagatrup et al., Plasticization of PEDOT:PSS by Common Additives for Mechanically Robust Organic Solar Cells and Wearable Sensors. Advanced Functional Materials 25, 427–436 (2015). doi:10.1002/adfm.201401758.

46. C. L. Choong et al., Highly stretchable resistive pressure sensors using a conductive elastomeric composite on a micropyramid array. Adv Mater 26, 3451–3458 (2014). doi:10.1002/adma.201305182.

47. N. Matsuhisa et al., High-Transconductance Stretchable Transistors Achieved by Controlled Gold Microcrack Morphology. Advanced Electronic Materials 5, 1900347 (2019). doi:10.1002/aelm.201900347.

48. B. Afsharipour, S. Soedirdjo, R. Merletti, Two-dimensional surface EMG: The effects of electrode size, interelectrode distance and image truncation. Biomedical Signal Processing and Control 49, 298–307 (2019). doi:10.1016/j.bspc.2018.12.001.

49. N. Vachicouras et al., Microstructured thin-film electrode technology enables proof of concept of scalable, soft auditory brainstem implants. Sci Transl Med 11, (2019). doi:10.1126/scitranslmed.aax9487.

50. D. E. Haines, Neuroanatomy: An Atlas of Structures, Sections, and Systems. (Lippincott Williams & Wilkins, ed. 8th edition, 2011).

