## Supplementary Information for "Topological supramolecular network enabled highly conductive and stretchable organic bioelectronics"

**This PDF file includes:**

Materials and Methods
Figs. S1 to S55

### Materials and Methods

#### I. Synthesis of polyrotaxane-based supramolecular crosslinkers

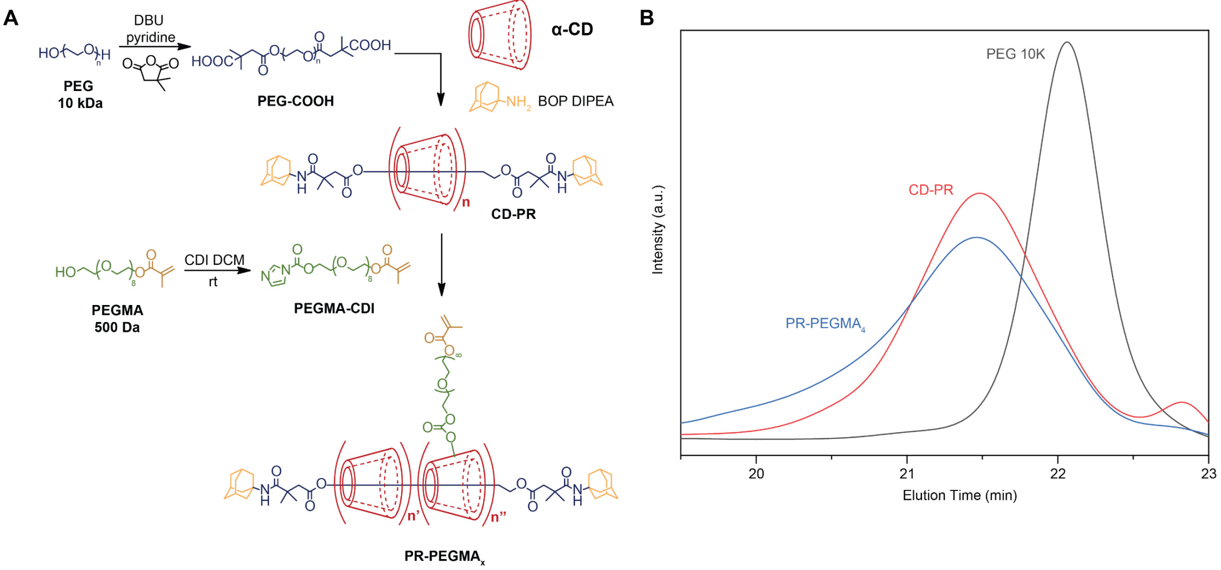

**Fig. S1. Synthetic route for PR-PEGMA (A) and SEC characterizations (B).** The final PR-PEGMA consists of a PEG 10 kDa backbone, adamantane stoppers, cyclodextrin rings, PEG 500 sidechains, and methacrylate terminals. For CD-PR, *n* is confirmed as ca. 10 by $^1$H NMR integration ratio of the cyclodextrin to the PEG axle. For PR-PEGMA$_x$, $n'$ (or $n''$) represents the number of grafted (or non-grafted) cyclodextrins, and *x* represents the average number of grafted PEGMAs on a polyrotaxane, which is calculated from $^1$H NMR integrations of the methacrylate and the PEG backbone.

(1) PEG-COOH

Polyethylene glycol (PEG, 10.0 g, average molecular weight 10,000, Sigma-Aldrich) was first dissolved in anhydrous pyridine (50 mL, Sigma-Aldrich) before 1,8-diazabicyclo[5.4.0]undec-7-ene (DBU, 250 µL, Sigma-Aldrich) and 2,2-dimethylsuccinic anhydride (6.0 mL, Sigma-Aldrich) were added. The solution was stirred at 60 °C overnight under nitrogen atmosphere. The reaction solution was dropped into Milli-Q water (200 mL), and then the pH was adjusted to 2-3 using concentrated hydrochloric acid (HCl, Sigma-Aldrich). The product was extracted with dichloromethane (DCM, Fisher Scientific) and then dried to obtain PEG-COOH (9.5 g) as a white solid.

(2) CD-PR

PEG-COOH (1.5 g) was dissolved in deionized water (10.0 mL), and then the aqueous solution of *α*–cyclodextrin (α-CD, 6.00 g/44 mL, Sigma-Aldrich) was added. The mixed solution was stirred for 30 minutes and then stored in a refrigerator for two days to obtain a turbid solution with white precipitate. The mixture was freeze-dried to obtain a white powder containing pseudo-polyrotaxane.

For the end-capping reaction, a solution of 1-adamantaneamine (195 mg, 1.3 mmol, Sigma-Aldrich), (benzotriazol-1-yloxy)tris(dimethylamino)phosphonium hexafluorophosphate (BOP reagent, 570 mg, 1.3 mmol, Sigma-Aldrich) and *N*-ethyldiisopropylamine (DIPEA, 222 µL, 1.3 mmol, Sigma-Aldrich) in anhydrous acetonitrile (75 mL, Sigma-Aldrich) was mixed with the powder of pseudo-polyrotaxane under nitrogen atmosphere. The suspension was stirred at room temperature for 48 hours. The resultant suspension was centrifuged to remove the supernatant. The obtained solid was washed with acetonitrile twice, and then the solid was dried under vacuum. The obtained white solid was dissolved in dimethyl sulfoxide (DMSO, 50 mL), and the solution was dropped into deionized water (500 mL). This aqueous solution was purified by dialysis tubing. Finally, the resultant solution was freeze-dried after filtrated by nylon filters (pore size: 1 µm, Whatman) to obtain CD-PR (1.86 g) as a white solid.

(3) PEGMA-CDI

An anhydrous DCM solution (30 mL, Sigma-Aldrich) of poly(ethylene glycol) methacrylate (PEGMA, 2g, Mn 500, Sigma-Aldrich) was added into another anhydrous DCM solution (50 mL) of 1,1'-carbonyldiimidazole (CDI, 0.71 g, 1.1 eq, Tokyo Chemical Industry). The clear solution was stirred at room temperature (rt) under nitrogen atmosphere for one day followed by saline washing (3 × 100 mL). The organic phase was dried with anhydrous sodium sulfate (Fisher Scientific) and concentrated in vacuo to give a yellow viscous liquid (yield: 2.1 g).

(4) PR-PEGMA*x*

CD-PR, PEGMA-CDI with different equivalents (2, 5, 10) to *α*-CD in CD-PR together and 4-(dimethylamino) pyridine (DMAP, 0.05 eq to PEGMA-CDI) were dissolved in anhydrous DMSO and the mixture was stirred at 35 ºC for 3 days. Phosphate buffered saline (PBS, Thermo-Fisher) solution was added to dilute DMSO, and the solution was dialyzed against water, followed by freeze drying to give the product as a white solid. The average number of PEGMA group on one CD-PR was determined by $^{1}$H NMR spectra.

Chemical structures of the intermediates and final products were confirmed by nuclear magnetic resonance (NMR) spectroscopy using a 500MHz Varian Inova spectrometer and Fourier transform infrared (FT-IR) spectroscopy using a Nicolet iS50 FT/IR spectrometer. Molecular weights were characterized by size exclusion chromatography (SEC) performed on a Viscotek GPC270 at 323 K with 0.8 mL/min, using refractive index detection, poly(methyl methacrylate) (PMMA) standards, and DMF as the eluent.

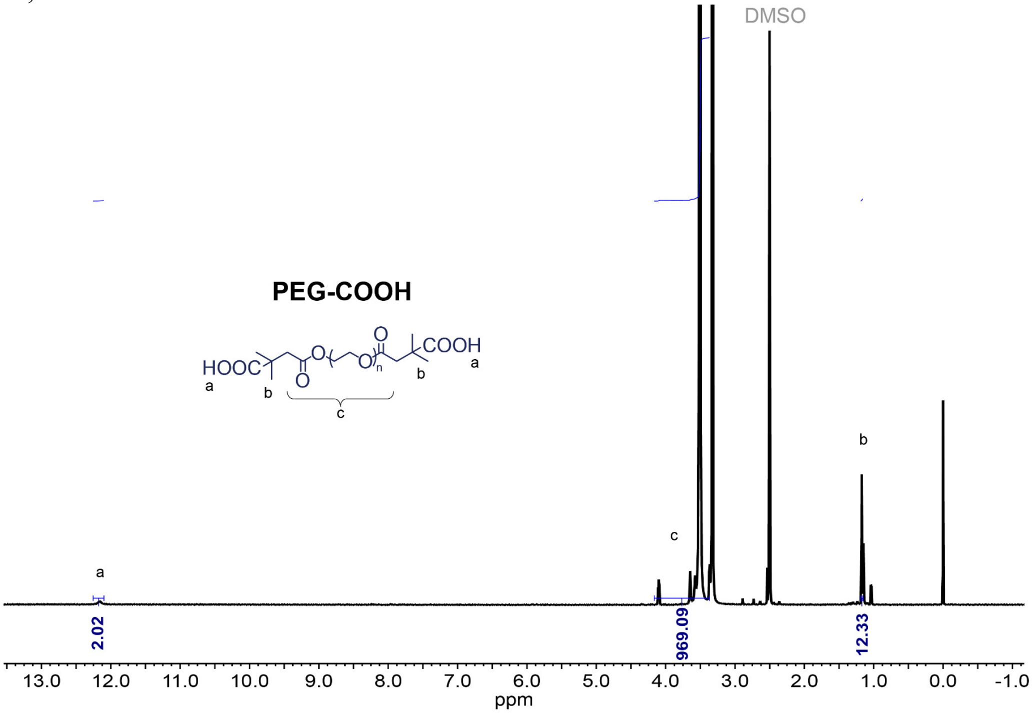

**Fig. S2. NMR spectrum of PEG-COOH in DMSO-*d*$_6$.**

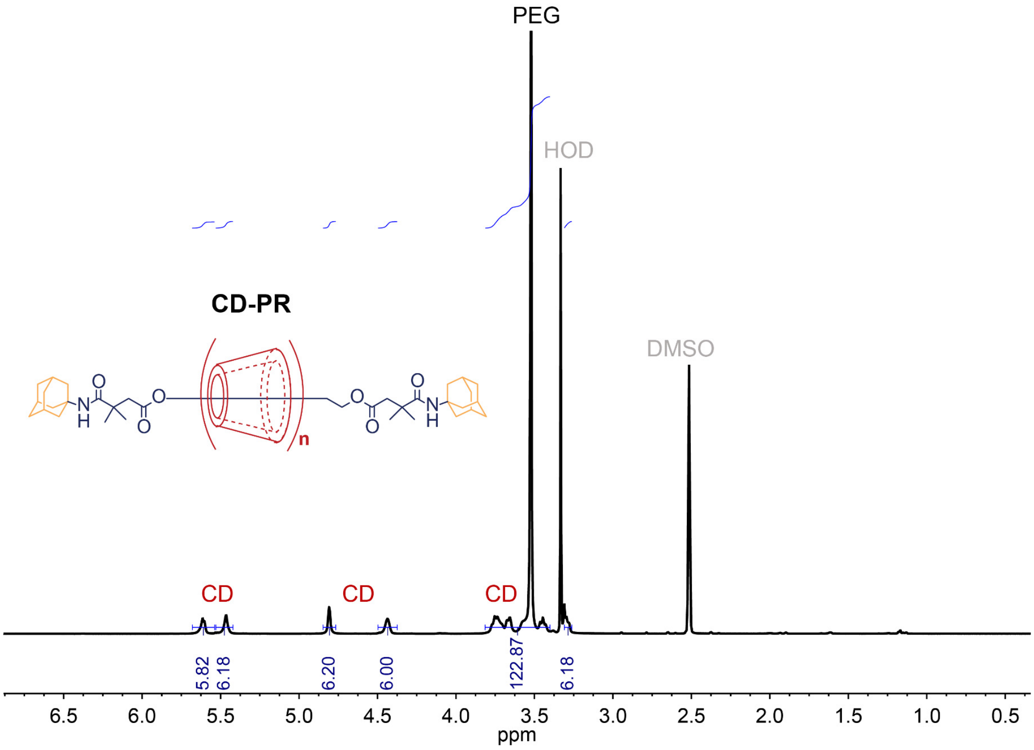

**Fig. S3. NMR spectrum of PR in DMSO-$d_6$.**

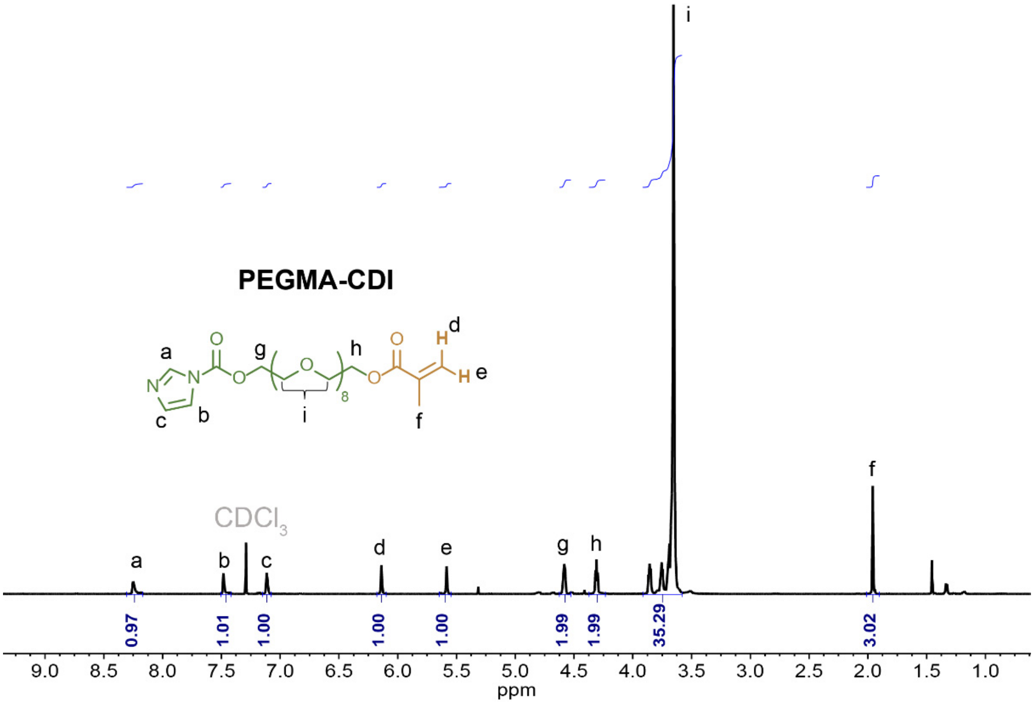

**Fig. S4. NMR spectrum of PEGMA-CDI in $CDCl_3$.**

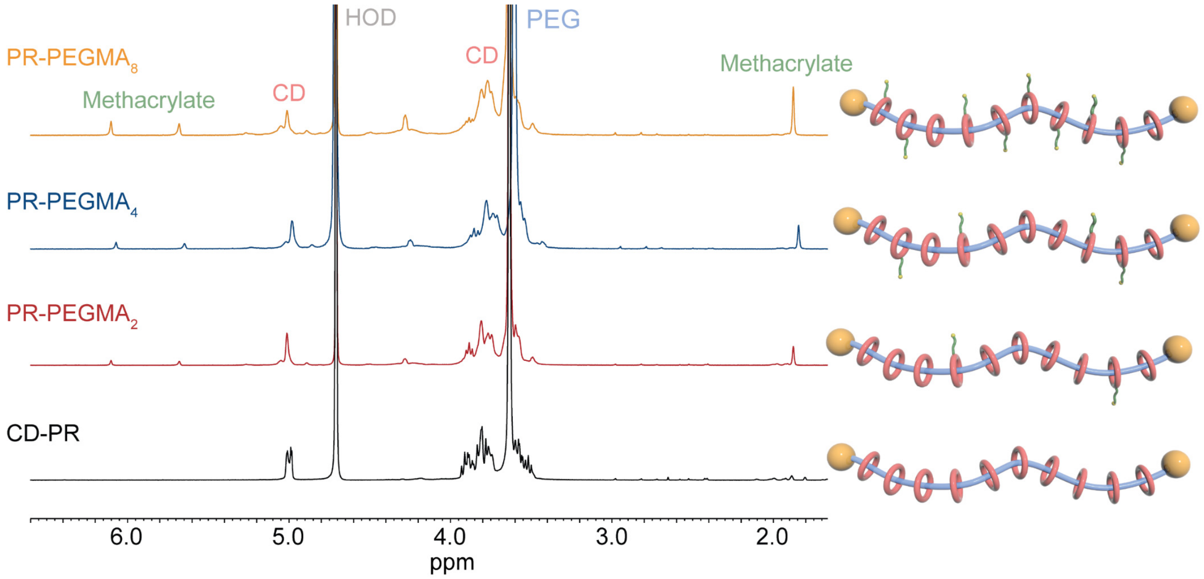

**Fig. S5. NMR spectra of PR-PEGMA with different number of sidechains in $D_2O$.**

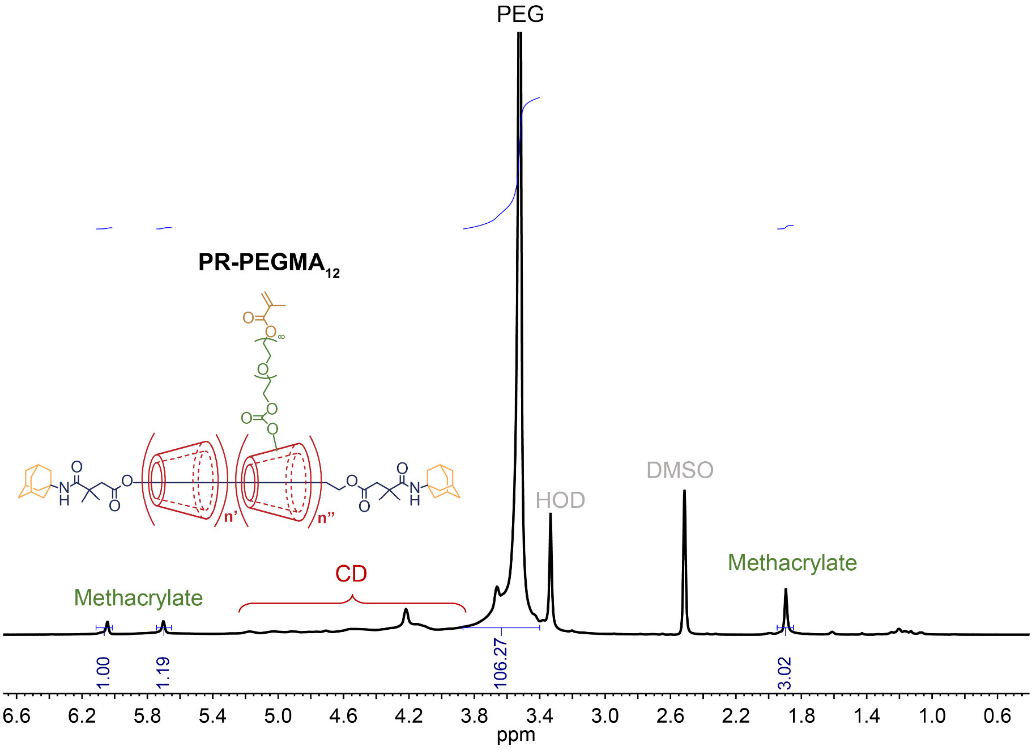

**Fig. S6. NMR spectrum of PR-PEGMA$_{12}$ in DMSO-$d_6$.**

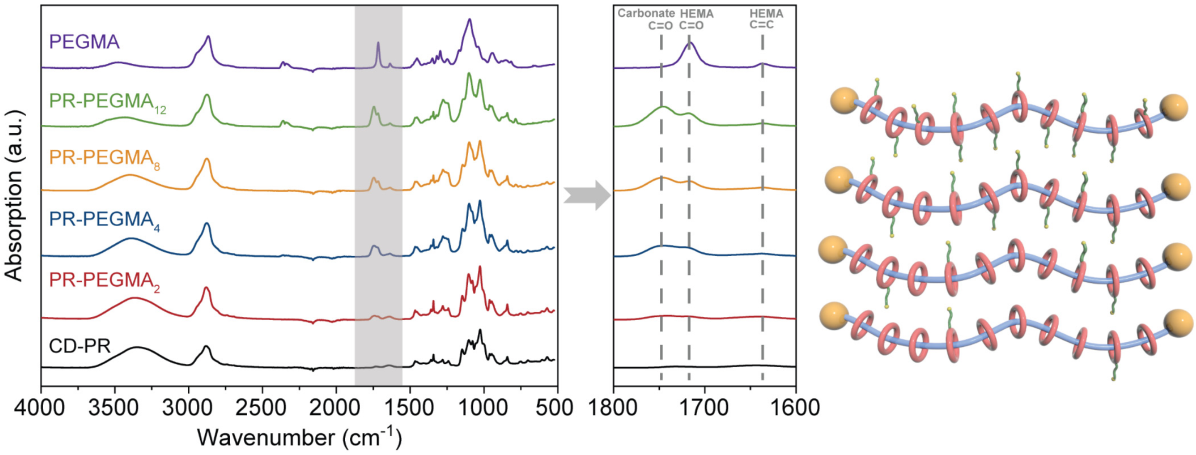

**Fig. S7. FT-IR spectra of PR-PEGMA with different number of sidechains.** For PR-PEGMA$_x$, the appearance of peaks at 1717 and 1636 cm$^{-1}$ is indicative of the successful grafting of methacrylate terminals, and the peak at 1747 cm$^{-1}$ clearly proves the formation of carbonate linkage as shown in the synthetic route (**Fig. S1**).

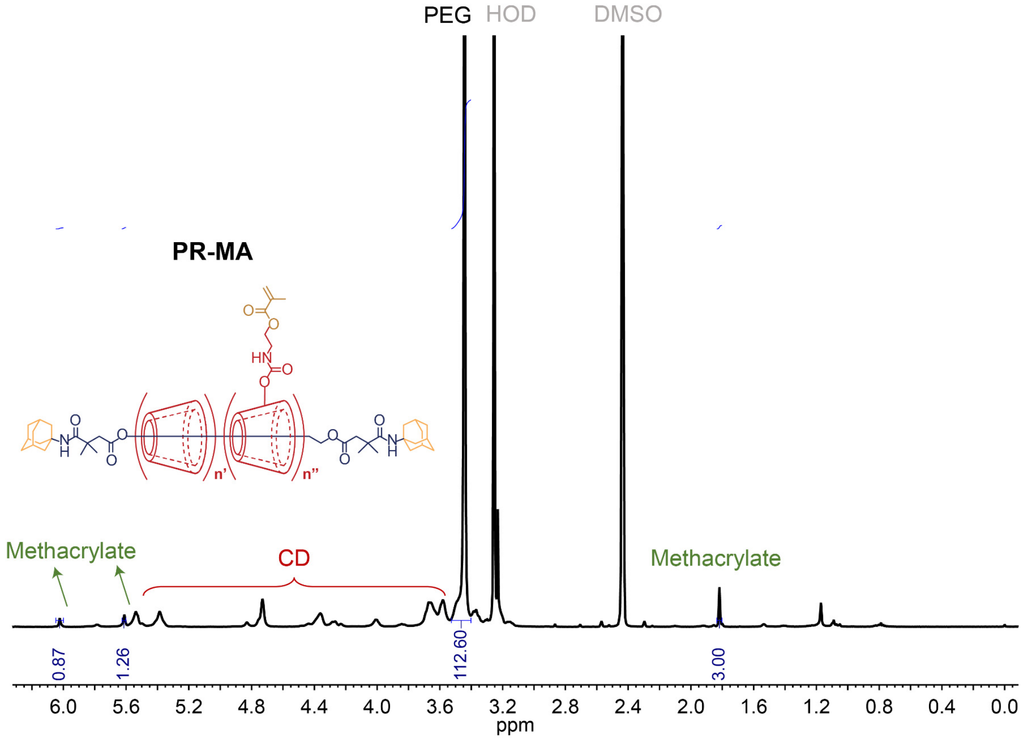

**Fig. S8. NMR spectrum of PR-MA in DMSO-$d_6$.** The average number of grafted methacrylates on a polyrotaxane is calculated to be 8 according to $^1$H NMR integrations of the methacrylate and the PEG backbone.

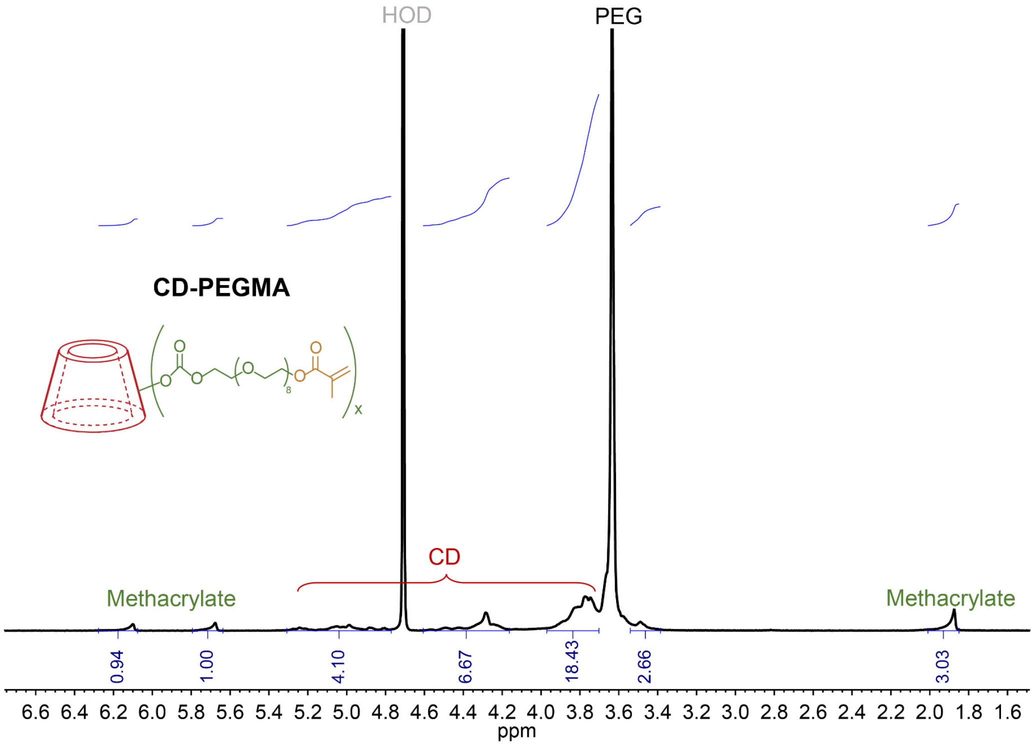

**Fig. S9. NMR spectrum of CD-PEGMA in $D_2O$.** The average number of grafted methacrylates on a cyclodextrin is calculated as 1.3 according to $^1$H NMR integrations of the methacrylate and the cyclodextrin.

**(II) Preparation of TopoE, TopoE-S and control sample films**

Poly(3,4-ethylenedioxythiophene) polystyrene sulfonate (PEDOT:PSS, Orgacon™ ICP 1050) was kindly provided by Agfa-Gevaert N.V as a surfactant-free aqueous dispersion with 1.1 wt% solid content. PR-based supramolecular crosslinkers were synthesized as above. Control samples of poly(ethylene glycol) diacrylate (PEGDA, Mn 10,000 or 575) were purchased from Sigma-Aldrich. Multi-arm PEG acrylate samples with molecular weight of 10,000 (8-arm PEG acrylate-tripentaerythritol, or 8-arm PEG acrylate-hexaglycerol) were purchased from JenKem Technology.

In a typical experiment, PR-PEGMA of various amounts (11 mg, 27.5 mg, 55 mg) together with 0.5 mg of photo initiator, 2-hydroxy-4′-(2-hydroxyethoxy)-2-methylpropiophenone (Irgacure 2959, Sigma-Aldrich), were added to 1 mL of PEDOT:PSS aqueous dispersion (1.1 wt%) and mixed using a vortex mixer. After filtering using nylon syringe filters (pore size: 1 μm, Whatman), the mixture would be spin coated onto oxygen ($O_2$) plasma treated glass, silicon (Si)/silicon dioxide ($SiO_2$), or elastomeric substrates at 2000 rpm for 1 min. Typically, the plasma treatment condition was 150 W for 30 s using a Technics Micro-RIE Series 800. The spin coated TopoE film would be later photo-crosslinked by UV exposure for 2 min using a Spectrum 100 Precision UV spot curing system (American Ultraviolet, 25 mW/cm$^2$) before rinsing by water. The as-prepared TopoE film would be further treated with acid by dipping into concentrated sulfuric acid (Sigma-Aldrich) for 10 s to yield the TopoE-S film for further testing and characterizations.

For all control films, 55 mg of control molecules were added to 1 mL of PEDOT:PSS dispersion with 0.5 mg of Irgacure 2959. The films were spin coated onto elastomeric substrates at 2000 rpm for 1 min before UV cured for 2 min using the same Spectrum 100 UV curing system before water development. No acid treatment was performed on the control samples.

**(III) Material and chemical characterizations**

Atomic force microscopy (AFM) images were collected in tapping mode using a Bruker Icon AFM. X-ray photoelectron spectroscopy (XPS) was performed on a PHI VersaProbe Scanning XPS Microprobe. Gas cluster ion beam (GCIB) sources were used for depth profiling to avoid damage of the PEDOT:PSS structure. UV-vis-NIR spectra were collected on an Agilent Cary 6000i model. A rotational polarizer was used to measure the absorption intensity with the polarization parallel and perpendicular to the stretching direction. Raman spectroscopy were performed using a Horiba Labram HR Evolution Raman System. Film thickness values were measured using a contact probe Dektak 150 profilometer. Differential scanning calorimetry (DSC) was conducted using a TA Instruments Q2000 DSC. Grazing incidence X-ray diffraction (GIXD) measurements were carried out at the Stanford Synchrotron Radiation Lightsource at beamline 11-3, with a photon energy of 12.735 keV and a sample-to-detector distance of 320 mm. The incident angle was fixed at 0.14° to probe the entire film with reduced substrate scattering.

**(IV) Mechanical characterizations**

For microscopic investigation of crack formation, PEDOT:PSS thin films with various compositions were hand stretched to different lengths corresponding to strain levels of 25%, 50%, 75%, and 100% before imaging under a upright microscope (Leica).

For the measurement of Young's modulus, PEDOT:PSS solutions mixed with different PR concentrations were drop-casted onto Si wafers and dried to form thick films of ~ 15 μm. The as-prepared Si wafers would be later mounted onto an aluminum puck using graphite paste. A Berkovich tip with a dynamic indentation method was used to probe the modulus of the films, and the measurement was performed on a Nanomechanics iNano Nanoindenter.

**(V) Electrical and electrochemical characterizations**

Conductivity measurements were carried out using a four-point probe method using a Keithley 4200 SC semiconductor analyzer. Electrodes were deposited by applying silver paste with 1 mm interelectrode spacing ($s$). The conductivity $\sigma$ is calculated by $\sigma=1/(2\pi sV/I)$ where $V$ is the potential difference between probe 2 and 3, $I$ is the current running through probe 1 and 4. A minimum of four measurements were obtained for an average value.

For the measurement of resistance change over strain, PEDOT:PSS films with different formulations would be mounted onto a home-made automated stretcher when the resistance values at different strain levels would be recorded using a LCR meter (Keysight Technologies, E4098A). Contacts between PEDOT:PSS and the LCR meter were made by silver paste and eutectic gallium-indium (EGaIn) liquid metal alloy.

Electrochemical impedance spectroscopy (EIS) was performed by using TopoE-S as the working electrode, platinum as the counter electrode, silver/silver chloride (Ag/AgCl) as the reference electrode, and PBS as the electrolyte. The exposed area for TopoE-S electrode was kept at 4 mm × 5 mm with 1 μm dry thickness. Impedance and phase angle as functions of frequency were acquired by a Bio-Logic VSP-300 workstation with a sine wave signal amplitude of 10 mV. Chronopotentiometry was conducted using the same Bio-Logic electrochemical workstation when a biphasic current pulse of plus/minus 5 mA with a duration of 2 ms for each phase was applied. Voltage between the working and the reference electrodes were recorded to compare the charge injection capacities. Chronoamperometry was performed using a pulse measure unit and a 4225-remote preamplifier/switch module connected to the Keithley 4200-SCS. Square wave voltage pulses of 100 mV and a duration of 50 ms were applied and currents were recorded simultaneously.

**(VI) Device fabrication of TopoE-S based stretchable electrode array**

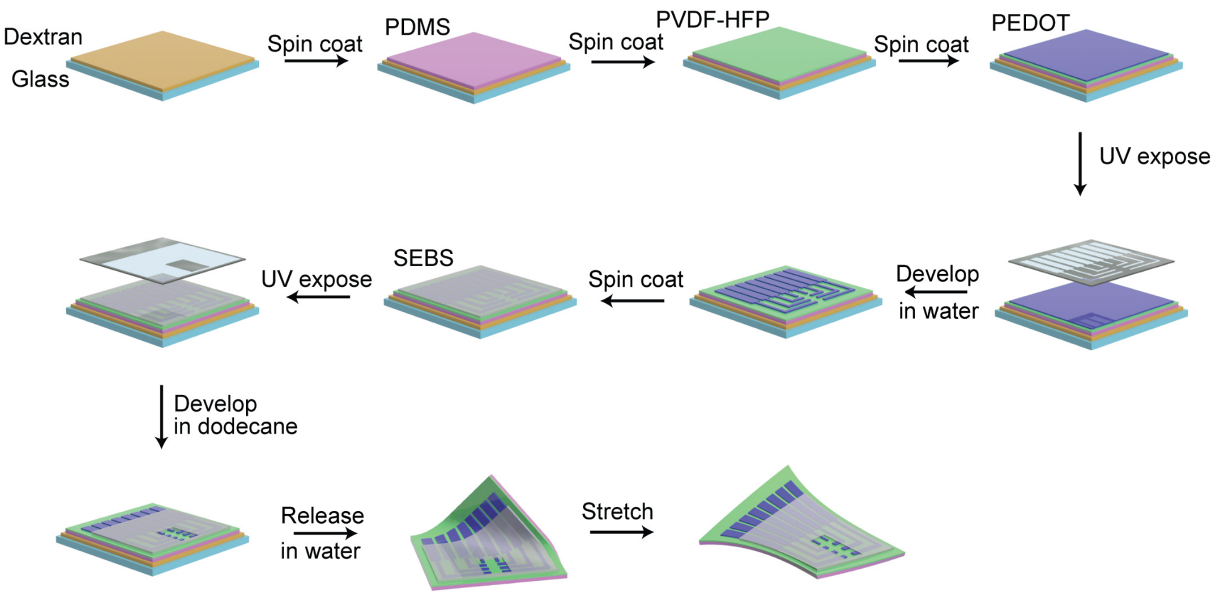

**Fig. S10. Schematic diagram illustrating the fabrication process of fully stretchable multielectrode array.**

In consideration of the chemical orthogonality and the surface energy of each elastomeric layers, a multi-layer device structure for the stretchable electrode array was developed including a polydimethylsiloxane (PDMS) backing layer for handling, a poly(vinylidene fluoride)-co-hexafluoropropylene (PVDF - HFP) substrate layer for favored surface energy and solvent resistance, a PR-PEGMA/PEDOT:PSS conducting layer for electrode arrays and interconnects, and a styrene-ethylene-butylene-styrene (SEBS) encapsulation layer for passivation of interconnects.

Glass substrates were first treated with $O_2$ plasma at 150 W for 30 s using a Technics Micro-RIE Series 800. Next, a dextran (from *Leuconostoc* spp., $M_r$~100,000 Sigma-Aldrich) aqueous solution (5 wt%) was spin coated at 500 rpm for 15 s as the sacrificial layer. The PDMS carrier layer was made by spin coating from Sylgard 184 (PDMS:crosslinker 10:1 weight ratio, Dow) at a speed of 2000 rpm for 20 s, and followed by annealing at 150 °C for 10 min. After curing, the PDMS substrate was treated with oxygen plasma at 150 W for 1 min before spin coating the next PVDF-HFP layer.

PVDF-HFP based fluoroelastomer (DAI-EL G-801) was provided by Daikin Co. as a peroxide curable copolymer. Before spin coating, a methyl ethyl ketone (MEK) solution of PVDF-HFP, triallyl isocyanurate (TAIC, Tokyo Chemical Industry), and benzoyl peroxide (Sigma-Aldrich) was first prepared with a weight ratio of 100:6:2 and the PVDF-HFP concentration of 125 mg/mL. After filtering through nylon syringe filters (pore size: 1 μm, Whatman), the mixture was spin coated at 1000 rpm for 2 min followed by curing at 180 °C for 1 hour.

Next, the PVDF-HFP layer was treated by oxygen plasma at 150 W for 3 min before spin coating the PEDOT:PSS layer. The PEDOT:PSS solution was consisting of 55 mg of PR-PEGMA and 0.5 mg of Irgacure 2959 dissolved in 1 mL of PEDOT:PSS aqueous dispersion (1.1 to 1.3 wt %). After filtering using nylon syringe filters (pore size: 1 μm, Whatman), the mixture was spin coated at 2000 rpm for 1 min. The PR-PEGMA/PEDOT:PSS film was later exposed using a Spectrum 100 Precision UV Spot Curing System (American Ultraviolet) before using water to develop the unexposed part. The as-patterned PEDOT:PSS array would be later treated with concentration sulfuric acid for 10 s before water rinsing and drying.

For the encapsulating SEBS layer, it was first prepared by mixing SEBS (H1062, free sample from Asahi Kasei, 80 mg/mL) and home-made azide crosslinker (decane-1,10-diyl bis(4-azido-2,3,5,6-tetrafluorobenzoate), 2.4 mg/mL) in toluene (Fisher Scientific). After filtering through Teflon syringe filters (pore size: 1 μm, Whatman), the mixture was spin coated at 500 rpm for 2 min followed by UV exposure for 10 min using the Spectrum 100 curing system. After a post-exposure bake at 120 $^{o}$C for 10 min, the unexposed part would be later developed using dodecane (Tokyo Chemical Industry) for 2 min.

For the subsequent bonding process to make input/output connections with external circuits, silver nanowires (AW060, Ke-Chuang Advanced Materials) were spray coated onto the contacting pads through shadow masks. The final contacts were made by sandwiching an anisotropic conductive adhesive (ACA, 3M 9703) between the contacting pads and a flexible flat cable (FFC).

The as-fabricated device would later be released from the substrate by immersing into water for at least 1 hour to dissolve the sacrificial dextran layer.

### (VII) Electrophysiological recording on human

In a typical recording, a 64-channel stretchable array with electrode size of 100 μm and inter-electrode distance of 500 μm was fabricated as described above. A customized printed circuit board (PCB) containing a 64-channel zero-insertion force (ZIF) connector (500 μm pitch, 150 mm length, Uxcell) and two Omnetics connectors (NPD-36-VV-GS, 36-Position Dual Row Male Nano-Miniature Connector) was manufactured to interface the FFC cable and two bipolar-input recording headstages based on RHD2216 digital electrophysiology interface chips (Intan Technologies).

The filter was set to an analogue bandpass of 0.1-6 kHz with a digital filter cutoff of 1 Hz. The single channel sample rate was set to 30 kHz. For hardware control, an Open Ephys acquisition board was used for the communication with other digital devices and the streaming of all the surface electromyography (sEMG) data from the RHD2000 amplifiers. The USB port of the module was linked with a USB cable to pipe the data stream in to and out of the computer.

The recorded data were stored as raw signals from the amplifiers. After filtering with a 60-Hz notch filter, a bandpass filter between 100 Hz and 1000 Hz was applied. Dynamic heat maps showing the spatiotemporal evolution of muscle activities were calculated based on moving averages of the envelope profile (obtained using a second order Butterworth filter with a cutoff frequency of 20 Hz) with a window size of 3.3 ms. Static heat maps were calculated based on the integration of raw EMG activities over time and normalized to assign the color code.

Recording of sEMG on human arms was pre-approved by Institutional Review Board (IRB) at Stanford University (IRB-54795, Noninvasive surface measurement of muscle electrophysiology). All participants are co-authors of the manuscript and have provided consents for participation.

### (VIII)Electrophysiological recording on octopus

All procedures were approved by the Institutional Animal Care and Use Committee (IACUC) for Stanford University with further guidance from the Marine Biological Laboratories Marine Resource Center. Adult California two-spot octopuses (Octopus bimaculoides) were ordered from Aquatic Research Consultants (San Pedro, CA).

Prior to the experiment, an octopus would be anesthetized by adding 2% ethanol into natural seawater and transferred to a customized Styrofoam container. During the recording, a 32-channel stretchable array with electrode size of 500 μm and inter-electrode distance of 1500 μm would be attached onto one of the octopus's arms. The control probe with a rigid substrate was fabricated using the same procedure as the experimental group but using polyimide of the same thickness as the carrier instead of PDMS. A customized PCB with a 32-channel ZIF connector (500 μm pitch, 50 mm length, Digi-Key) and an Omnetic connector was made for input/output connection. A unipolar-input recording headstage based on a RHD2132 digital electrophysiology interface chip (Intan Technologies) was used for the recording.

Electrical stimulation of the octopus arm was performed by applying a 2-ms long pulse of 8 mA with two electrodes placed near the base and tip of the arm using a Model A360 analog stimulus isolator (World Precision Instruments). Raw signals were recorded using the Open Ephys acquisition system (http://open-ephys.org/). The filter was set to an analogue bandpass of 0.1-6 kHz with a digital filter cutoff of 1 Hz. The single channel sample rate was set to 30 kHz. The recorded data were stored as raw signals from the amplifiers. A bandpass Butterworth filter between 60 Hz and 800 Hz was applied. Differences between adjacent electrodes on the same row were calculated to plot the peristimulus time histogram (PSTH). The frequency spectra and level of variances were calculated using custom-written code in Python.

### (IX) Electrical stimulation on rat

All procedures were approved by the Institutional Animal Care and Use Committee (IACUC) for Stanford University with the protocol number of APLAC-33717 (Soft polymeric electronics for neural interfaces). Female Sprague Dawley rats were ordered from Charles River Laboratories with weight of 150 g at the time of arrival. All rats were housed under 12 h:12 h light/dark cycles until experiments.

Before the surgery to interface the stretchable electrode array with the brainstem, the rat was deeply anesthetized with isoflurane (2 L/min $O_2$ mixed with 3% isoflurane). The level of anesthesia was continuously monitored based on whisker movements and paw-pinching/eye-blinking reflexes. After anesthesia, the rat head was shaved with its temperature kept at 37°C with a heating pad. A mid-line skin incision was first made over the craniovertebral junction and the overlying muscles were cut and retracted to expose the occipital bones. An occipital craniotomy was performed using a dental drill, and the dura was peeled using a fine tweezer to ensure a full exposure of the cerebellum and brainstem. Using a surgical cerebral plate, the cerebellum was lifted to expose the floor of the $4^{th}$ ventricle where a stretchable electrode array would be subsequently attached to. To fit the size of the rat brainstem, a 32-channel array with electrode size of 50 μm and inter-electrode distance of 300 μm was used.

After the implantation of the electrode array, subsequent stimulations were performed by applying square wave current pulses of 200 mA for 1 ms through an A-M Systems Model 3800 stimulator coupled with a Model 3820 stimulus isolation unit. Following the localized stimulation through brainstem, EMG signals were simultaneously recorded by inserting needle electrodes at the tongue, whisker, and neck. The recorded signals were amplified by three BioPac electromyogram amplifiers (EMG100C) at a sampling rate of 20 kHz. The recorded data were later processed after a 60 Hz notch filter and a 100-1000 Hz bandpass filter. The activation maps were calculated by normalizing the evoked EMG signals recorded at each muscle group.

**(X) Immunohistology**

Following the implantation of both soft and rigid probes at the brainstem/cerebellum interface for one day, rats were deeply anesthetized and transcardially perfused with saline followed by 4% paraformaldehyde in PBS. The devices were carefully explanted before the brains were removed and fixed in 4% paraformaldehyde in PBS overnight at 4 °C. The brains were subsequently dehydrated using gradient washing of ethanol/PBS mixtures and embedded using paraffin. Final blocks were sectioned in the sagittal plane at 8 μm on a Leica microtome (RM2255) and mounted onto glass slides.

For immunohistochemical analysis, slides were first deparaffinized in xylene, and then rehydrated in ethanol/PBS mixtures. The slides were then washed three times in PBS. Next the slices underwent antigen retrieval using 1× sodium citrate pH 6.0 (100× diluted in PBS; Abcam) in deionized (DI) $H_2O$. The slices, submerged in the solution, were warmed in a microwave on full power for 90 seconds. One minute later, at 60% power for 60 seconds. One minute later, the slices in solution were places in 4 °C for 30 minutes and subsequently soaked in DI $H_2O$ for 5 minutes. The slices were then washed three times in PBS.

Slices were permeabilized for intracellular antigens with 0.2% Trion X-100 (Sigma-Aldrich) for 10 minutes, then washed three times with PBS. A PAP/hydrophobic pen was used to isolate slices. Slices were then blocked with a solution consisting of 5% goat serum (vol/vol; Sigma-Aldrich) in PBS for 2 hours at room temperature in a humidified chamber.

Slices were then incubated for 1 day at 4 °C with primary antibodies in blocking solution. For the first set of markers, GFAP (GA5) Mouse mAb (#3670, 1:50, Cell Signaling) for glial cells and Neurofilament-L (C28E10) Rabbit mAb (#2837, 1:100, Cell Signaling) for neurons were used. For the second set, Iba1/AIF-1 (E4O4W) XP® Rabbit mAb (#17198, 1:800, Cell Signaling) for activated microglia and NeuN (E4M5P) Mouse mAb (#94403, 1:100, Cell Signaling) for neurons were used.

Sections were then washed three times with PBS for 30 min and then stained for 2 h at 4 °C with corresponding secondary antibodies (Goat anti-Mouse IgG (H+L) Highly Cross-Adsorbed Secondary Antibody, Alexa Fluor Plus 488, A32723, 1:250, Thermo-Fisher; Goat anti-Rabbit IgG (H+L) Highly Cross-Adsorbed Secondary Antibody, Alexa Fluor Plus 647, A32733, 1:250, Thermo-Fisher). Slices were washed three times with PBS and incubated with DAPI (4′ 6-diamidino-2-phenylindole) (1:50,000) for 30 min. All fluorescent images were acquired with a laser scanning confocal microscope (Leica SP8) using a 10× air lens.

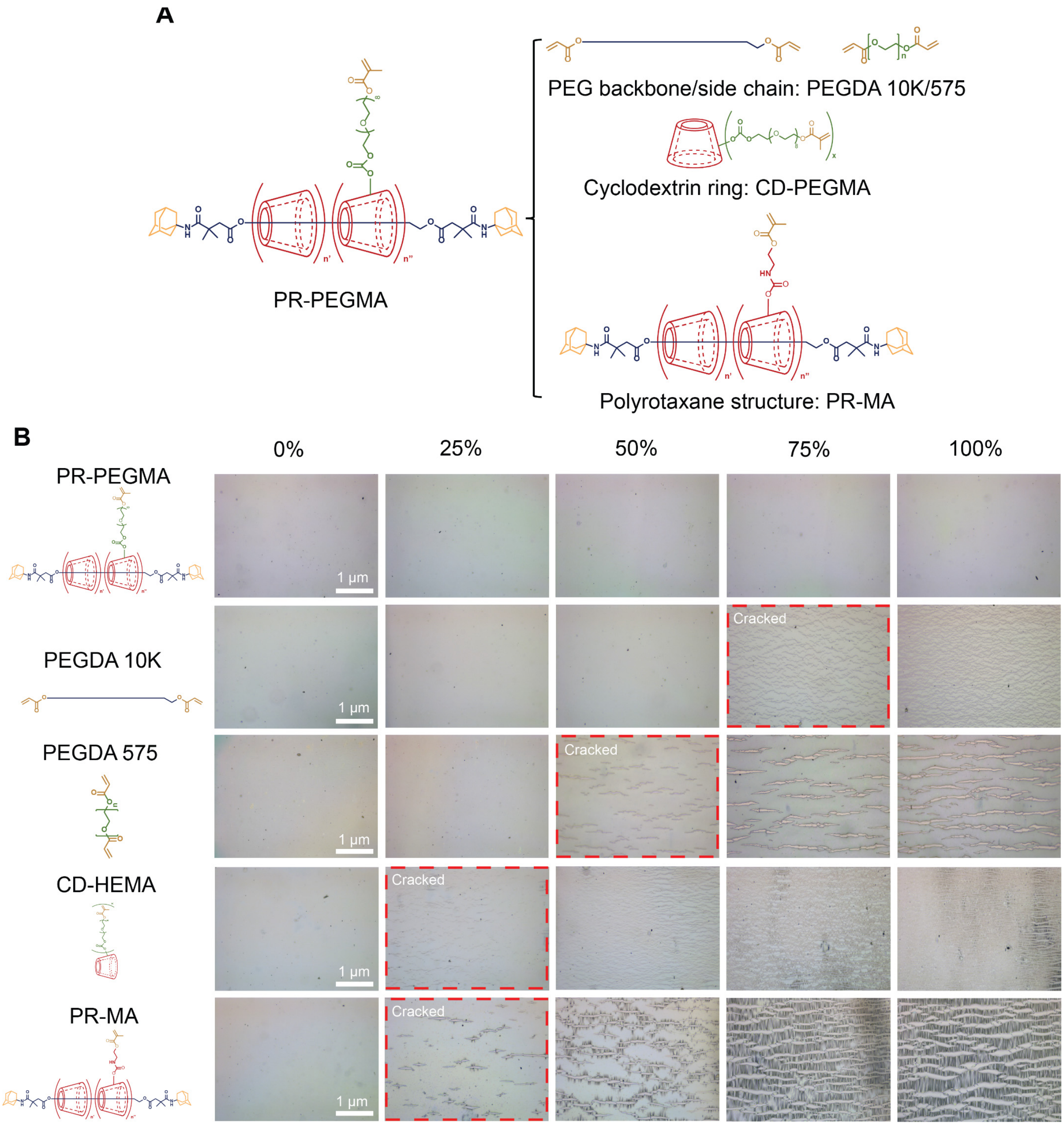

**Fig. S11. Stretching tests of PEDOT:PSS blended with PR-PEGMA and control samples mimicking individual building blocks. (A)** Chemical structures of samples for stretching tests. **(B)** Optical microscope images of different PEDOT:PSS films showing the crack formation under strain. PR-PEGMA/PEDOT:PSS hybrid has substantially higher stretchability than all control samples. In all cases, 55 mg of additives were added to 1 ml of PEDOT:PSS solution.

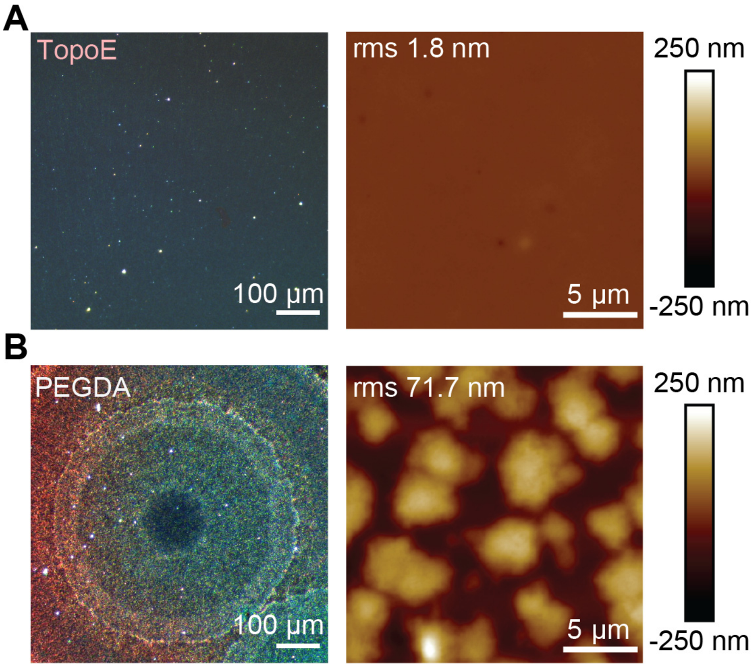

**Fig. S12. Dark field optical microscope (left column) and AFM (right column) images of TopoE (A) and PEGDA/PEDOT:PSS (B) films.** Due to the high degree of crystallization of PEGDA, the as-blended film had significantly higher surface roughness than the TopoE film.

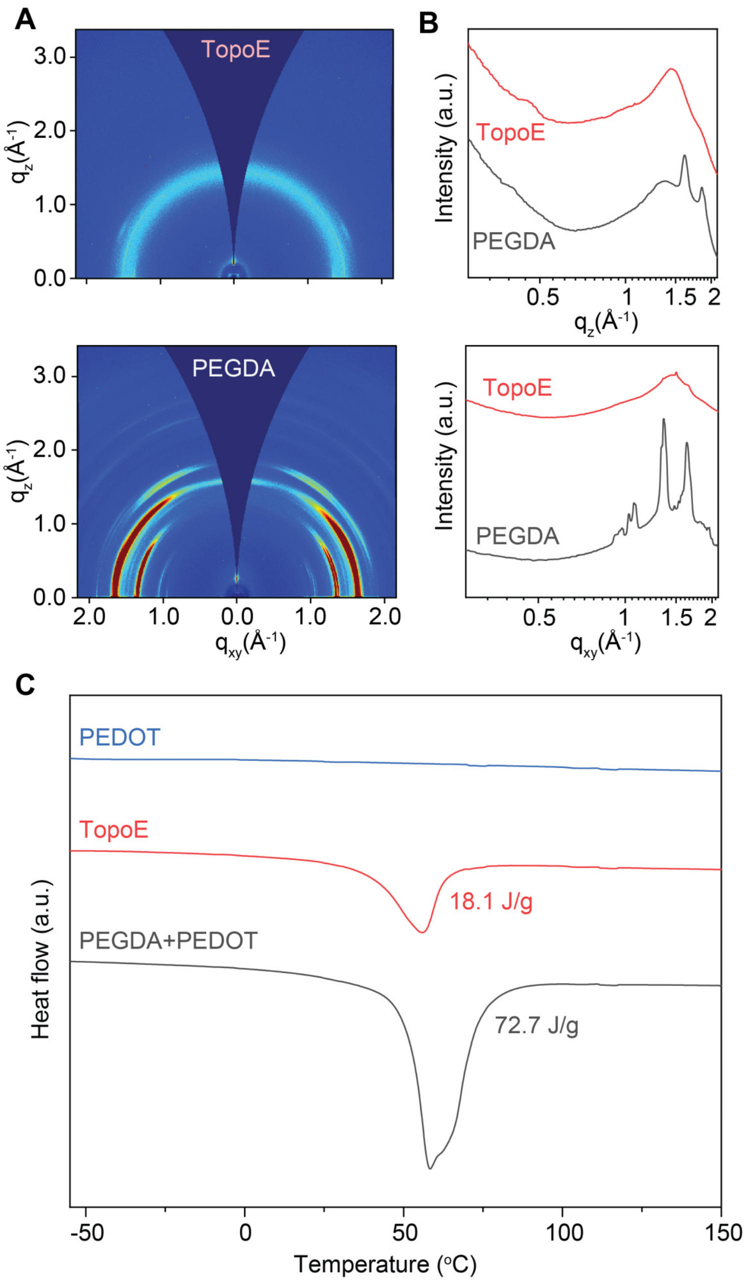

**Fig. S13. GIXD and DSC spectra of TopoE and PEGDA blended PEDOT:PSS films.** Compared to TopoE, the PEGDA film has significantly higher crystallinity as evidenced by the strong diffraction peaks (**A-B**) and heat of melting measured by DSC (**C**). The 1D scattering profile in (**B**) was calculated by taking the azimuthal average values along *z* (upper row) and *xy* (lower row) directions of the raw 2D image in (**A**).

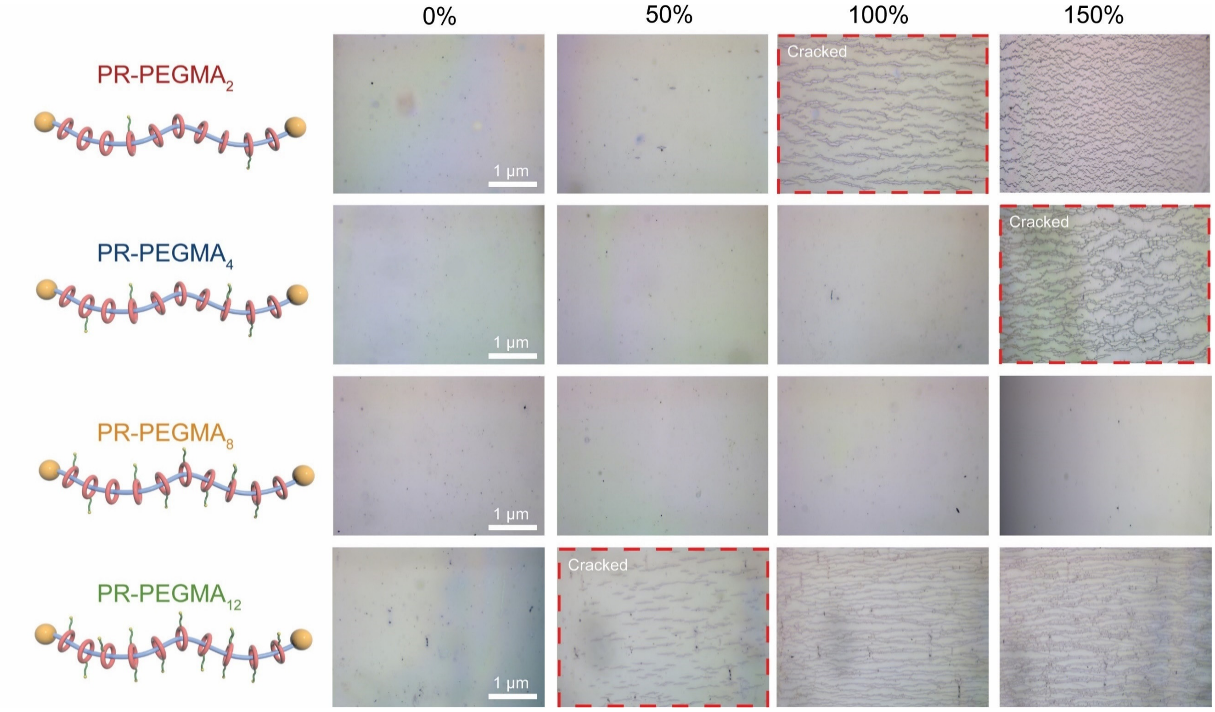

**Fig. S14. Stretching tests of TopoE with different number of side chains on PR-PEGMA.** Optical microscope images of different PEDOT:PSS films after photocrosslinking and water development showing the crack formation under strain. PR-PEGMA$_8$ has better stretchability than other samples. In all cases, 55 mg of PR-PEGMA were added to 1 ml of PEDOT:PSS solution.

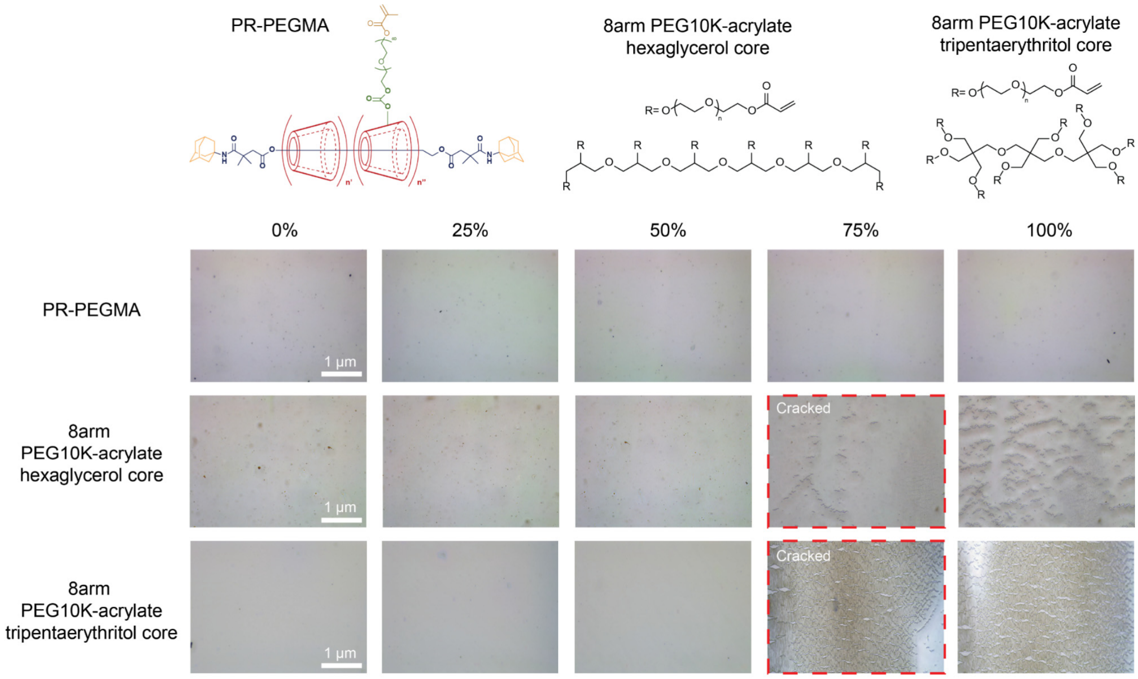

**Fig. S15. Stretching tests of TopoE and PEDOT:PSS blended with multi-arm PEGDA samples.** Optical microscope images of different PEDOT:PSS films showing the crack formation under strain. Multi-arm PEGDA samples cannot promote the film stretchability. In all cases, 55 mg of additives were added to 1 ml of PEDOT:PSS solution.

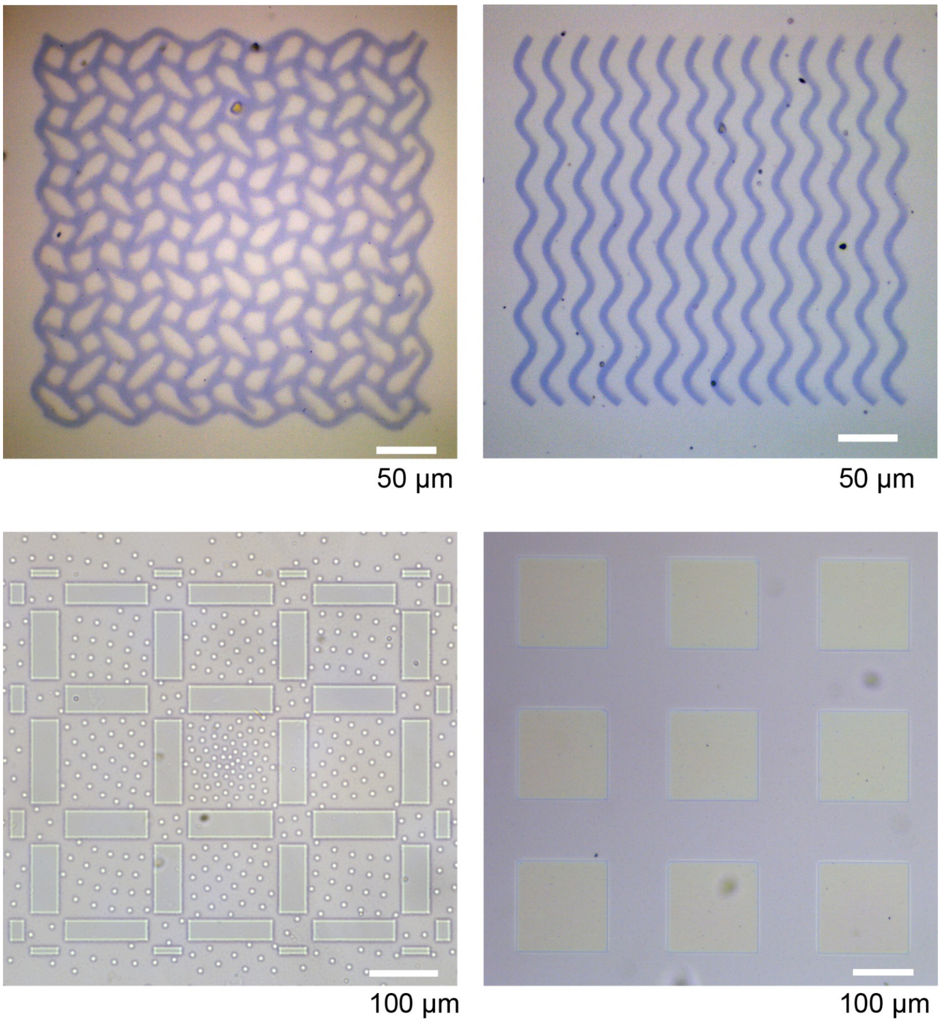

**Fig. S16. Optical microscope images of photopatterned PEDOT:PSS films.**

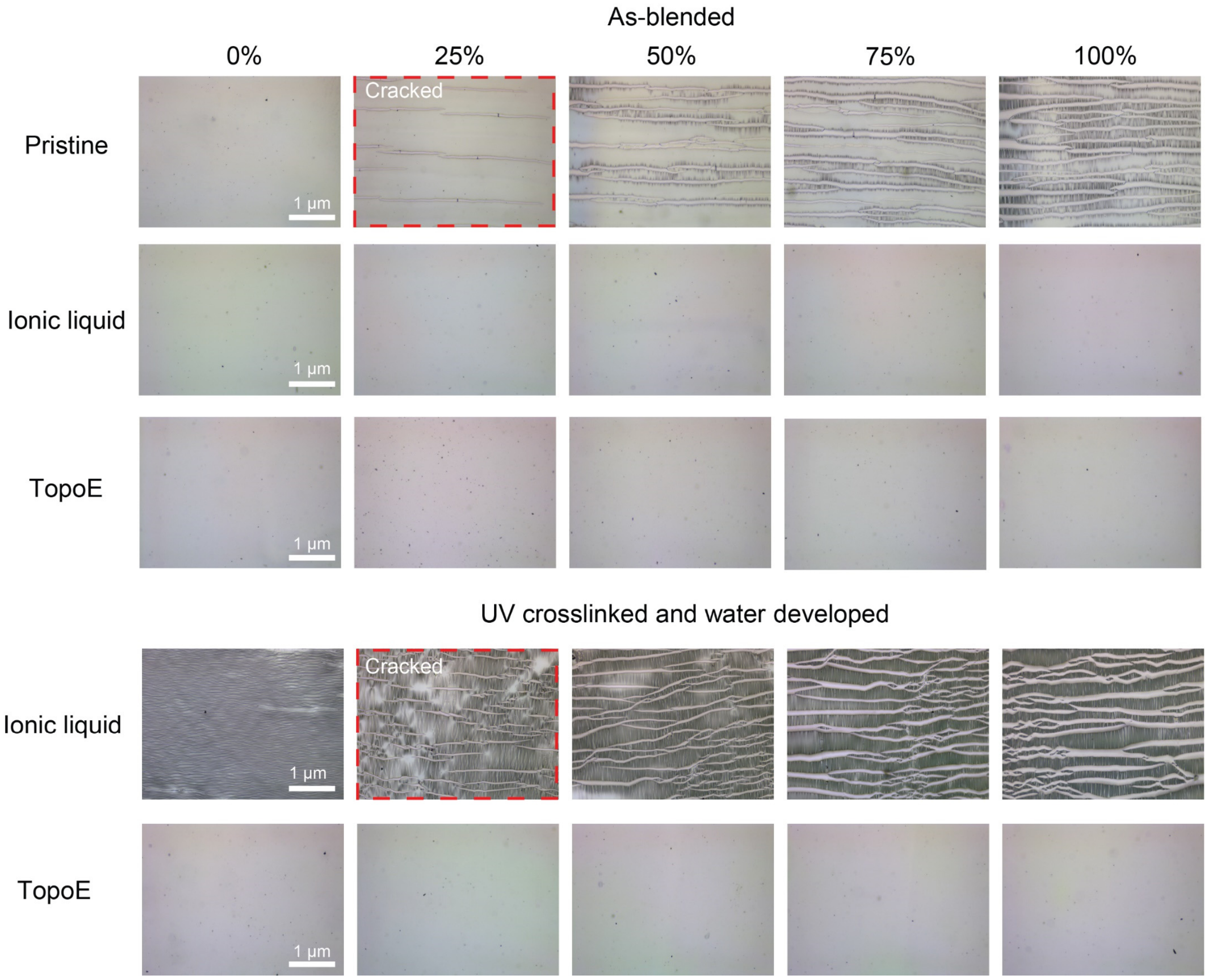

**Fig. S17. Stretching tests of TopoE and PEDOT:PSS blended with ionic liquid.** Optical microscope images of different PEDOT:PSS films showing the crack formation under strain. Ionic liquid can promote the stretchability of PEDOT:PSS right after blending. However, ionic liquid will leach out after water development and the entire film would completely lose its stretchability. On the other hand, crosslinked PR can be well preserved in the film for stable film stretchability after water development, which is essential for subsequent device fabrications and biological applications. 4-(3-butyl-1-imidazolio)-1-butanesulfonic acid triflate was chosen as the representative ionic liquid additive. In both cases, 55 mg of ionic liquid or TopoE was added to 1 mL of PEDOT:PSS solution.

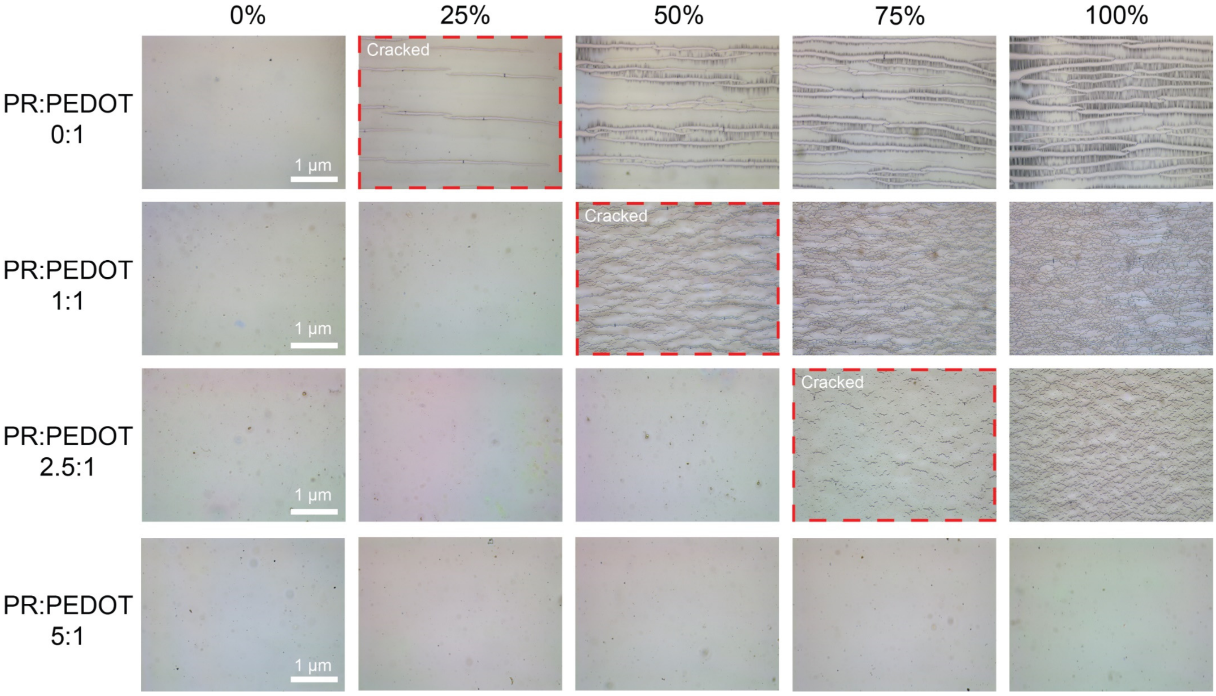

**Fig. S18. Stretching tests of TopoE with different PR:PEDOT ratios.** Optical microscope images of different TopoE films showing the crack formation under strain. The more PR added to the PEDOT:PSS solution, the higher the film stretchability. The weight ratio was calculated by the dry mass ratio between PR and PEDOT:PSS. Assuming approximately 11 mg of PEDOT:PSS per 1 mL of solution, 11 mg, 27.5 mg, and 55 mg of PR were added to yield PR:PEDOT weight ratios of 1:1, 2.5:1, and 5:1.

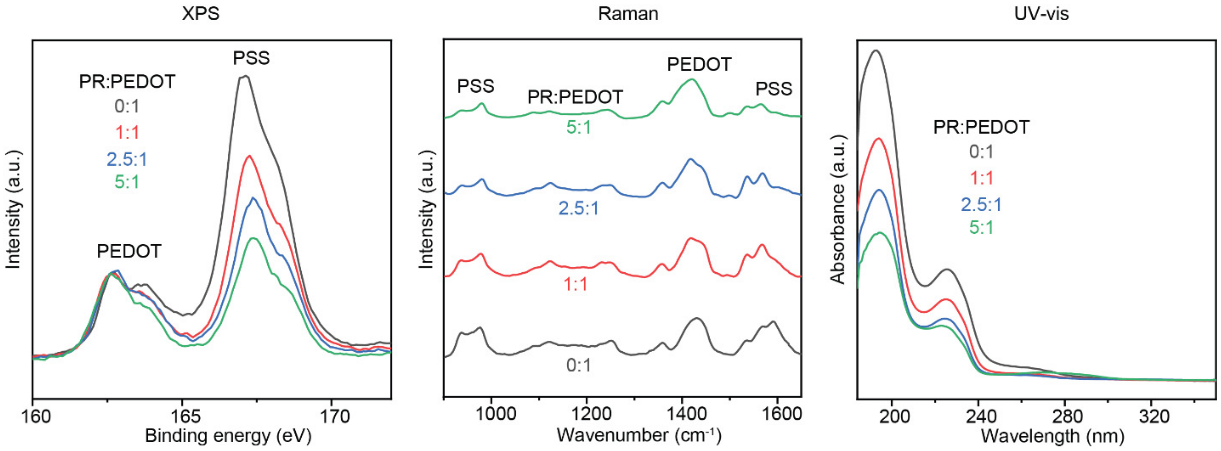

**Fig. S19. XPS, Raman, and UV-vis spectra of TopoE with different PR:PEDOT ratios.** All three measurements confirmed the same trend that PR would induce the removal of PSS from PEDOT. For the XPS spectra (left column), after normalizing the sulfur peaks (S 2p) of PEDOT thiophene (166-162 eV), the S 2p peak intensity of the PSS (170-166 eV) apparently decreased with respect to the PR:PEDOT ratio (Ref. S1). For the Raman spectra, more PR in the TopoE film led to decreased peak intensities of PSS at 1500-1650 $cm^{-1}$ and 900-1050 $cm^{-1}$ (Refs. S2,3). For the UV-vis spectra, the two peaks originated from the aromatic rings of PSS between 180 nm and 240 nm (Ref. S4) got significantly decreased with more PR in the TopoE film.

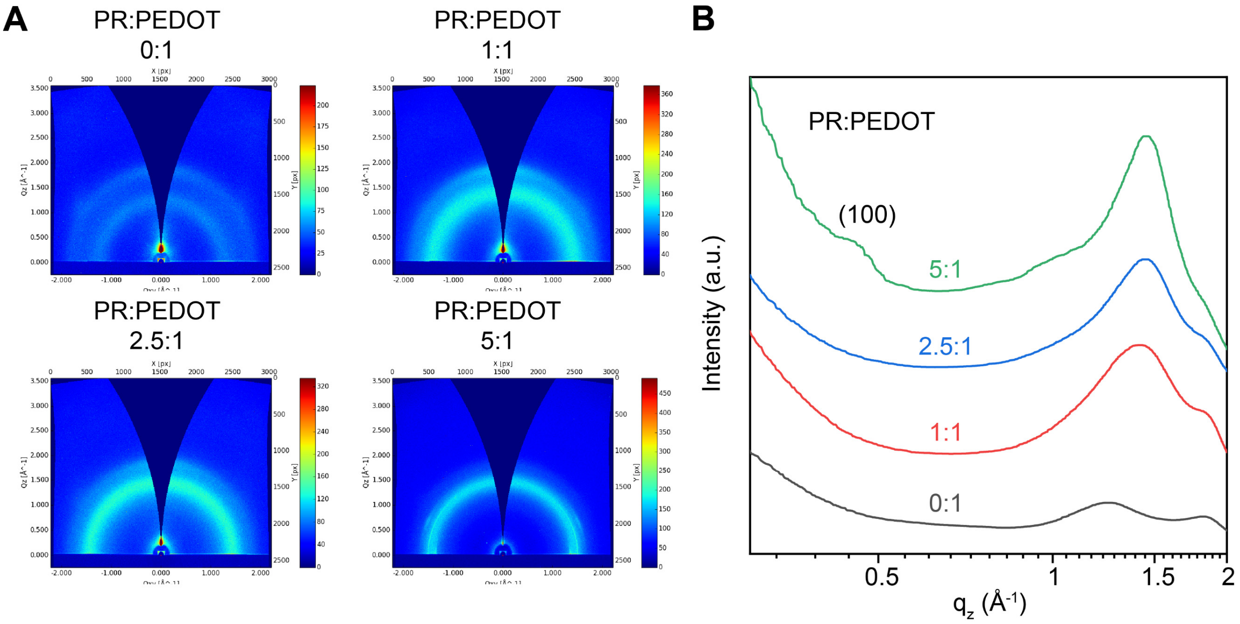

**Fig. S20. GIXD spectra of TopoE with different PR:PEDOT ratios.** The 1D scattering profile in (**B**) was calculated by taking the azimuthal average of the raw images (**A**) in all directions. With more PR in the TopoE film, the PEDOT lamellar packing (100) peak became pronounced, indicating enhanced PEDOT aggregation, which gave a higher conductivity.

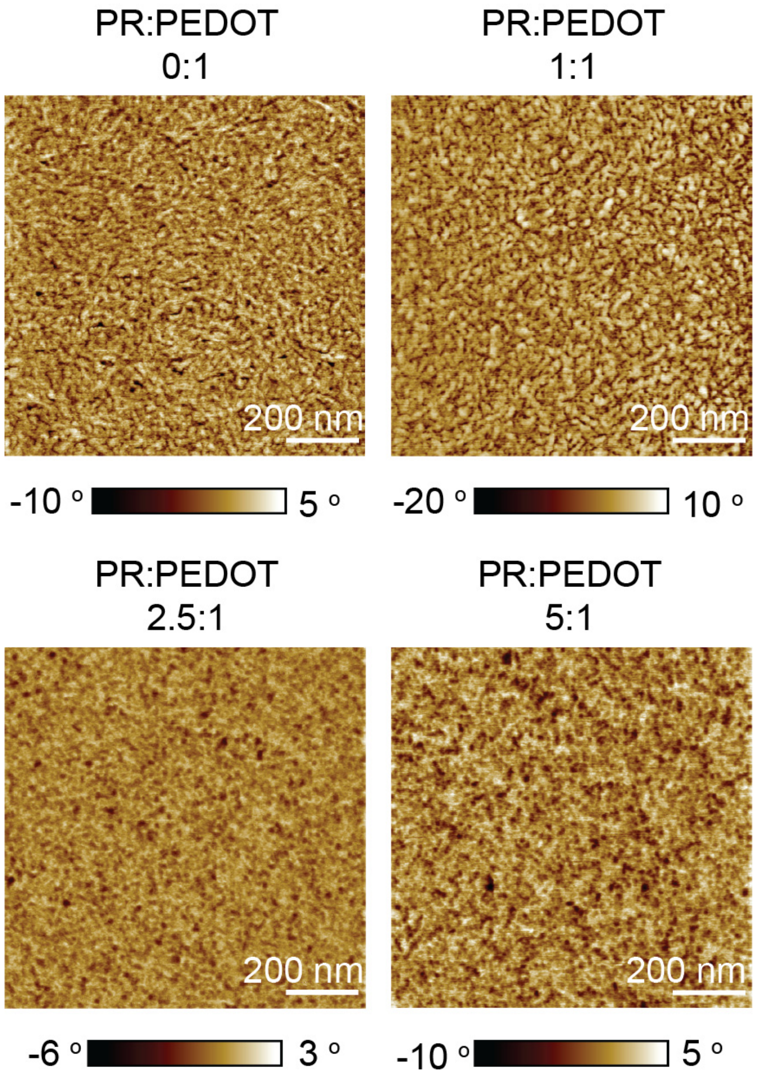

**Fig. S21. AFM phase images of TopoE with different PR:PEDOT ratios after photocrosslinking.** With higher concentration of PR in the PEDOT:PSS solution, more micro-fiber structures can be formed.

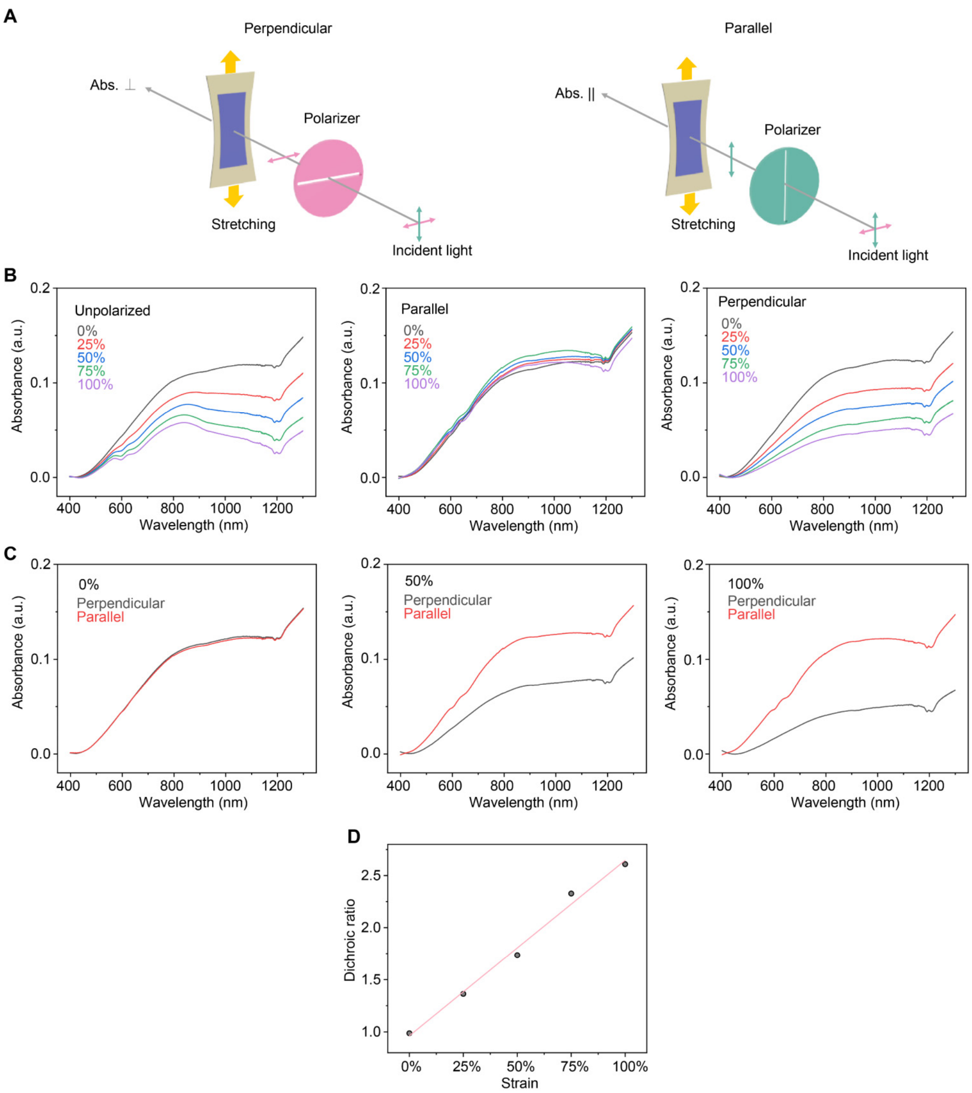

**Fig. S22. Polarized UV-vis spectra of TopoE films under strain (A-C) and the calculated dichroic ratio (D). (A)** Schematic diagrams illustrating the relative position of the incident light, the film stretching direction and the polarizer orientations. **(B)** Upon stretching, the film thickness of TopoE gradually decreases such that the total absorption after unpolarized light decreases with respect the strain levels (left column). Because of the chain alignment of PEDOT:PSS microfibers along the stretching direction, the absorption of the polarized light parallel to the stretching direction remains roughly the same despite the reduced film thickness (middle column) whereas the absorption perpendicular to the stretching direction decreased substantially at higher strain (right column). **(C-D)** Comparison of light absorption parallel and perpendicular to the stretching direction at different strains (**C**) and their ratios (**D**). The dichroic ratio ($A_{\parallel}/A_{\perp}$) increased almost linearly with respect to the strain suggesting the microfibers were highly aligned without cracking. The dichroic ratio was calculated using the absorption at 810 nm, corresponding with de-doped PEDOT. The films were prepared by spin coating of PR/PEDOT:PSS (5/1) mixture on SEBS substrates with an initial thickness of ~200 nm.

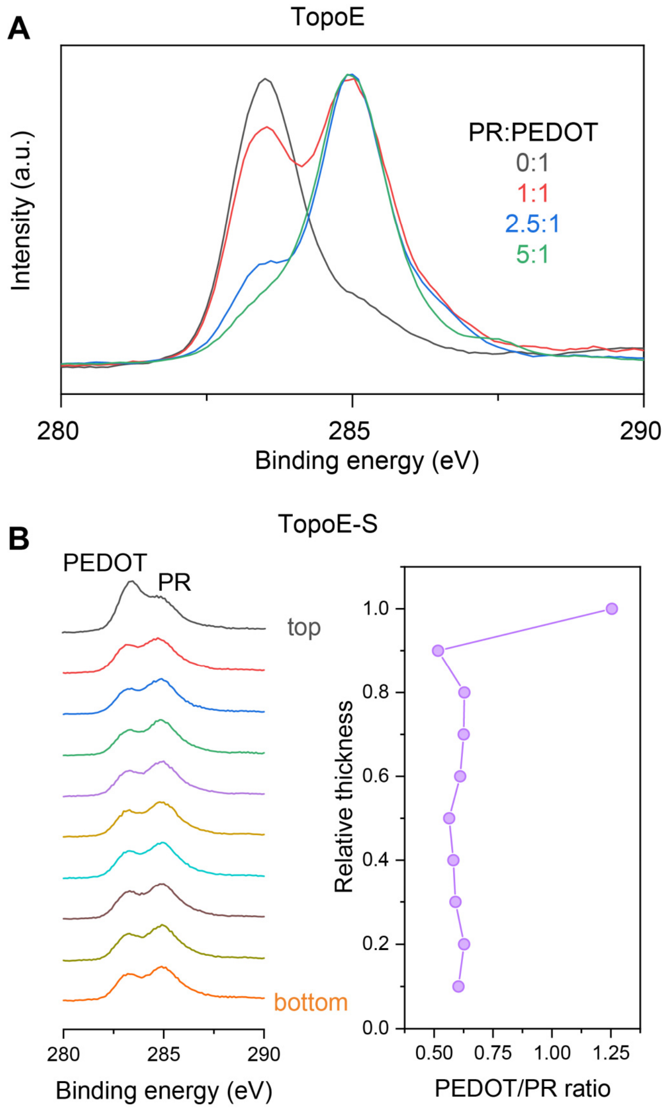

**Fig. S23. XPS of the TopoE film before (A) and after (B) acid treatment.** (**A**) XPS spectra of TopoE after blending and photocrosslinking showing the relative intensity changes corresponding to the conjugated PEDOT:PSS (283.5 eV, C 1s $sp^2$) and PR (284.9 eV, C 1s $sp^3$). (**B**) Depth profile of the acid treated film indicating more PEDOT at the surface (right column). The PEDOT/PR ratio was calculated by dividing the integrated areas of the deconvoluted $sp^2$ and $sp^3$ carbon peaks. After acid treatment the PR-PEGMA remains in the TopoE-S as indicated by the presence of the PR peak at 284.9 eV, which is essential to maintain the stretchability of the film.

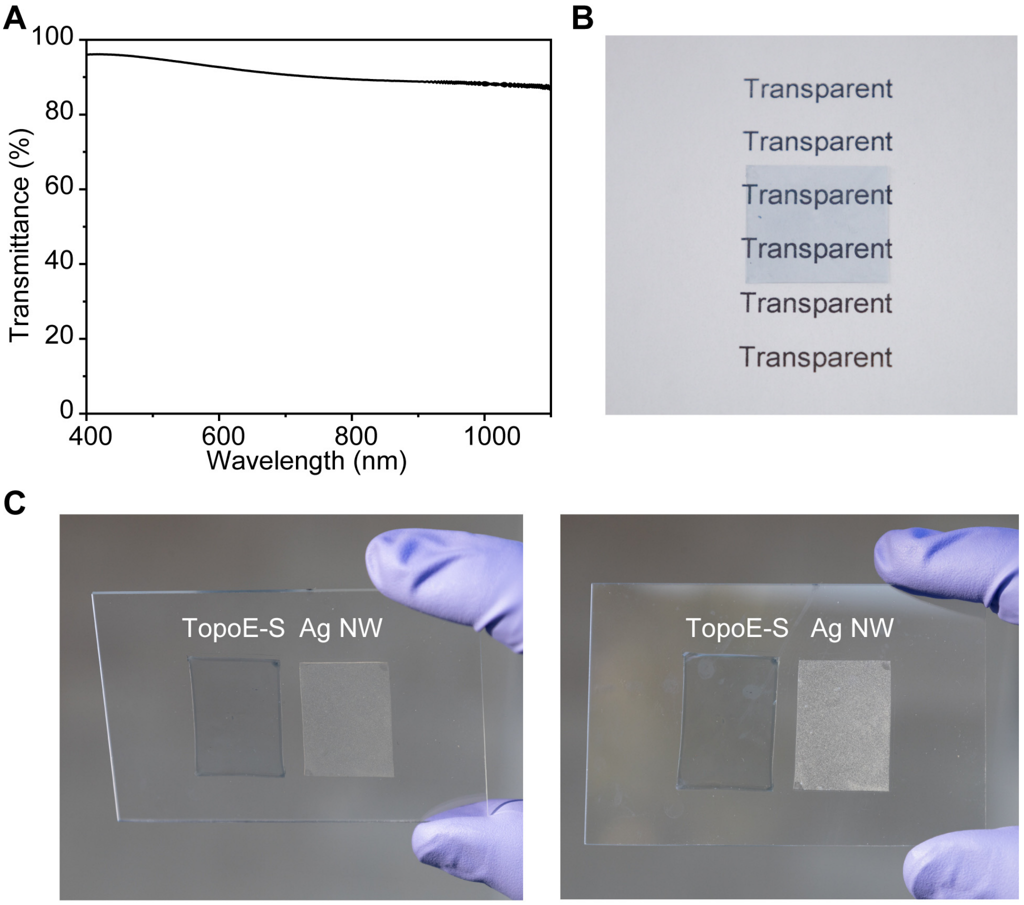

D

| Material | Transmittance | Sheet resistance | Reference |
|---|---|---|---|
| Carbon nanotube (CNT) | 88% | 500 Ω $sq^{-1}$ | 5 |
| Indium tin oxide (ITO) | 90.2% | 30 Ω $sq^{-1}$ | 6 |
| Silver nanowire (Ag NW) | 80% | 20 Ω $sq^{-1}$ | 7 |
| Non-stretchable PEDOT:PSS | 90% | 46.1 Ω $sq^{-1}$ | 8 |
| TopoE-S | 94% | 69 Ω sq-1 | This work |

**Fig. S24. The TopoE-S film is optically transparent in the visible range. (A)** UV-vis spectrum of the TopoE-S film. The transmittance value at 550 nm is 94% with the sheet resistance of 69 Ω $sq^{-1}$. **(B)** Photo of a TopoE-S film demonstrating its transparency. The film was prepared by spin coating the PR/PEDOT:PSS (5/1) mixture onto a SEBS substrate followed by sulfuric acid treatment. **(C)** Photos of TopoE-S and silver nanowire coated SEBS substrates showing the hazy characteristic of Ag NW as a transparent conductor. **(D)** Comparison of TopoE-S versus common transparent conductors including carbon nanotube (CNT)[5], indium tin oxide (ITO)[6], silver nanowire (Ag NW)[7], and non-stretchable PEDOT:PSS[8].

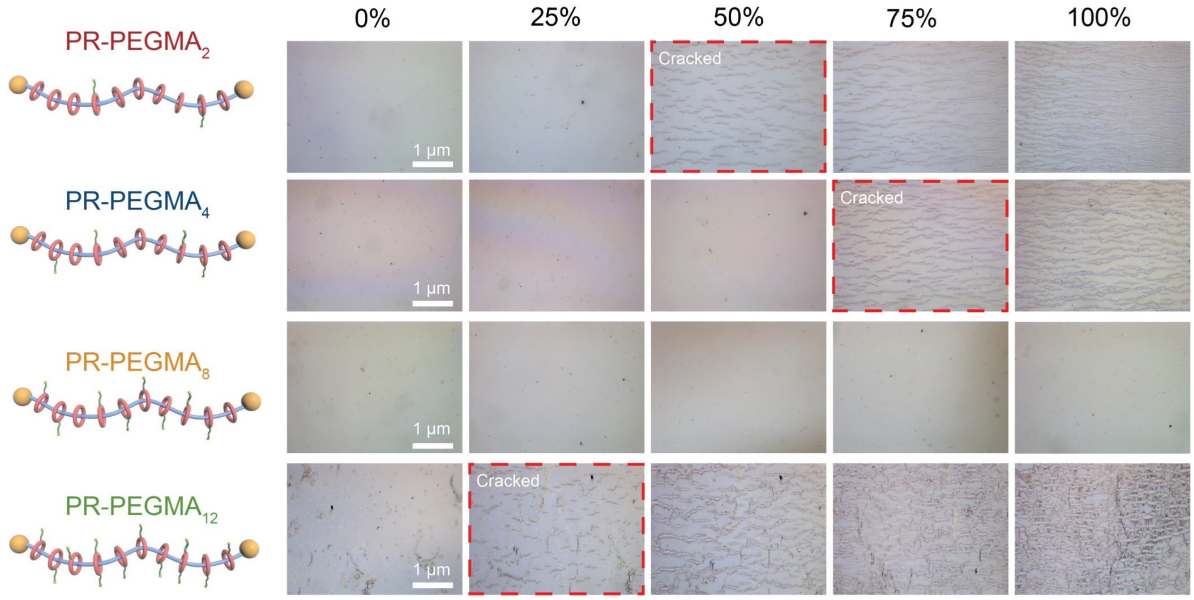

**Fig. S25. Stretching tests of TopoE-S with different number of side chains on PR.** Optical microscope images of different TopoE-S films showing the crack formation under strain. PR-PEGMA$_8$ has better stretchability than other samples even after acid treatment. All films were prepared by spin coating the PR/PEDOT:PSS (5/1) mixture onto SEBS substrates followed by concentrated sulfuric acid treatment. The final film thickness was around 100 nm.

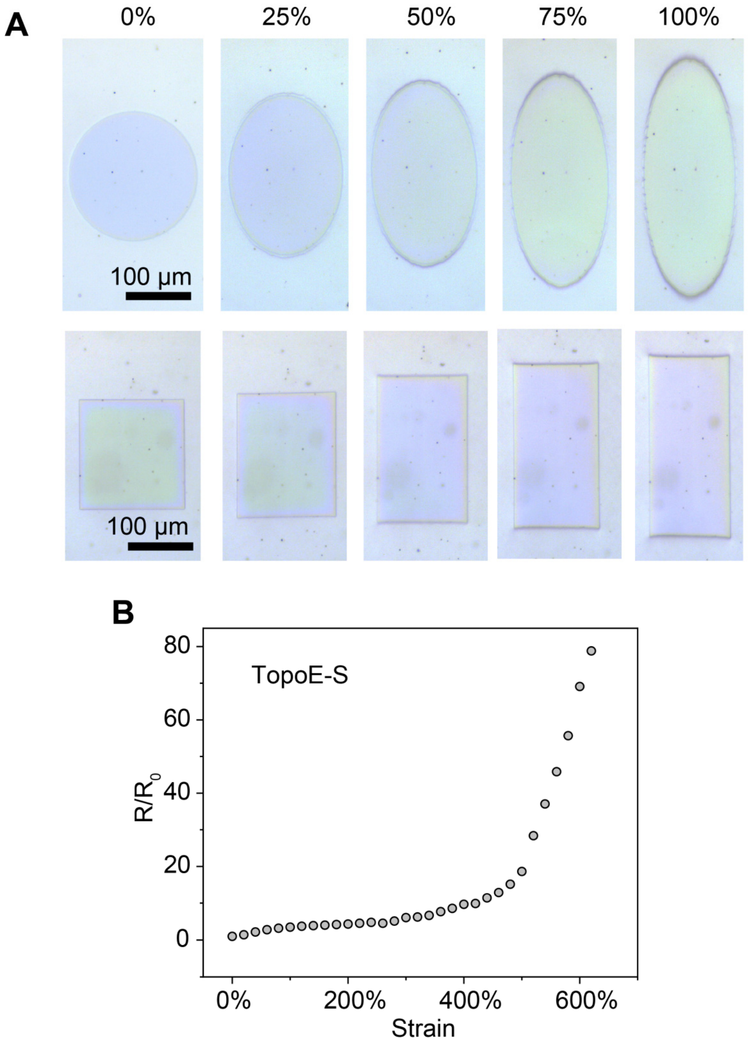

**Fig. S26. Optical microscope images and resistance measurements of TopoE-S under strain.** After acid treatment, the crosslinked TopoE-S composite can be stretched to 100% without any cracks (**A**) with the ultimate strain up to ~600% (**B**). All films were prepared by spin coating the PR/PEDOT:PSS (5/1) mixture onto SEBS substrates followed by concentrated sulfuric acid treatment. The final film thickness was around 100 nm.

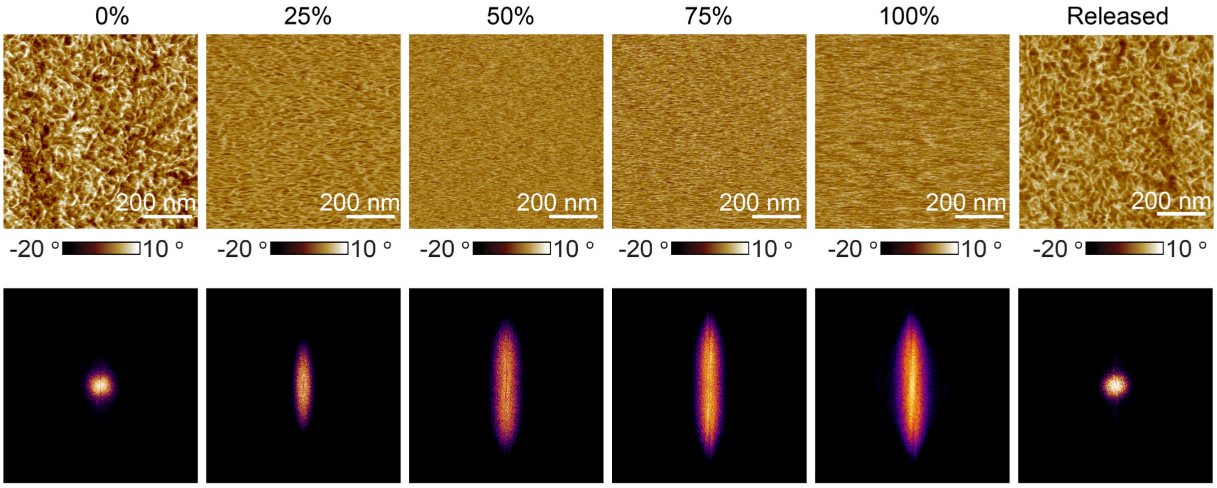

**Fig. S27. AFM phase images (upper row) and corresponding FFT diffractograms (lower row) of TopoE-S under strain.** Upon stretching, PEDOT:PSS micro-fibers after acid treatment will be aligned along the strain axis. The fibers will return to its original isotropic configuration after releasing.

A

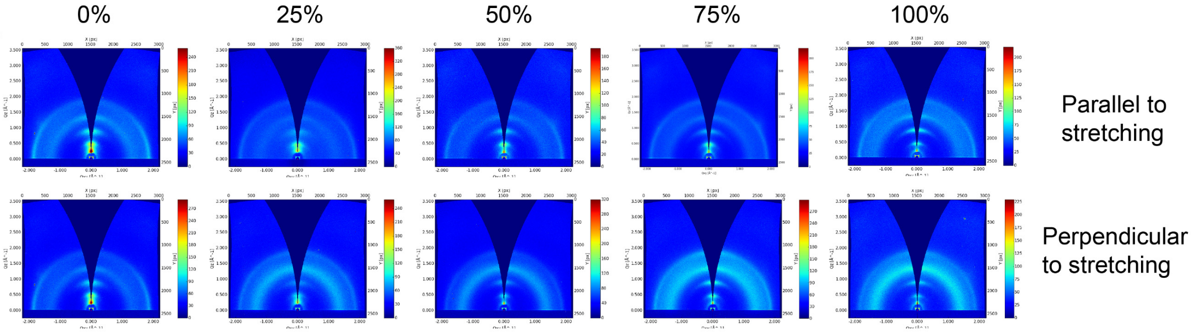

B

| Strain | Parallel (100) Location | Parallel (100) FWHM | Perpendicular (100) Location | Perpendicular (100) FWHM |
|---|---|---|---|---|
| 0% | 0.4995 | 0.0953 | 0.4970 | 0.0864 |
| 25% | 0.4767 | 0.0876 | 0.4932 | 0.1015 |
| 50% | 0.4823 | 0.1090 | 0.4785 | 0.1162 |
| 75% | 0.4774 | 0.1080 | 0.4702 | 0.0984 |
| 100% | 0.4702 | 0.0983 | 0.4707 | 0.0905 |

**Fig. S28. GIXD spectra of TopoE-S films under strain.** PEDOT packings are preserved in directions parallel and perpendicular to the stretching direction as evidenced by the marginal changes of peak position and full width half maximum (FWHM) of the (100) lamellar packing over different strain levels. The film was prepared by spin coating the PR/PEDOT:PSS (5/1) mixture onto an octadecyltrimethoxysilane (OTS) treated silicon wafer. The film was then transferred onto SEBS substrates followed by stretching and acid treatment before being transferred back onto a bare silicon wafer before GIXD measurements. The initial film thickness was around 100 nm.

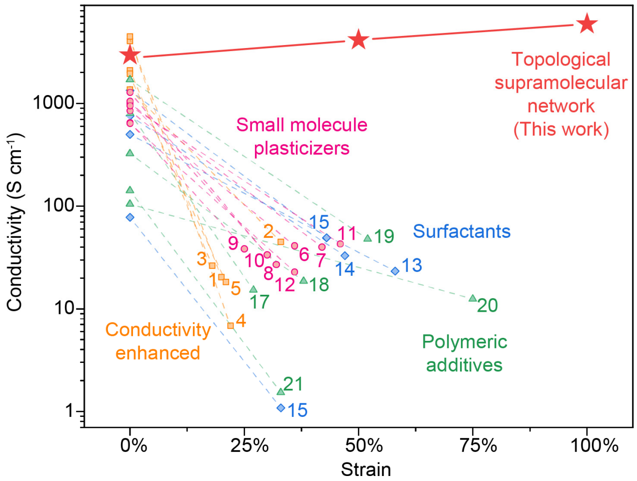

| | Index | Chemical name/ Treatment method | Chemical structure | Initial conductivity | Conductivity under strain | Reference |
|---|---|---|---|---|---|---|
| Conductivity enhanced | 1 | Sulfuric acid | $H_2SO_4$ | 4380 S cm$^{-1}$ | 20.4 S cm$^{-1}$@20% | *Adv. Mater.* **26**, 2268–2272 (2014) |
| | 2 | Methanol | $CH_3OH$ | 1362 S cm$^{-1}$ | 44.9 S cm$^{-1}$@33% | *Energy Environ. Sci.* **5**, 9662-9671 (2012) |
| | 3 | Nitric acid | $HNO_3$ | 2100 S cm$^{-1}$ | 26.4 S cm$^{-1}$@18% | *Adv. Electron. Mater.* **5**, 1800654 (2019) |
| | 4 | Shearing | N/A | 4600 S cm$^{-1}$ | 6.79 S cm$^{-1}$@22% | *Proc. Natl. Acad. Sci.* **112** 14138 (2015) |
| | 5 | Methylammonium iodide (MAI) Dimethylformamide (DMF) | $CH_3NH_3^+I^-$ / $HCON(CH_3)_2$ | 2100 Scm$^{-1}$ | 18.3 S cm$^{-1}$@21% | *ACS Appl. Mater. Interfaces* **8**, 11629–11638 (2016) |
| Small molecule plasticizers | 6 | Glycerol | 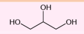 | 850 S cm$^{-1}$ | 41.3 S cm$^{-1}$@36% | *Macromolecules* 54, 1234–1242 (2021) |
| | 7 | Sorbitol | 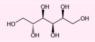 | 1000 S cm$^{-1}$ | 40.1 S cm$^{-1}$@42% | *ACS Appl. Mater. Interfaces* **11**, 26185-26193 (2019) |
| | 8 | 4-(3-Butyl-1-imidazolio)-1-butanesulfonic acid triflate | 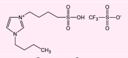 | 1050 S cm$^{-1}$ | 27.1 S cm$^{-1}$@32% | *Sci. Adv.* **3**, e1602076 (2017) |
| | 9 | Lithium bis(trifluoromethanesulfonyl) imide (LiTFSI) | 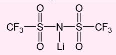 | 652 S cm$^{-1}$ | 38.5 S cm$^{-1}$@25% | *Sci. Adv.* **3**, e1602076 (2017) |
| | 10 | 1-Ethyl-3-methylimidazolium tetracyanoborate (EMIM TCB) | 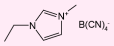 | 1280 S cm$^{-1}$ | 33.7 S cm$^{-1}$@30% | *ACS Applied Materials & Interfaces* **9**, 819–826 (2017) |
| | 11 | Dimethyl sulfoxide (DMSO) | $(CH_3)_2SO$ | 950 S cm$^{-1}$ | 43.1 S cm$^{-1}$@46% | *Macromolecules* 54, 1234–1242 (2021) |
| | 12 | Ethylene glycol (EG) | $HOCH_2CH_2OH$ | 640 S cm$^{-1}$ | 23.0 S cm$^{-1}$@36% | *J. Mater. Chem. A* **1**, 9907–9915 (2013) |
| Surfactants | 13 | 4-Dodecylbenzenesulfonic acid (DBSA) | 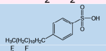 | 500 S cm$^{-1}$ | 23.4 S cm$^{-1}$@58% | *APL Materials* **3**, 014911 (2015) |
| | 14 | Zonyl FS-30 | 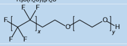 | 755 S cm$^{-1}$ | 33.0 S cm$^{-1}$@47% | *Adv. Funct. Mater.* **22**, 421–428 (2012) |
| | 15 | Triton X-100 | 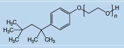 | 78 S cm$^{-1}$ | 1.1 S cm$^{-1}$@33% | *Adv. Mater.* **28**, 4455–4461 (2016) |
| | 16 | Sodium dodecyl sulfate (SDS) | $CH_3(CH_2)_{10}CH_2SO_3Na$ | 1335 S cm$^{-1}$ | 49.4 S cm$^{-1}$@43% | *RSC Adv.* **7**, 5888–5897 (2017) |
| Polymeric additives | 17 | Poly(ethylene glycol) 400 | 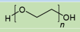 | 805 S cm$^{-1}$ | 15.3 S cm$^{-1}$@27% | *J. Mater. Chem. A* **1**, 9907–9915 (2013) |
| | 18 | Poly(ethylene glycol) 1000 | 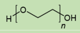 | 325 S cm$^{-1}$ | 18.7 S cm$^{-1}$@38% | *J. Mater. Chem. A* **1**, 9907–9915 (2013) |
| | 19 | Poly(ethylene glycol)-block-poly(propylene glycol)-block-poly(ethylene glycol) | 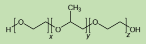 | 1700 S cm$^{-1}$ | 47.9 S cm$^{-1}$@32% | *ACS Appl. Mater. Interfaces* **10**, 28027-28035 (2018) |
| | 20 | Waterborne polyurethane | 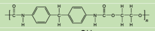 | 105 S cm$^{-1}$ | 12.5 S cm$^{-1}$@75% | *Adv. Mater.* **26**, 3451–3458 (2014) |
| | 21 | Poly(vinyl alcohol) | 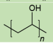 | 142 S cm$^{-1}$ | 1.5 S cm$^{-1}$@33% | *ACS Appl. Mater. Interfaces* **7**, 18415–18423 (2015) |

**Fig. S29. Comparison of conductivity and stretchability of TopoE-S versus literature reported strategies.** After solvent treatment during device fabrication and immersion in the physiological environment, previously reported PEDOT:PSS systems with post-treatment, small-molecule plasticizers, surfactants, polymeric additive were either not stretchable or have low conductivity, whereas our crosslinked topological network strategy can maintain all of its high performance with record breaking values, i.e., ~2700 S cm$^{-1}$ initial conductivity; ~6000 S cm$^{-1}$ at 100% strain along the stretching direction. All samples were prepared on SEBS substrates according to reported procedures and rigorously washed by water for 5 min before blow drying and measurements.

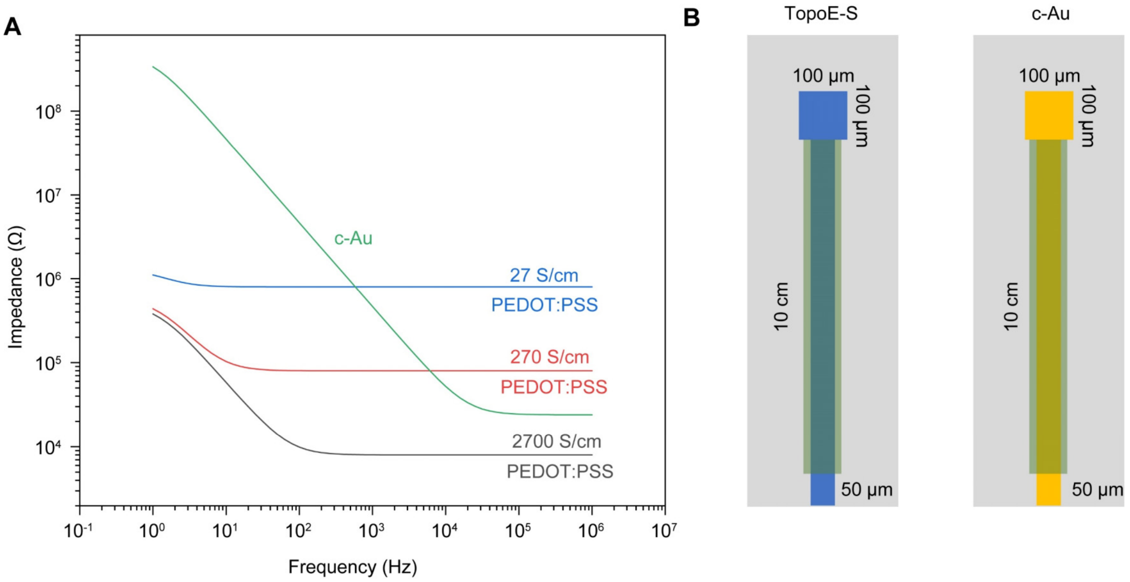

**Fig. S30. Simulated electrochemical impedance spectra of stretchable Au and PEDOT:PSS with different conductivities.** The equivalent circuit for PEDOT:PSS was a transmission line model adopted from Ref. S9 with the interfacial capacitance and resistance values taken from EIS measurements for TopoE-S. The model for cracked Au (c-Au) was a Randles circuit model with the sheet resistance value taken from Ref. S10, interfacial capacitance and resistance values from EIS measurements. Only PEDOT with a dry state conductivity value of 2700 S $cm^{-1}$ can consistently yield lower impedance across the entire frequency spectrum compared to c-Au.

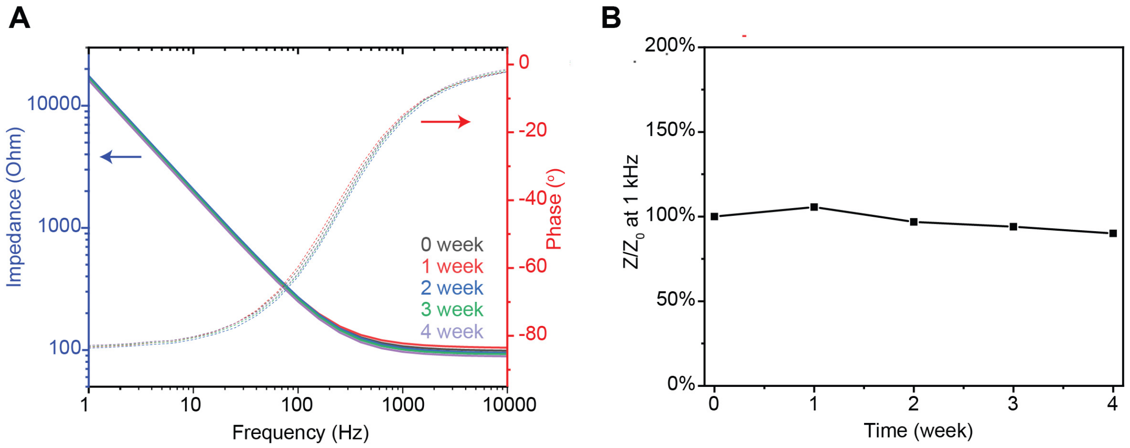

**Fig. S31. Electrochemical impedance spectrum of the TopoE-S electrode after immersion in PBS for four weeks.** The impedance value of TopoE-S can be well maintained over time. The film was prepared by spin coating the PR/PEDOT:PSS (5/1) mixture onto glass substrates followed by acid treatment. The final film thickness was around 100 nm.

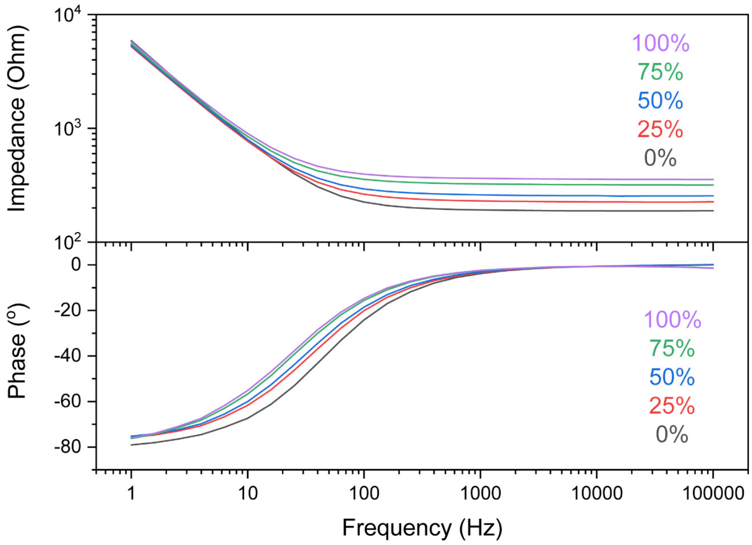

**Fig. S32. EIS spectrum of the TopoE-S electrode under strain.** The impedance value of PEDOT:PSS show only minor increases due to the geometrical changes under strain.

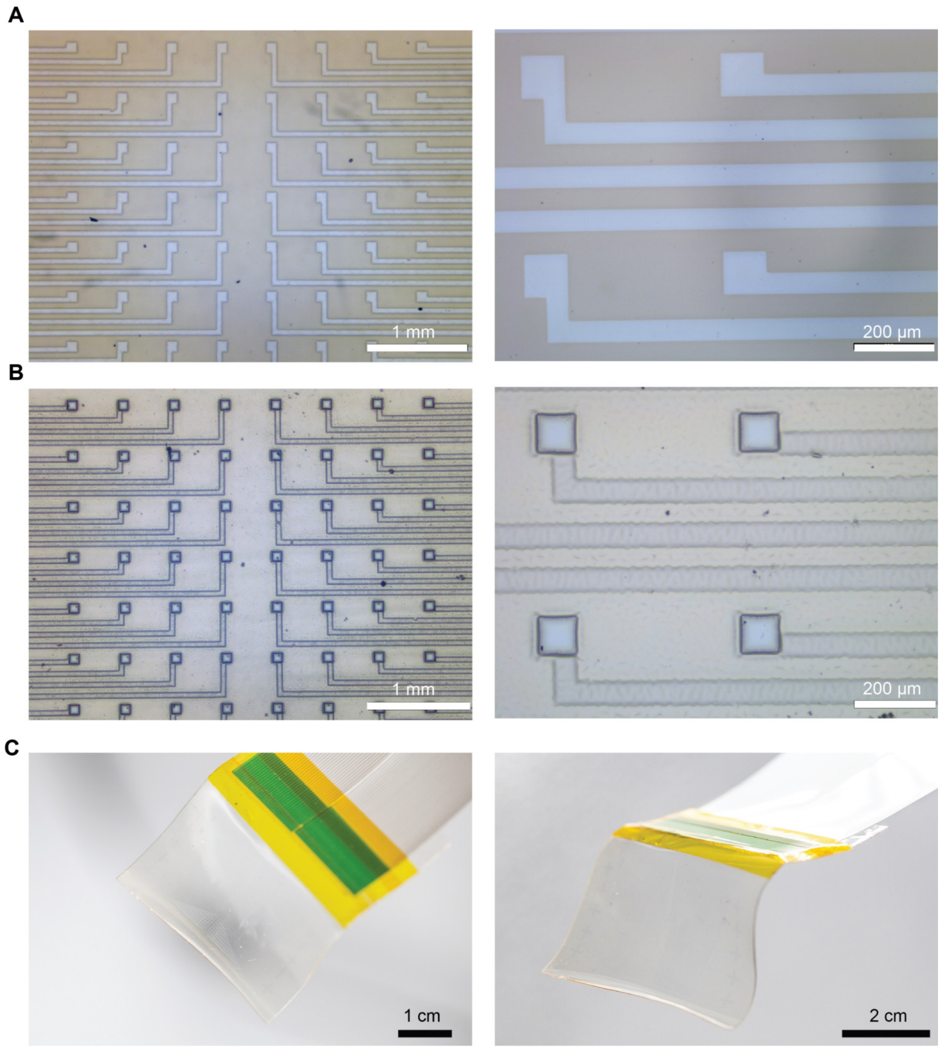

**Fig. S33. Images of the PEDOT:PSS electrode array. (A-B)** Optical microscope images of photopatterned TopoE-S electrode array before (**A**) and after (**B**) encapsulation. **(C)** Photographic images of the as-fabricated device.

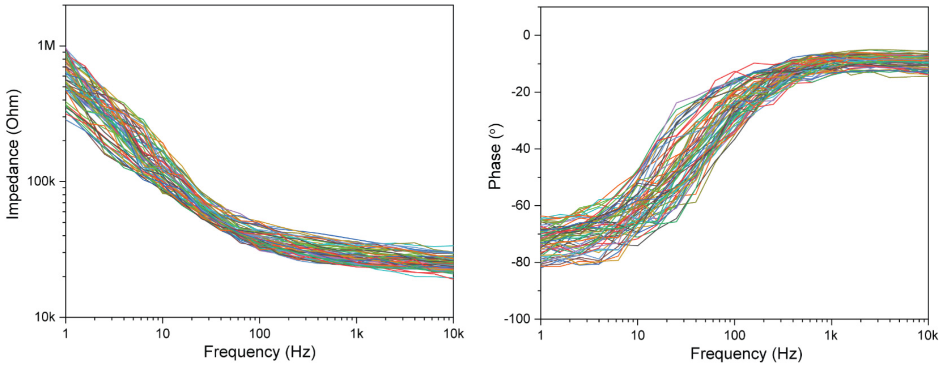

**Fig. S34. EIS data collected from a 64-channel electrode array showing the low device variation after the scalable fabrication process.**

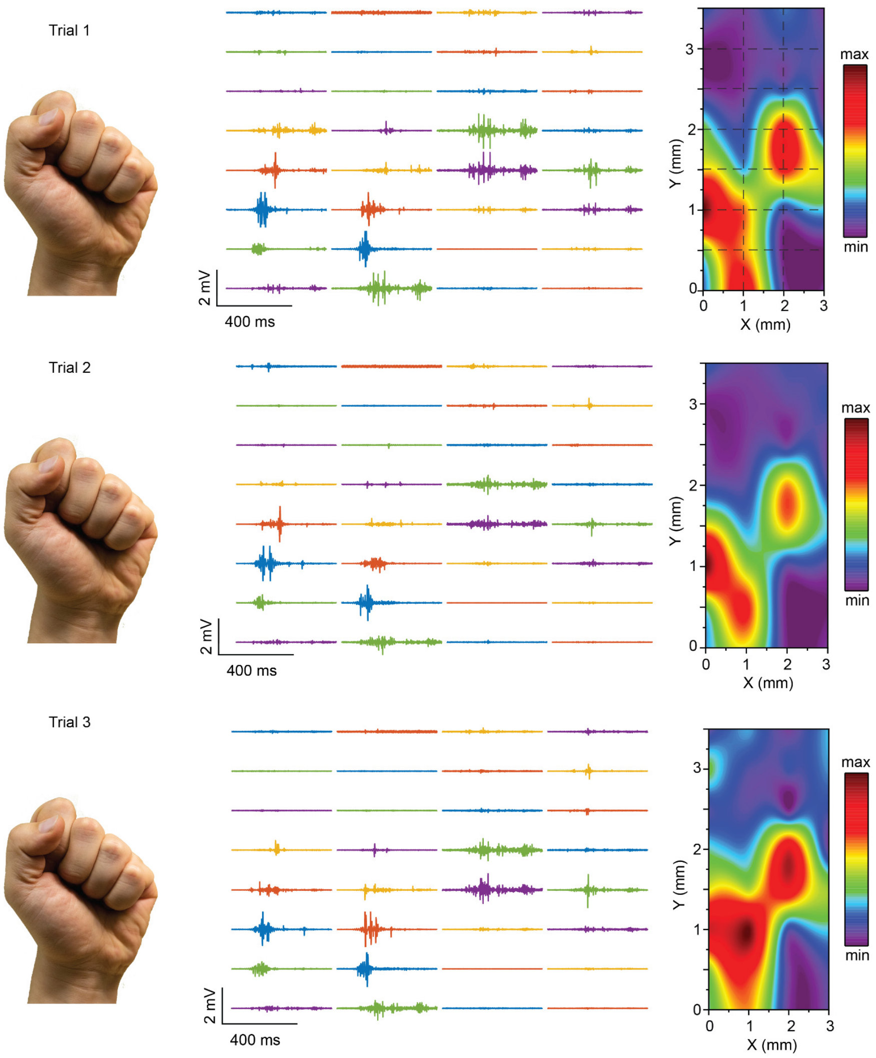

**Fig. S35. Three representative EMG data and calculated heat maps of gesture fist showing the high reproducibility of the stretchable electrode array for sEMG measurement.** During the experiment, the stretchable array was placed at the same muscle location to record the gesture for three separate times. The location of the electrode array is shown in **Fig. 4C**.

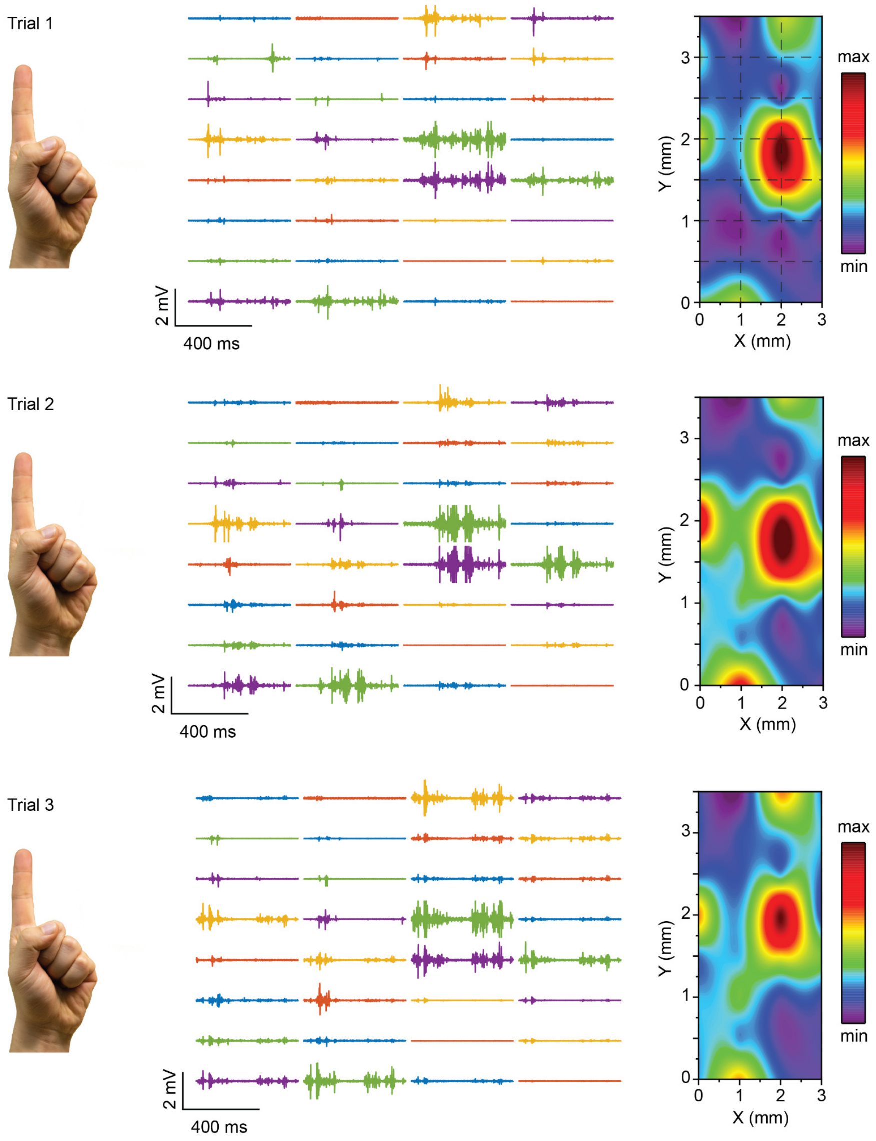

**Fig. S36. Three representative EMG data and calculated heat maps of gesture number one showing the high reproducibility of the stretchable electrode array for sEMG measurement.** During the experiment, the stretchable array was placed at the same muscle location to record the gesture for three separate times. The location of the electrode array is shown in **Fig. 4C**.

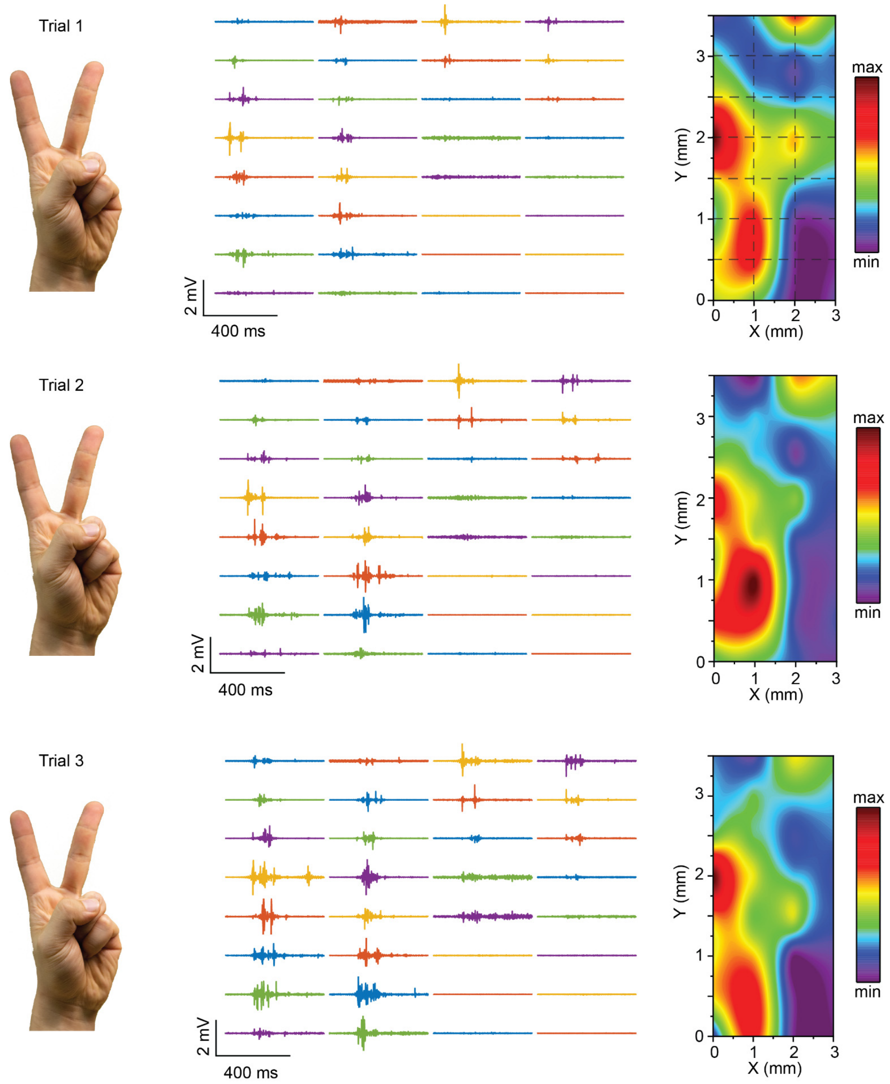

**Fig. S37. Three representative EMG data and calculated heat maps of gesture number two showing the high reproducibility of the stretchable electrode array for sEMG measurement.** During the experiment, the stretchable array was placed at the same muscle location to record the gesture for three separate times. The location of the electrode array is shown in **Fig. 4C**.

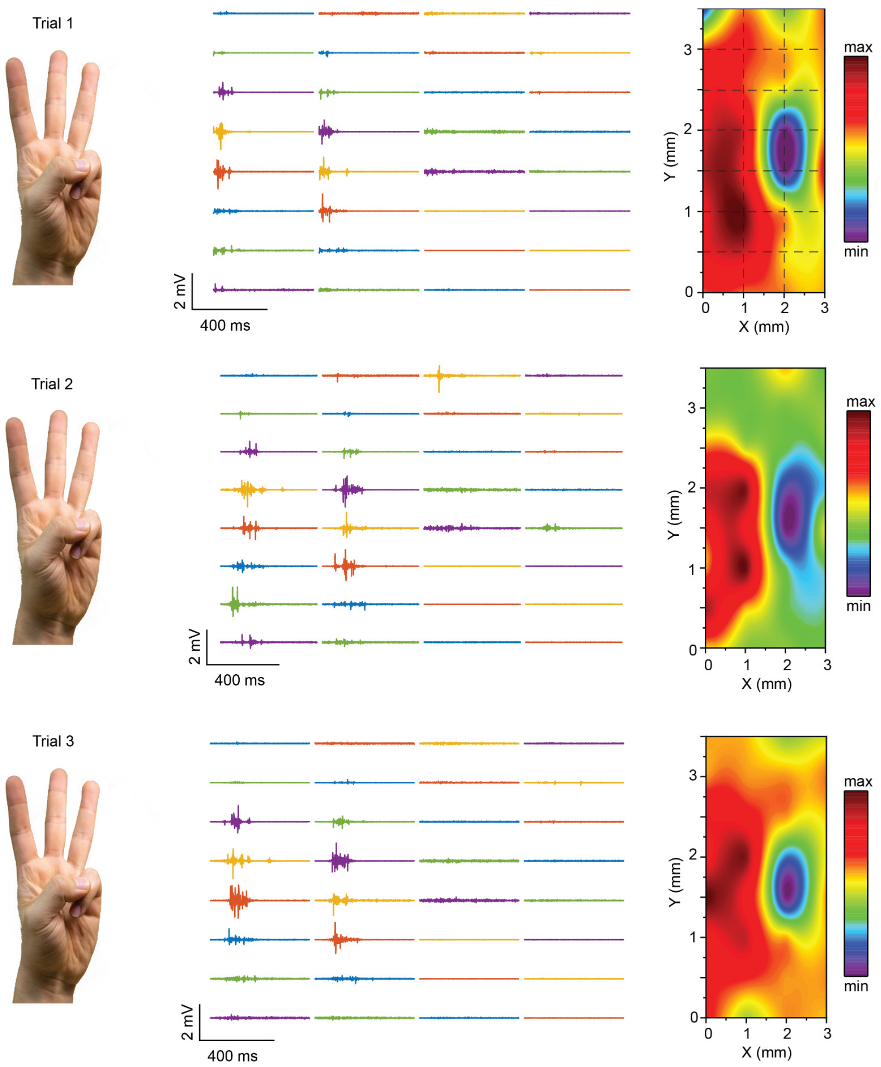

**Fig. S38. Three representative EMG data and calculated heat maps of gesture number three showing the high reproducibility of the stretchable electrode array for sEMG measurement.** During the experiment, the stretchable array was placed at the same muscle location to record the gesture for three separate times. The location of the electrode array is shown in **Fig. 4C**.

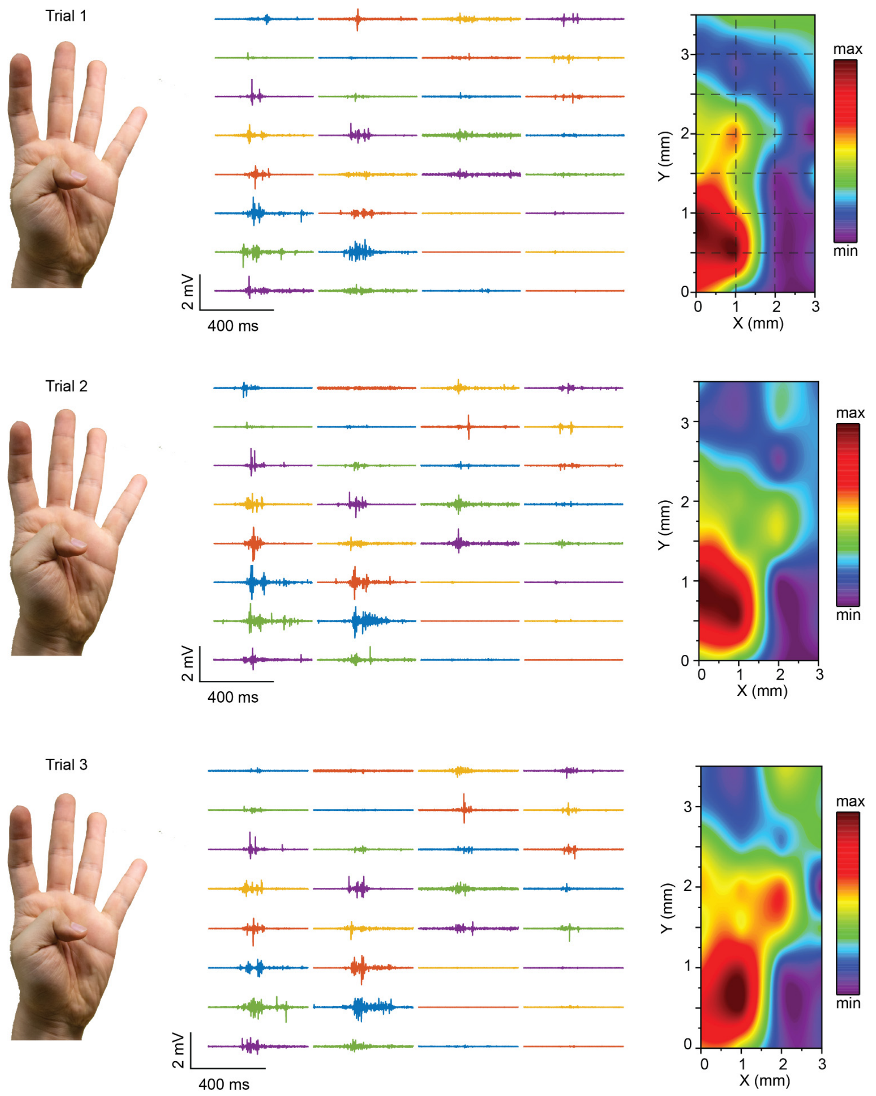

**Fig. S39. Three representative EMG data and calculated heat maps of gesture number four showing the high reproducibility of the stretchable electrode array for sEMG measurement.** During the experiment, the stretchable array was placed at the same muscle location to record the gesture for three separate times. The location of the electrode array is shown in **Fig. 4C**.

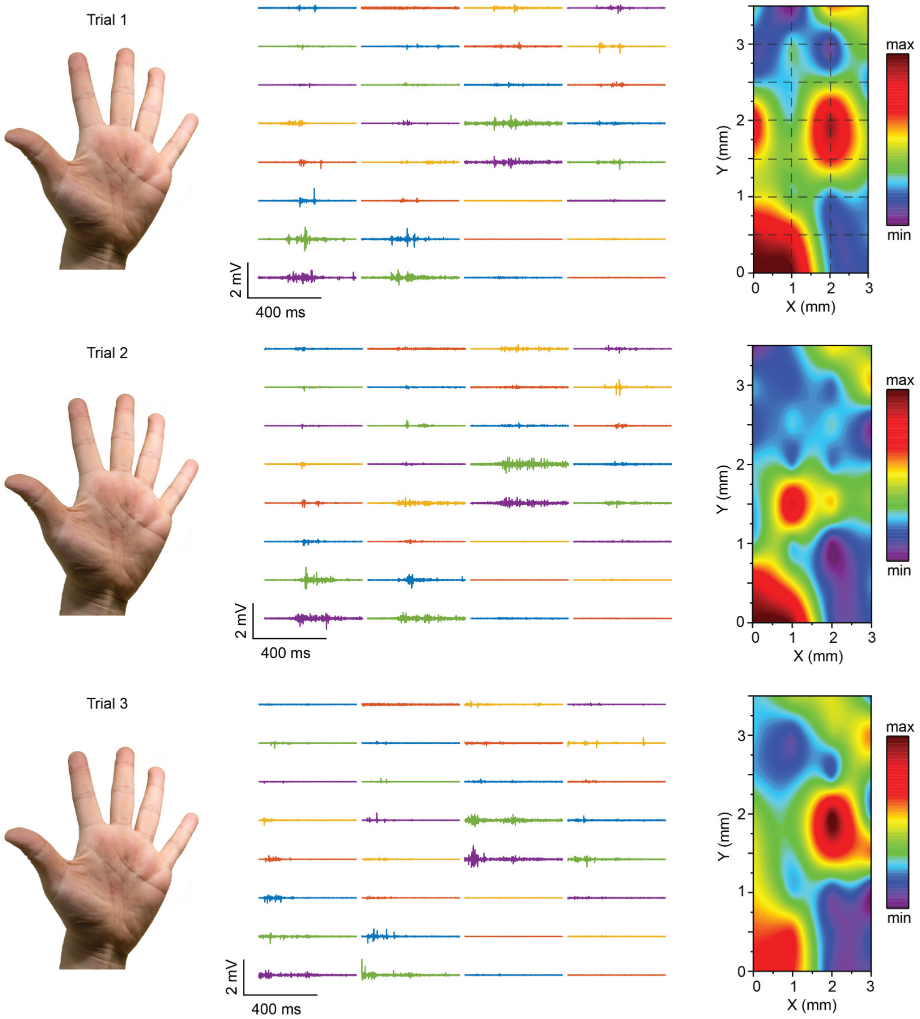

**Fig. S40. Three representative EMG data and calculated heat maps of gesture number five showing the high reproducibility of the stretchable electrode array for sEMG measurement.** During the experiment, the stretchable array was placed at the same muscle location to record the gesture for three separate times. The location of the electrode array is shown in **Fig. 4C**.

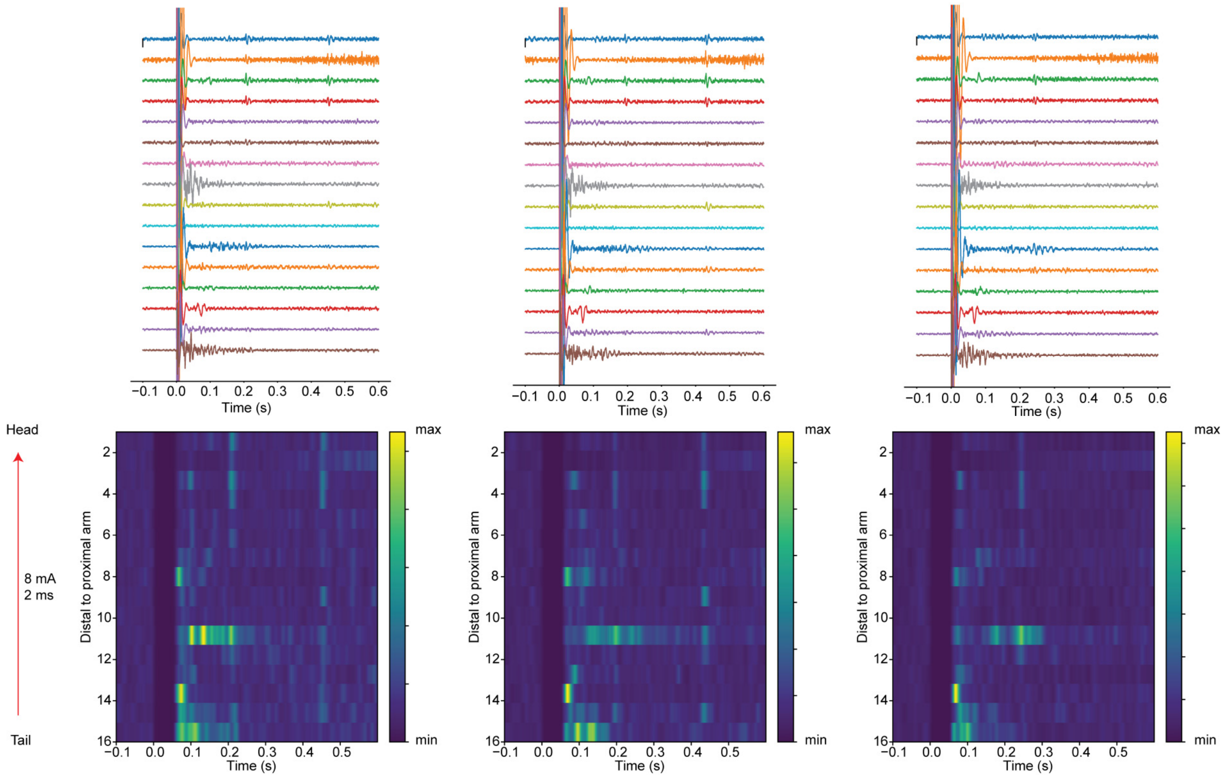

**Fig. S41. Three representative EMG data collected using stretchable electrode array and calculated peristimulus time histogram (PSTH) of octopus arm upon stimulation with the electric field from distal end of the arm to head.** The stretchable probe could move together with the octopus arm and consistently record the same muscle activities.

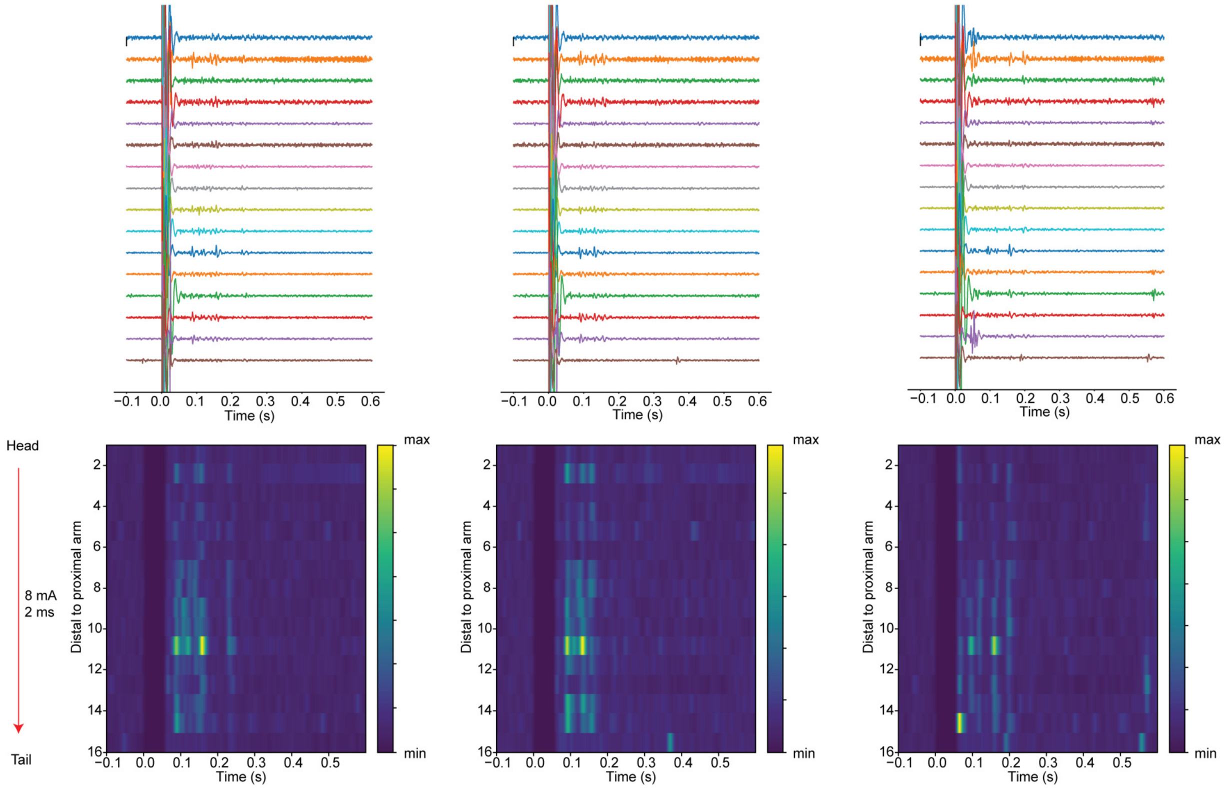

**Fig. S42. Three representative EMG data collected using stretchable electrode array and calculated PSTH of octopus arm upon stimulation with the electric field from head to distal end of the arm.** The stretchable probe could move together with the octopus arm and consistently record the same muscle activities. With the reversed polarity of electrical stimulation, a different yet reproducible pattern of muscle activity could still be captured by the stretchable electrode array, indicating the recorded signals were indeed EMG activities rather than artifacts from octopus arm movements.

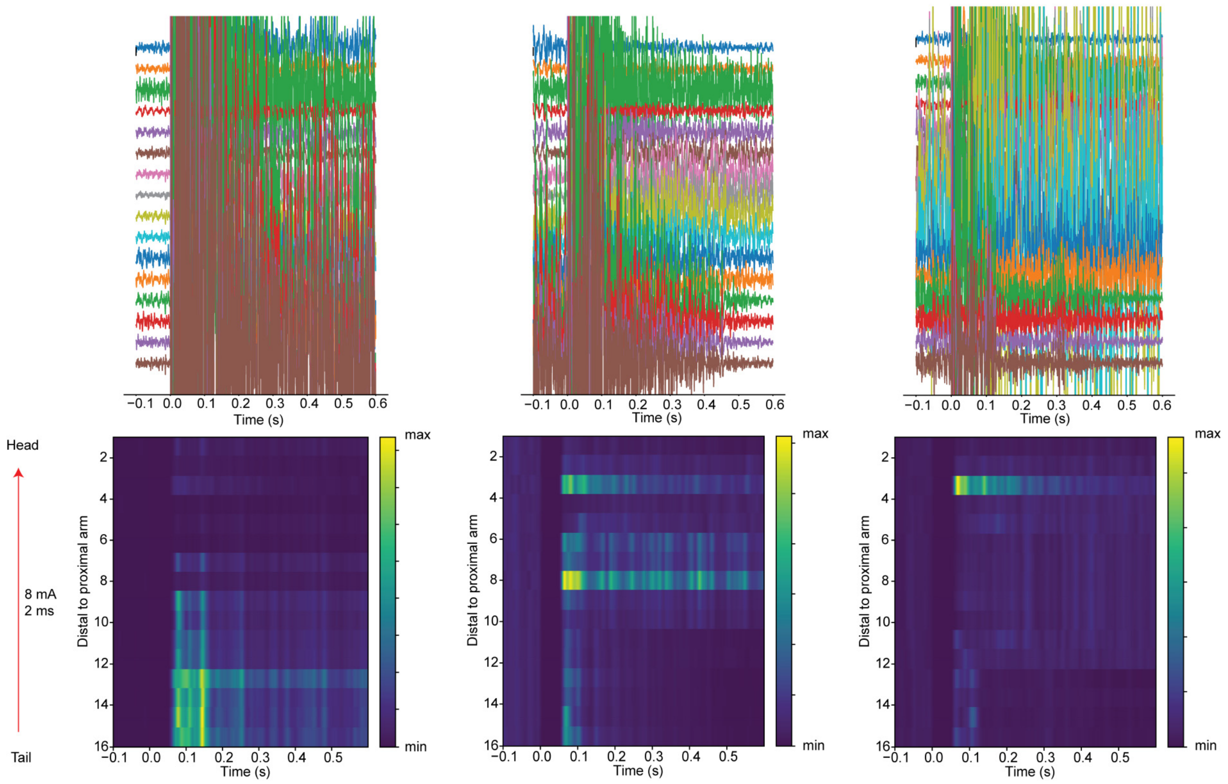

**Fig. S43. Three representative EMG data collected using rigid electrode array and calculated PSTH of octopus arm upon stimulation with the electric field from distal end of the arm to head.** The rigid probe on a polyimide substrate cannot follow the movement of the arm, leading to significant noises during the recording.

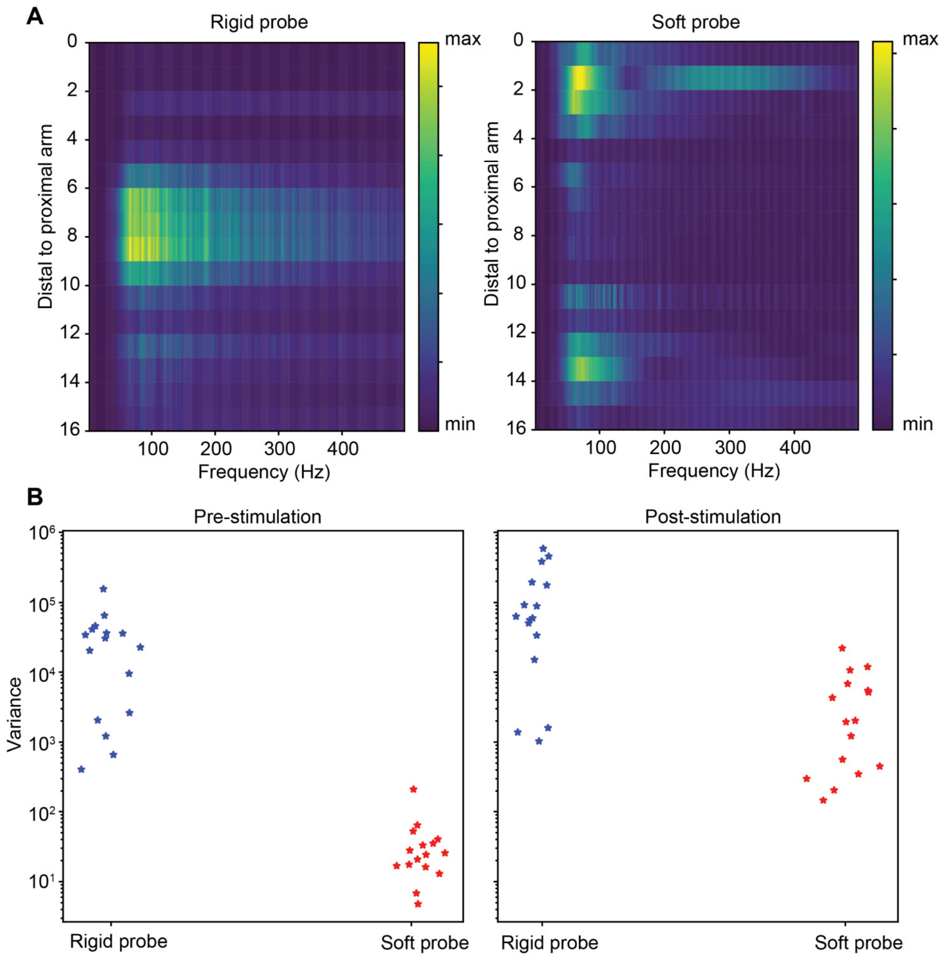

**Fig. S44. Comparison of FFT spectra (A) and variance (B) between soft and rigid probes.** The soft probe has a significantly higher SNR because of its ability to conform during muscle movements.

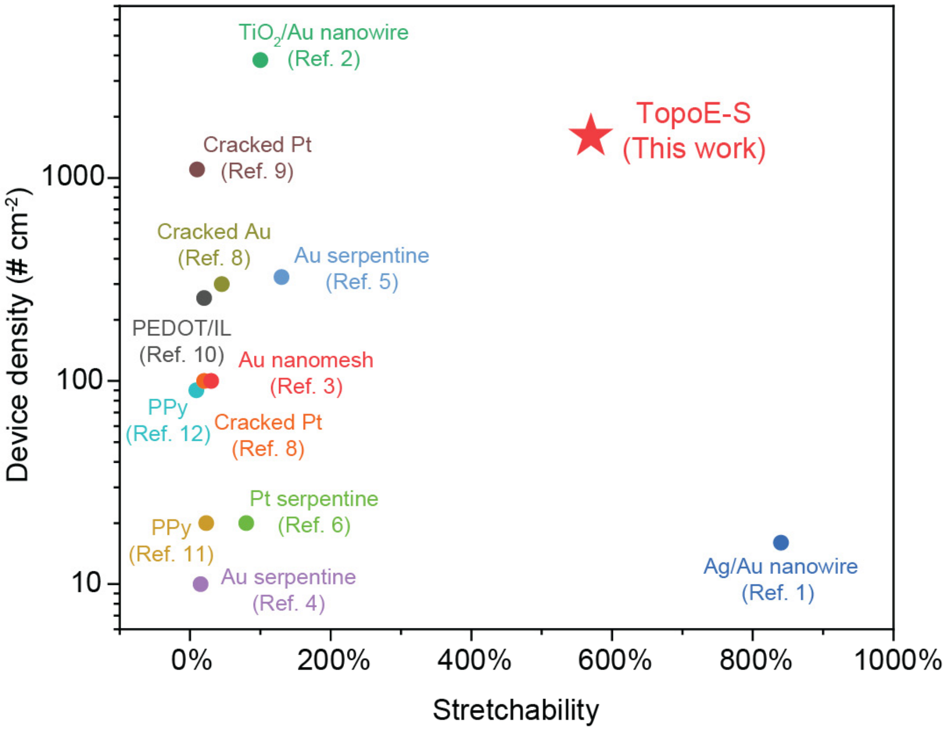

| | Index | Material | Reference |
|---|---|---|---|
| Metal nanomaterials | 1 | Silver/gold nanowire (Ag/Au NW) | *Nat. Nanotechnol.* **13**, 1048–1056 (2018) |
| | 2 | Titanium oxide/gold nanowire ($TiO_2$/Au NW) | *Adv. Mater.* **30**, 1706520 (2018) |
| | 3 | Au nanomesh | *Nat. Nanotechnol.* **14**, 156–160 (2019) |
| Metal microstructures | 4 | Au serpentine | *Nat. Commun.* **5**, 3329 (2014) |
| | 5 | Au serpentine | *Nat. Mater.* **10**, 316–323 (2011) |
| | 6 | Platinum (Pt) serpentine | *Sci. Transl. Med.* **11**, eaax9487 (2019) |
| Microcracked metals | 7 | Cracked Au | *Science* **347**, 159-163 (2015) |
| | 8 | Cracked Pt | *Nat. Biomed. Eng.* **4**, 1010–1022 (2020) |
| | 9 | Cracked Pt | *Adv. Funct. Mater.* **22**, 640–651 (2012) |
| Conducting polymers | 10 | PEDOT:PSS/ionic liquid | *Proc. Natl. Acad. Sci.* **117**, 14769-14778 (2020) |
| | 11 | Polypyrrole (PPy) | *Adv. Mater.* **26**, 1427–1433 (2014) |
| | 12 | Polypyrrole (PPy) | *Adv. Mater.* **29**, 1702800 (2017) |

**Fig. S45. Comparison of device density and stretchability for TopoE-S based stretchable electrode arrays and literature reports.** Rigid inorganic materials can be incorporated into stretchable electronics using special micro- or macro-structure designs. However, these approaches can only realize stretchability at a cost of compromised device density. On the other hand, intrinsically stretchable materials, such as TopoE-S, allow simultaneously high device density and stretchability, which is critical to establish high-resolution biointerfaces with good conformability and minimal invasiveness.

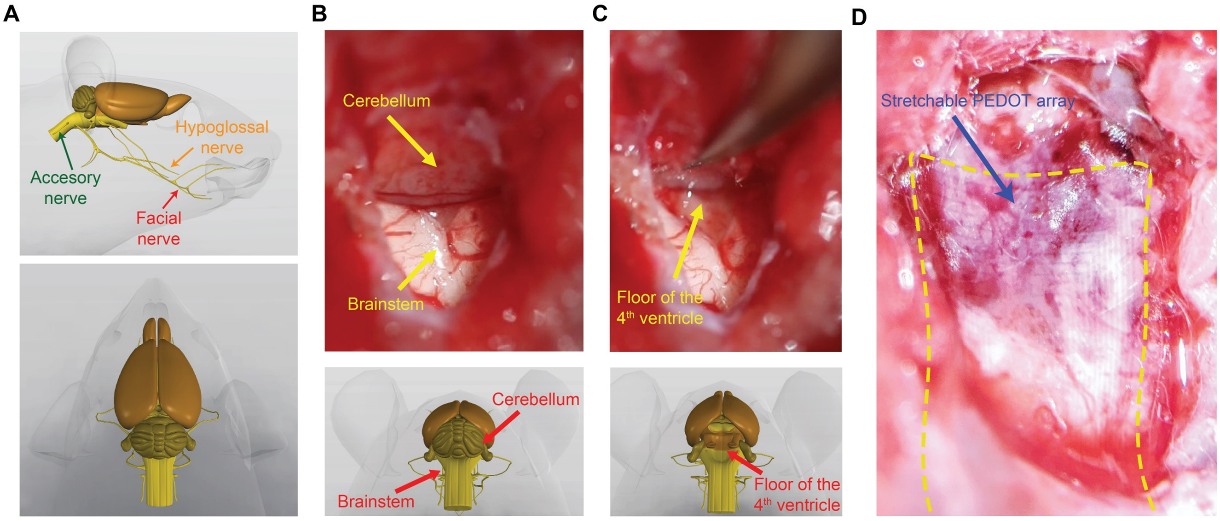

**Fig. S46. Overall design of localized neuromodulation through brainstem using the stretchable electrode array. (A)** Anatomy of three cranial nerves originating from the brainstem that innervate the tongue, whisker, and neck, respectively. **(B-D)** Photographic images showing the surgical procedures to expose the brainstem and a stretchable electrode array conforming to the floor of the $4^{th}$ ventricle.

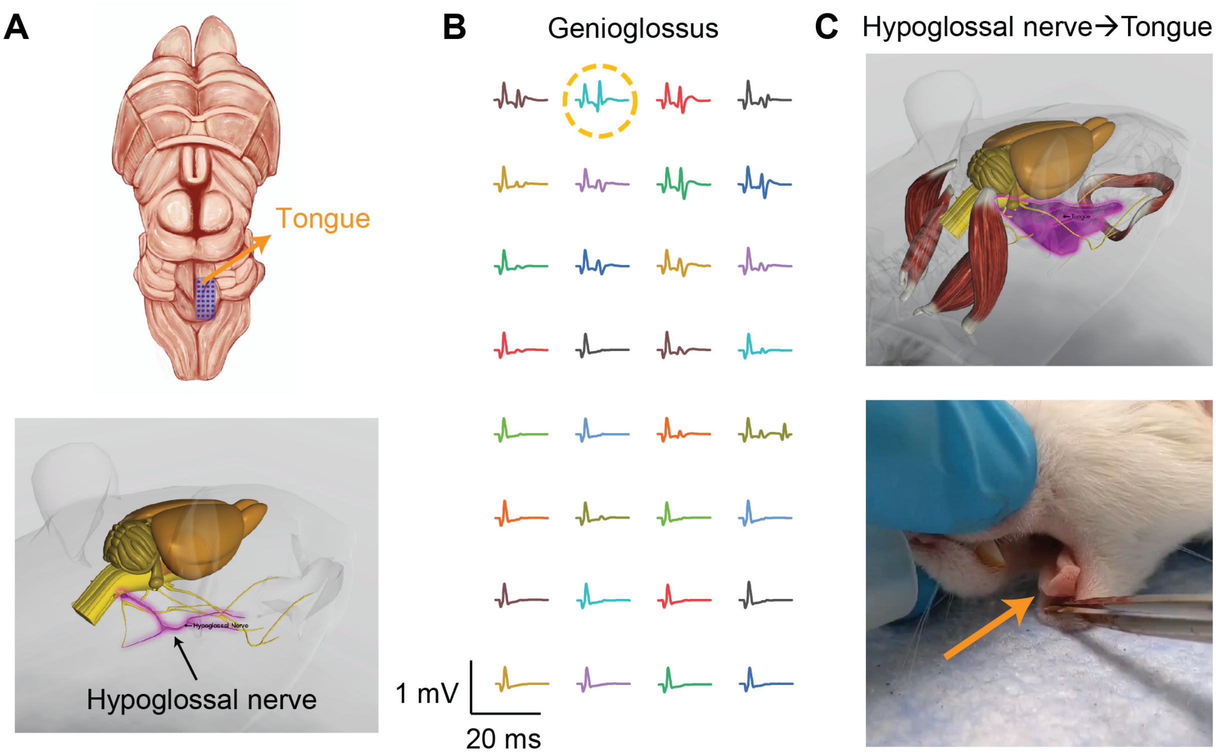

**Fig. S47. Schematic diagrams and recorded data showing the ability of stretchable PEDOT:PSS array for precise control of tongue movements. (A)** Schematic diagrams showing the relative location of the stretchable array on the floor of the fourth ventricle (upper row) and the organization of the right hypoglossal nerve. **(B)** Recorded EMG signals from the tongue showing different responses with respect to the sequential stimulation throughout the entire stretchable array. The orange dashed circle marks the channel with the highest EMG activity of the genioglossus. EMG signals were recorded using a needle electrode inserted into the right half of the tongue. **(C)** Schematic diagram showing the hypoglossal nerve/genioglossus pair (upper row) and a photo (lower row) showing the movement of the right half of the tongue through localized stimulation of the right hypoglossal nucleus.

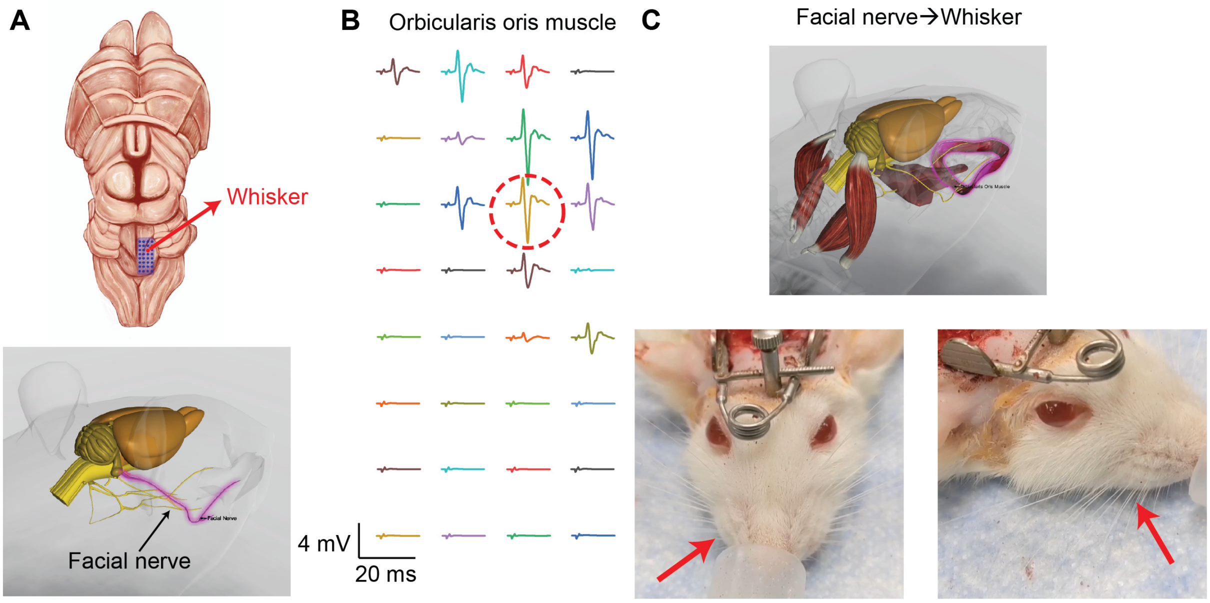

**Fig. S48. Schematic diagrams and recorded data showing the ability of stretchable PEDOT:PSS array for precise control of whisker movements. (A)** Schematic diagrams showing the relative location of the stretchable array on the floor of the fourth ventricle (upper row) and the organization of the right facial nerve. **(B)** Recorded EMG signals from the whisker showing different responses with respect to the sequential stimulation throughout the entire stretchable array. The red dashed circle marks the channel with the highest EMG activity of the orbicularis oris muscle. EMG signals were recorded using a needle electrode inserted into the right half of the orbicularis oris muscle. Schematic diagram showing the facial nerve/orbicularis oris pair (upper row) and two photos showing the movement of the right half of the tongue through localized stimulation of the right facial colliculus.

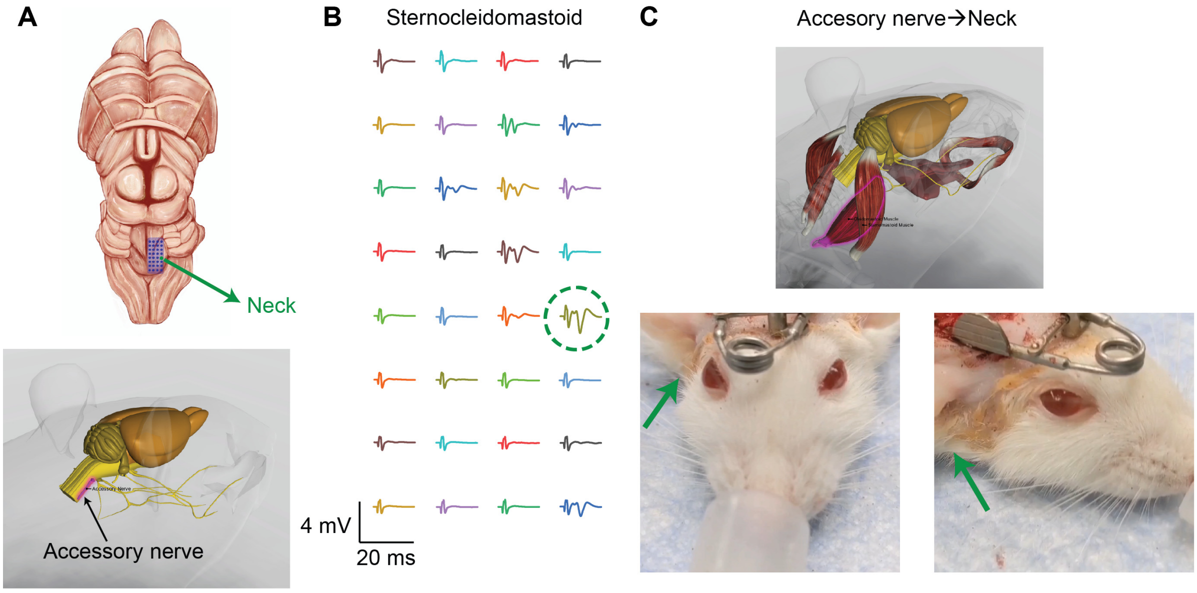

**Fig. S49. Schematic diagrams and recorded data showing the ability of stretchable PEDOT:PSS array for precise control of neck movements through localized stimulation of the accessory nucleus the accessory nerve/sternocleidomastoid pair. (A)** Schematic diagrams showing the relative location of the stretchable array on the floor of the fourth ventricle (upper row) and the organization of the right accessory nerve. **(B)** Recorded EMG signals from the neck showing different responses with respect to the sequential stimulation throughout the entire stretchable array. The green dashed circle marks the channel with the highest EMG activity of the sternocleidomastoid. EMG signals were recorded using a needle electrode inserted into the right half of the sternocleidomastoid. **(C)** Schematic diagram showing the accessory nerve/sternocleidomastoid pair (upper row) and two photos (lower row) showing the movement of the right half of the neck through localized stimulation of the right accessory nucleus.

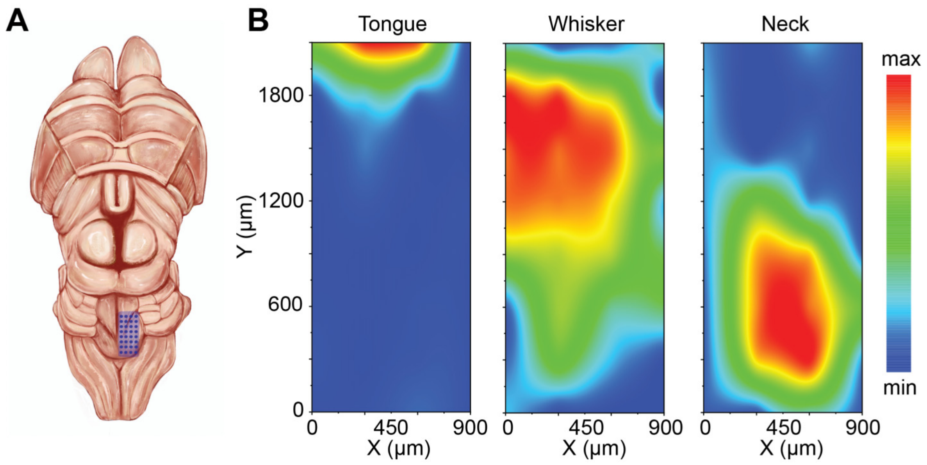

**Fig. S50. The localized neuromodulation can be reproduced on another rat with similar patterns of the activation map.** The result was collected from another rat to show the reproducibility of the functional mapping of the brainstem nucleuses.

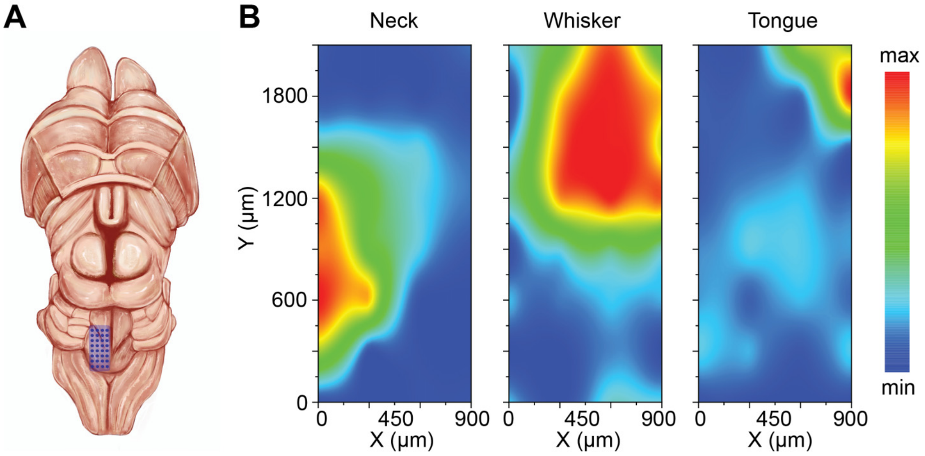

**Fig. S51. Stretchable electrode array placed on the left side of the brainstem could elicit responses of downstream muscles on the same side.**

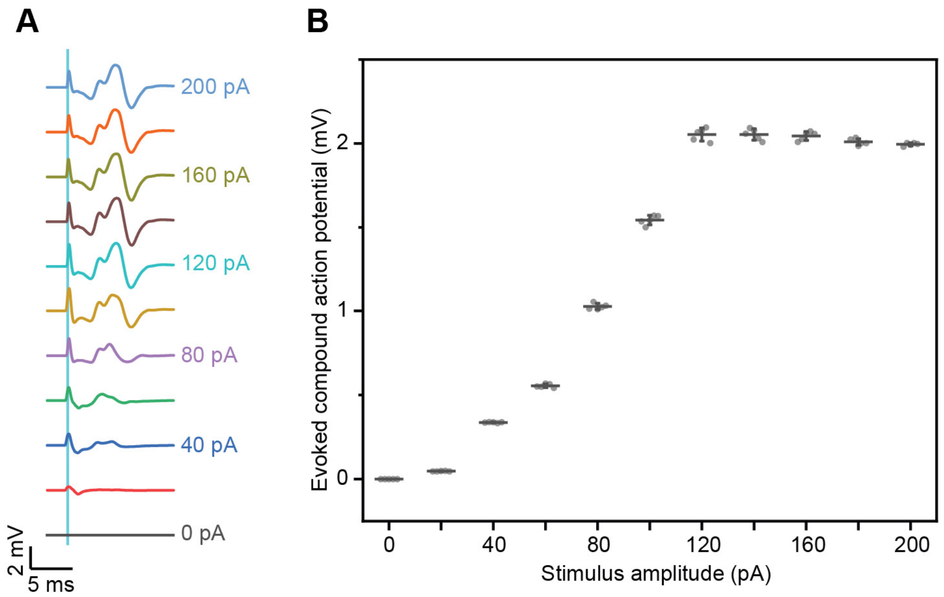

**Fig. S52. The evoked muscle response is dependent on the stimulus amplitude. (A)** Recorded EMG activities with different stimulus current amplitudes from 0 pA to 200 pA for 1 ms. The cyan bar represents the stimulation artifact. During the experiment, current pulses with different amplitudes were applied through one channel of the electrode array while the evoked EMG signals were recorded using a needle electrode inserted in the right half of the orbicularis oris muscle. **(B)** The evoked muscle activity is dependent on the stimulus amplitude, suggesting the potential to fine-tune the amplitudes of muscle movements for even more precise neuromodulation.

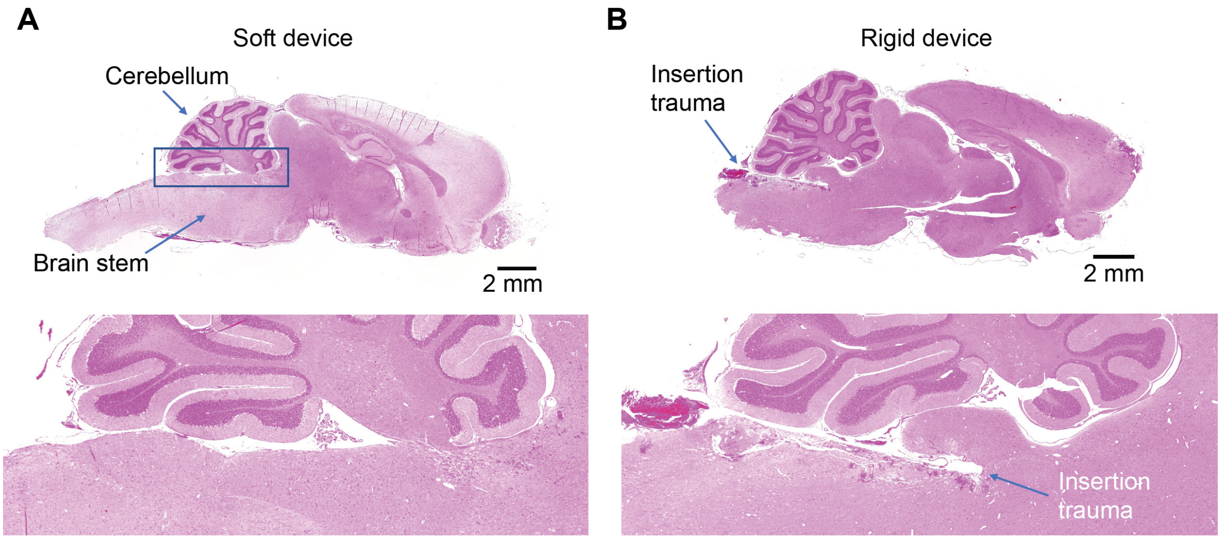

**Fig. S53. H&E staining results of brain slices after insertion of the soft (A) and rigid (B) probes.** The rigid probe would cause severe insertion trauma at the brain stem. The soft probe was prepared on a PDMS substrate while the rigid probe was prepared on a polyimide substrate. In both cases, PEDOT:PSS was used as the electrode material.

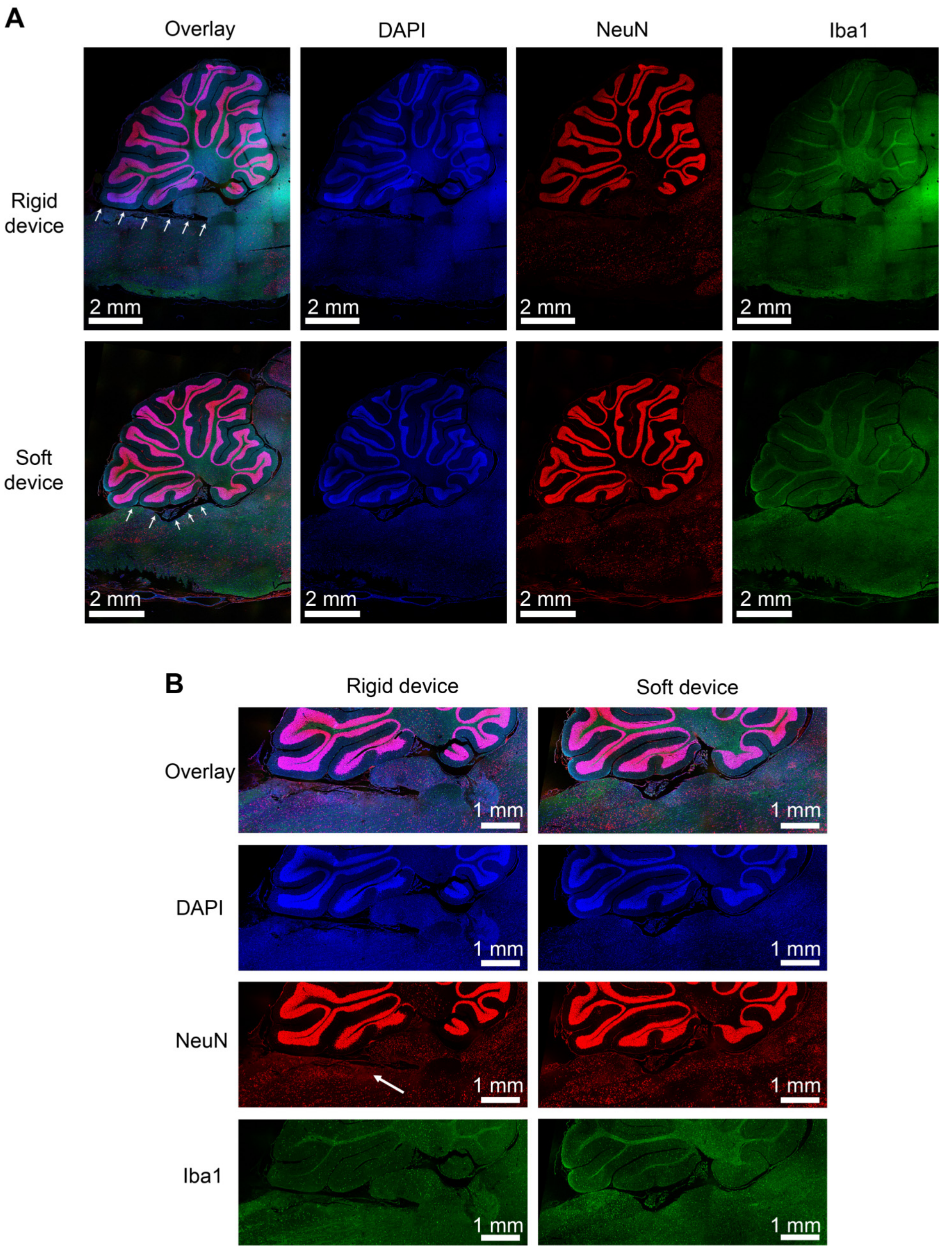

**Fig. S54. Immunofluorescence results of brain slices after insertion of the soft and rigid probes stained with DAPI, NeuN, and Iba1.** The insertion of the rigid probe would cause local death of neurons near the insertion site. Both probes were retracted before tissue sectioning.

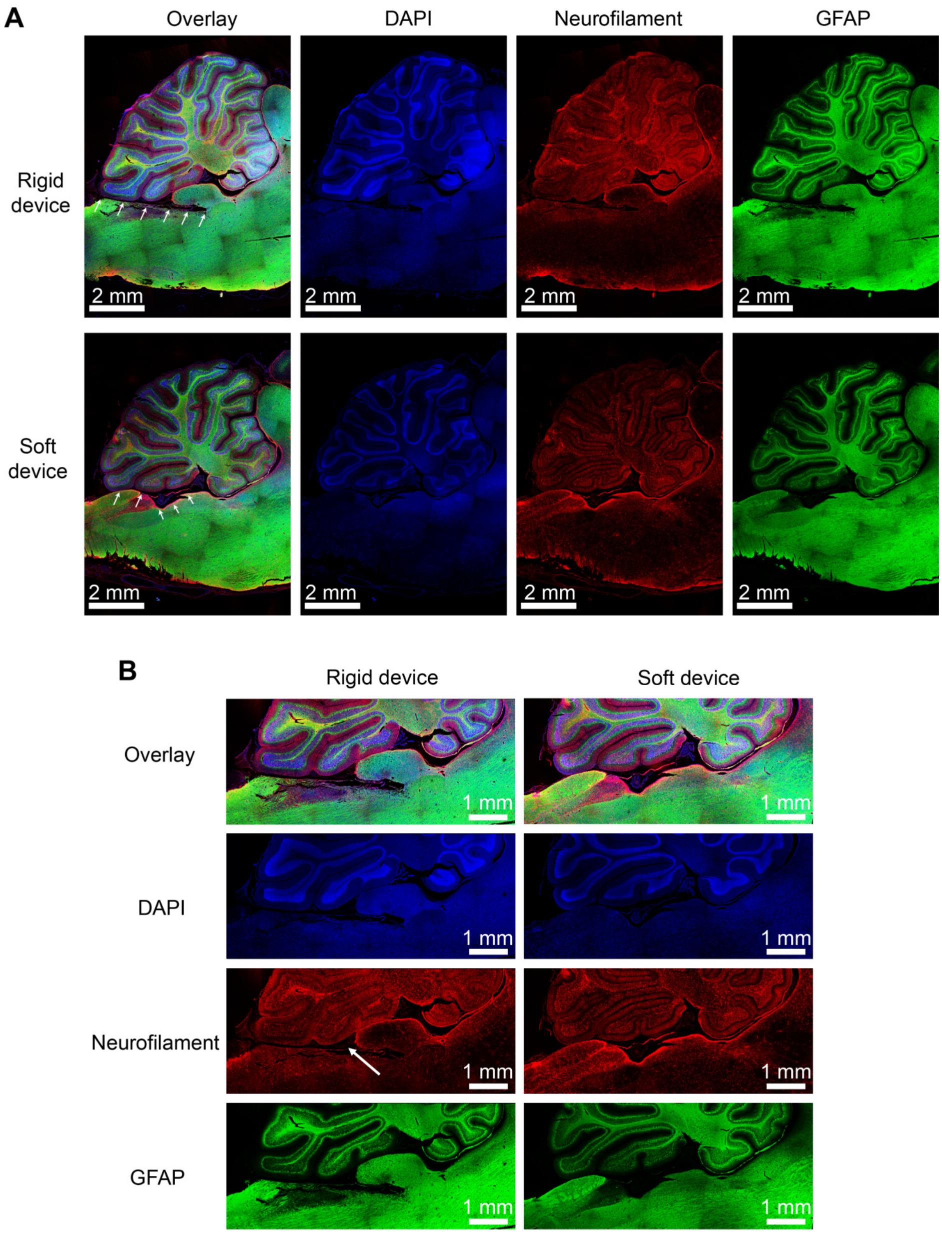

**Fig. S55. Immunofluorescence results of brain slices after insertion of the soft and rigid probes stained with DAPI, Neurofilament, and GFAP.** The insertion of the rigid probe would cause local damage of neurofilament near the insertion site. Both probes were retracted before tissue sectioning.

### References

1 Greczynski, G., Kugler, T. & Salaneck, W. R. Characterization of the PEDOT-PSS system by means of X-ray and ultraviolet photoelectron spectroscopy. *Thin Solid Films* **354**, 129-135, doi:10.1016/S0040-6090(99)00422-8 (1999).

2 Funda, S., Ohki, T., Liu, Q., Hossain, J., Ishimaru, Y., Ueno, K. & Shirai, H. Correlation between the fine structure of spin-coated PEDOT:PSS and the photovoltaic performance of organic/crystalline-silicon heterojunction solar cells. *Journal of Applied Physics* **120**, 033103, doi:10.1063/1.4958845 (2016).

3 Chang, S. H., Chiang, C., Kao, F., Tien, C. & Wu, C. Unraveling the Enhanced Electrical Conductivity of PEDOT:PSS Thin Films for ITO-Free Organic Photovoltaics. *IEEE Photonics Journal* **6**, 1-7, doi:10.1109/JPHOT.2014.2331254 (2014).

4 Alemu, D., Wei, H.-Y., Ho, K.-C. & Chu, C.-W. Highly conductive PEDOT:PSS electrode by simple film treatment with methanol for ITO-free polymer solar cells. *Energy & Environmental Science* **5**, 9662-9671, doi:10.1039/C2EE22595F (2012).

5 Hecht, D. S., Thomas, D., Hu, L., Ladous, C., Lam, T., Park, Y., Irvin, G. & Drzaic, P. Carbon-nanotube film on plastic as transparent electrode for resistive touch screens. *Journal of the Society for Information Display* **17**, 941, doi:10.1889/jsid17.11.941 (2009).

6 Chen, Z., Li, W., Li, R., Zhang, Y., Xu, G. & Cheng, H. Fabrication of Highly Transparent and Conductive Indium–Tin Oxide Thin Films with a High Figure of Merit via Solution Processing. *Langmuir* **29**, 13836-13842, doi:10.1021/la4033282 (2013).

7 Hu, L., Kim, H. S., Lee, J. Y., Peumans, P. & Cui, Y. Scalable coating and properties of transparent, flexible, silver nanowire electrodes. *ACS Nano* **4**, 2955-2963, doi:10.1021/nn1005232 (2010).

8 Kim, N., Kee, S., Lee, S. H., Lee, B. H., Kahng, Y. H., Jo, Y.-R., Kim, B.-J. & Lee, K. Highly Conductive PEDOT:PSS Nanofibrils Induced by Solution-Processed Crystallization. *Advanced Materials* **26**, 2268-2272, doi:10.1002/adma.201304611 (2014).

9 Liu, Y., Liu, J., Chen, S., Lei, T., Kim, Y., Niu, S., Wang, H., Wang, X., Foudeh, A. M., Tok, J. B. & Bao, Z. Soft and elastic hydrogel-based microelectronics for localized low-voltage neuromodulation. *Nat Biomed Eng* **3**, 58-68, doi:10.1038/s41551-018-0335-6 (2019).

10 Matsuhisa, N., Jiang, Y., Liu, Z., Chen, G., Wan, C., Kim, Y., Kang, J., Tran, H., Wu, H.-C., You, I., Bao, Z. & Chen, X. High-Transconductance Stretchable Transistors Achieved by Controlled Gold Microcrack Morphology. *Advanced Electronic Materials* **5**, 1900347, doi:10.1002/aelm.201900347 (2019).
